# Diverse megamammals exploited by humans, chronology and palaeoecology at Taima-Taima, Late Pleistocene, South America

**DOI:** 10.64898/2026.01.22.701027

**Authors:** Jorge D. Carrillo-Briceño, Karina V. Chichkoyan, Arturo Jaimes, Lorena Becerra-Valdivia, Oscar E. Wilson, Kévin Le Verger, Dimila Mothé, Diego Vargas, Andrés E. Reyes-Céspedes, Lautaro Hilbert, Rodolfo Sánchez, Nohé Gilson, Olivier Tombret, Astrid Bauville, Marine Libot, Alfredo A. Carlini, Antoine Zazzo, Marcelo R. Sánchez-Villagra

## Abstract

The arrival of humans to the Americas at the end of the Pleistocene has been one of the foundations of the hypothesis that hunter-gatherer groups had a direct role in the extinction of megafauna. However, empirical evidence of the exploitation of megafauna by humans and associated chronological context are still scarce, especially in South America. Using evidence of modifications on osteological remains, we provide a reassessment of the megafauna that was exploited at the classic Taima-Taima site (Venezuela), with an updated chronology and paleoenvironmental inferences. Our results show that, at least, five species of extinct megaherbivores were probably exploited at the site, expanding the previous records, restricted to the proboscidean *Notiomastodon platensis* and the giant armadillo *Glyptotherium* cf. *G*. *cylindricum*. The new data of the other exploited taxa include the terrestrial sloth *Glossotherium*, the macrauchenid cf. *Xenorhinotherium bahiense* and a toxodontid; both killing and slaughtering activities were likely practised at the site. Seven new enamel radiocarbon dates show an age of 15.3–17.8 cal kyr BP for the fossiliferous strata, although this is likely to be a minimum age given the material’s propensity for modern carbon uptake. The study of phytoliths and starch grains from the dental calculus of *N. platensis* identified five plant taxa from the phytoliths: arboreal dicotyledons, Poaceae, Marantaceae, Asteraceae and Arecaceae. Ecometrics studies of dental diversity reveal how no other single site on the continent preserves such a diversity of grazing and browsing megaherbivores exploited by humans.

## 1. Introduction

The American continent witnessed one of the major megafauna (body mass > 44 kg) extinctions at the Pleistocene–Holocene transition (Barnosky et al., 2004), and although it is currently understood as a multicausal event (Cione et al., 2003; Nogués-Bravo et al., 2010; Politis et al., 2009; Barnosky and Lindsey, 2010; Broughton and Weitzel, 2018; MacPhee, 2018; Fariña and Vizcaíno, 2024), the topic remains an ongoing and highly controversial debate (Stewart et al., 2025). The arrival of the first humans at the end of the Pleistocene (Dillehay et al., 1992; Politis et al., 2009; Fariña et al., 2014; Ardelean et al., 2020; Bennett, 2021; Pansani et al., 2023; Del Papa et al., 2024), has been correlated with this extinction, forming the basis for the overhunting hypothesis (Martin, 1967, 1984; Martin and Steadman, 1999). However, archaeological evidence for megafaunal killing/scavenging by humans is limited, although sites with evidence of these activities have been reported for the entire American continent (Borrero, 2008; Grayson and Meltzer, 2003; Bampi et al., 2022; Del Papa et al., 2024; Fariña and Vizcaíno, 2024, Solís-Torres et al., 2025; and references therein). Recent data with a chronological and quantitative faunal framework (Prates and Perez, 2021; Prates et al., 2025) suggest that extinct megafauna (especially in regions with high abundance and diversity) were the main prey of early hunter-gatherer groups in southern South America from ∼13,000 to 11,600 cal yr BP.

Located at the northern gateway to South America, sites in Colombia and Venezuela are critical for tracing the arrival of humans to the continent during the Late Pleistocene, their interaction with local fauna, and their subsequent demographic and migration dynamics (Cooke, 2021). One of the prime sites in this region is Taima-Taima, located in the coastal area of Falcón State, northwestern Venezuela (Fig. 1), which preserves some of the best-known faunal assemblages from Late Pleistocene in northern South America (Carrillo-Briceño, 2015; Reyes-Céspedes et al., 2023)—including giant land tortoises and mammals of different body sizes, (electronic supplementary material S1). Here, projectile point technology and other lithic artifacts have been found in close, physical association with diverse megafaunal remains (Cruxent, 1970, 1978, 1979; Ochsenius and Gruhn, 1979; Oliver and Alexander, 2003). This includes the discovery of two El Jobo projectile points (Jaimes et al., 2024a; Vargas et al., 2025) embedded in the pelvis of two individuals of the gomphothere *Notiomastodon platensis* (Bryan et al., 1978; Cruxent, 1979; Ochsenius and Gruhn, 1979; see electronic supplementary material S1). As such, since the 1970s, the unequivocal evidence of human-megafauna interaction at Taima-Taima has been particularly important in First Americans research. However, despite its historical and empirical importance, the site had not been re-examined until recently.

**Figure 1.**
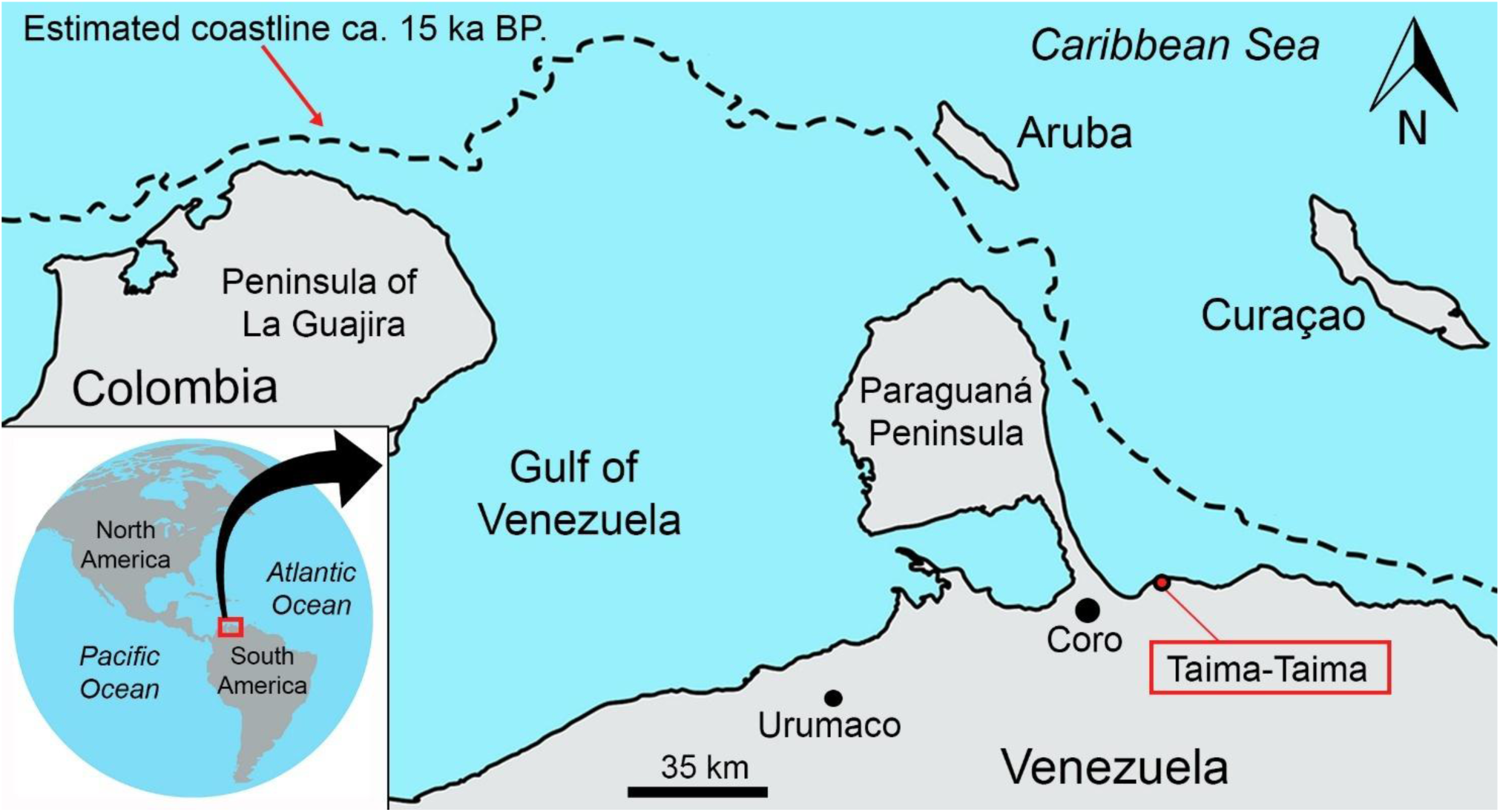
Geographic location of the Taima-Taima site. Estimated coastal line around 15 ka yr BP is based approximately on the 100-meter bathymetric line (Peltier and Fairbanks, 2006).

In 2022, Carlini et al. reported on glyptodont skulls from Taima-Taima, identified as *Glyptotherium* cf. *G*. *cylindricum*, breakage patterns interpreted as intentional percussion blows associated with hunting. Two inverted carapaces were also reported (electronic supplementary material S1), with the most complete recovered during the first excavation campaigns (Cruxent, 1967, 1978) and only preserving the articulated pelvis inside (currently in the Museo De Ciencias Naturales de Caracas, disarticulated and fragmented). The second carapace was discovered during the excavation of 1976, preserving part of the pelvis (ilium) and some vertebrae (Bryan, 1979). Both carapaces were found *in situ* in an upside-down position, probably placed this way intentionally by humans for processing (Bryan, 1979).

Here, we provide a reassessment of the diversity and temporal distribution of megaherbivores present in Taima-Taima and identify evidence of probable human-made modifications in osteological remains (see also electronic supplementary material S1 and S2). Results show that this ancient watering hole was likely a strategic location for subsistence exploitation of multiple megaherbivore species, offering new insights into human-megafauna dynamics during the Late Pleistocene in a key region within South America.

## 2. Materials and methods

### 2.1 Taxonomy and bone surface modifications

We investigated the collections housed in the Centro de Investigaciones Antropológicas, Arqueológicas y Paleontológicas (CIAAP) of the Universidad Experimental Francisco de Miranda (UNEFM), and Museo Comunitario de Taratara Cristóbal Higuera (MCH-Pv-) in Falcón State, Laboratorio de Arqueología del Instituto Venezolano de Investigaciones Científicas (IVIC) in Miranda State, and the Museo de Ciencias Naturales de Caracas (MCNC), all in Venezuela.

We report on extinct megaherbivore bones with evidence of probable pre-burial modifications caused by humans, represented by a sample of 11 bone elements (Table 1). The taxonomic assignment of the faunal assemblage is documented in the electronic supplementary material S3, and their stratigraphic context in the electronic supplementary material S2. For each bone, only the surface modifications of probable anthropic origin were macroscopically identified and analyzed at the microscopic level with a low magnification stereoscopic lens (Wild M8×40). Photos were taken using a Nikon camera with lenses AF-S Macro Nikkor 60 mm and AF-S Nikkor 18-200 mm. Bone modifications were characterised through classic morphological attributes shown in Table 1, which are based on the most conservative criteria (e.g., Lyman, 1987, 1994; Bello and Soligo, 2008; Domínguez-Rodrigo et al., 2009; Fernández-Jalvo and Andrews, 2016, among others; see electronic supplementary material S4). The latter allows us to support our hypothesis that the bone modifications presented here from the Taima-Taima site are likely of anthropic nature and not from non-human agents, such as those produced by predators/scavengers or trampling (electronic supplementary material S4). The modifications identified in the bones studied here and presented in Table 1 are described in detail in the electronic supplementary material S4.

**Table 1.** Fossil specimens of megaherbivores from Taima-Taima with surface modifications. Specimens belong to the same individual, referred to here as the 1976 skeleton (*)

| Systematics | Catalog N° | Stratigraphical unit | Element/s | Chop mark | Slicing mark | Cut mark | Hertzian cones | Internal microsteps | Internal micro-strations | Shoulder effect | Scraping marks | Percusion marks |
| --- | --- | --- | --- | --- | --- | --- | --- | --- | --- | --- | --- | --- |
| XENARTHRA (Pilosa) |  |  |  |  |  |  |  |  |  |  |  |  |
| *Glossotherium* cf. *robustum* | IVIC-AP-035 | Us | Left humerus | X | ? |  | X |  | X |  |  |  |
| Indet. | MCNC-Pal-1831 | ?Ms | Phalange II | X |  |  | X | ? | X |  |  |  |
| †LITOPTERNA |  |  |  |  |  |  |  |  |  |  |  |  |
| cf. *Xenorhinotherium bahiense* | MCNC-Pal-1834 | ?Ms | Right fused ulna-radius |  |  | X |  |  |  | X |  |  |
| †NOTOUNGULATA |  |  |  |  |  |  |  |  |  |  |  |  |
| Indet. (probably *Mixotoxodon*) | CIAAP-1533 | Us | Left ulna |  | X | X | ? |  |  |  |  |  |
| PROBOSCIDEA |  |  |  |  |  |  |  |  |  |  |  |  |
| *Notiomastodon platensis* | CIAAP-46 | Ms | *Left humerus | X |  | X |  |  |  |  |  |  |
| *Notiomastodon platensis* | CIAAP-47-229 | Ms | *Left scapula | X | X |  |  |  |  |  |  |  |
| *Notiomastodon platensis* | CIAAP-111 | Ms | *Right femur |  |  | X | ? |  |  |  |  |  |
| *Notiomastodon platensis* | CIAAP-109-211 | Ms | *Right tibia |  | X | X |  |  |  |  |  |  |
| *Notiomastodon platensis* | CIAAP-119-110-210 | Ms | Right scapula |  | X | X |  |  |  |  |  |  |
| *Notiomastodon platensis* | CIAAP-19-233 | Ms | Left ilium fragment | X | X | X | X |  |  |  | X | X |
| *Notiomastodon platensis* | CIAAP-213-230 | ?Bs | Right scapula | ? | X | X |  |  |  |  |  | X |

### 2.2 Radiocarbon dating

#### 2.2.1 Bone/dentine collagen dating

Twenty-nine samples of megaherbivores from Taima-Taima (electronic supplementary material S1) were sent to the Oxford Radiocarbon Accelerator Unit (ORAU; UK) for radiocarbon dating of bone/dentine collagen. The dated material comes from different individuals of *N*. *platensis*, *Eremotherium*, *Glyptotherium*, and other indeterminate mammals collected in different excavation campaigns in Taima-Taima (see Table S1.2 in electronic supplementary material S1), and housed in national collections of the CIAAP, IVIC, and MCNC. All the samples were collected and transported for study with authorization from the Instituto del Patrimonio Cultural de Venezuela (IPC) using permissions: VE-IPC-CEBC-PP-06/2022-1 and VE-IPC-CEBC-PP-01/2023.

The collagen preservation screening was carried out prior to radiocarbon dating and to assessing collagen preservation, and the 29 bone/dentine samples were pre-screened for nitrogen content. This uses %N as a proxy for collagen abundance since nitrogen is derived solely from the proteinaceous component, the majority of which being collagen (Sillen and Parkington, 1996). If the N% of the sample is >0.76, then it is likely that the bone will yield enough collagen for radiocarbon dating (Brock et al., 2012). Pre-screening involved measuring the %N of 2 mg of whole bone powder (placed into a tin capsule) in a CHN elemental analyser (Carlo-Erba NA-2000) coupled to a continuous flow isotope ratio mass spectrometer (Sercon 20/20) (Brock et al., 2010a, b; Brock et al., 2012). Samples were run alongside an internal standard, L-Alanine (Sigma-Aldrich). For the radiocarbon dating, bones were first manually cleaned by air abrasion and cut using a small diamond disc. Fragments were chemically pretreated and targeted for “collagen” (following the use of this term in (DeNiro and Weiner, 1988; Hedges and van Klinken, 1992; van Klinken, 1999) by means of an acid-base-acid protocol, gelatinization and Ezee-filtration (ORAU pretreatment code, AG), (Brock et al., 2010a, b). Ultrafiltration was omitted given the likely poor collagen preservation at the site.

#### 2.2.2 Tooth enamel dating

Tooth enamel dating was also performed on seven fossil teeth from Taima-Taima at the radiocarbon laboratory of the Muséum national d’Histoire naturelle (MNHN), Paris and AMS 14C dating was performed using the AMS ECHoMicadas system at LSCE (Saclay). The dated tooth enamel comes from molars of five individuals of *N*. *platensis* and a molar and an incisor of *Equus* sp., with their stratigraphic origin shown in Table 2 and Fig. 2. The enamel surface of the specimens was cleaned of sediments, and the dentine was removed with a Dremel to isolate the enamel from the rest of the dental tissue. The enamel (∼2 g) was then grounded using a steel mortar and pestle, followed by grinding in an agate mortar to a particle size of < 100 microns. The powder was then finely (5-10 microns) crushed using a McCrone Microniser Retsch following (Wood et al., 2016) as this approach increases the efficiency of the acetic acid pre-treatment aimed at removing diagenetic carbonates and therefore improves the radiocarbon ages obtained. The resulting powder was pre-treated under a light vacuum for 20 hours with a solution of 1 N acetic acid then rinsed with milliQ water and oven-dried at 50 °C. About 250 mg of powder was then reacted under vacuum with orthophosphoric acid at 70 °C for around 20 min. The CO2 gas produced was then reduced in the presence of hydrogen and iron to produce graphite. Samples were then pressed into targets and 14C ages were measured on the compact AMS ECHoMicadas at LSCE, Saclay.

**Figure 2.**
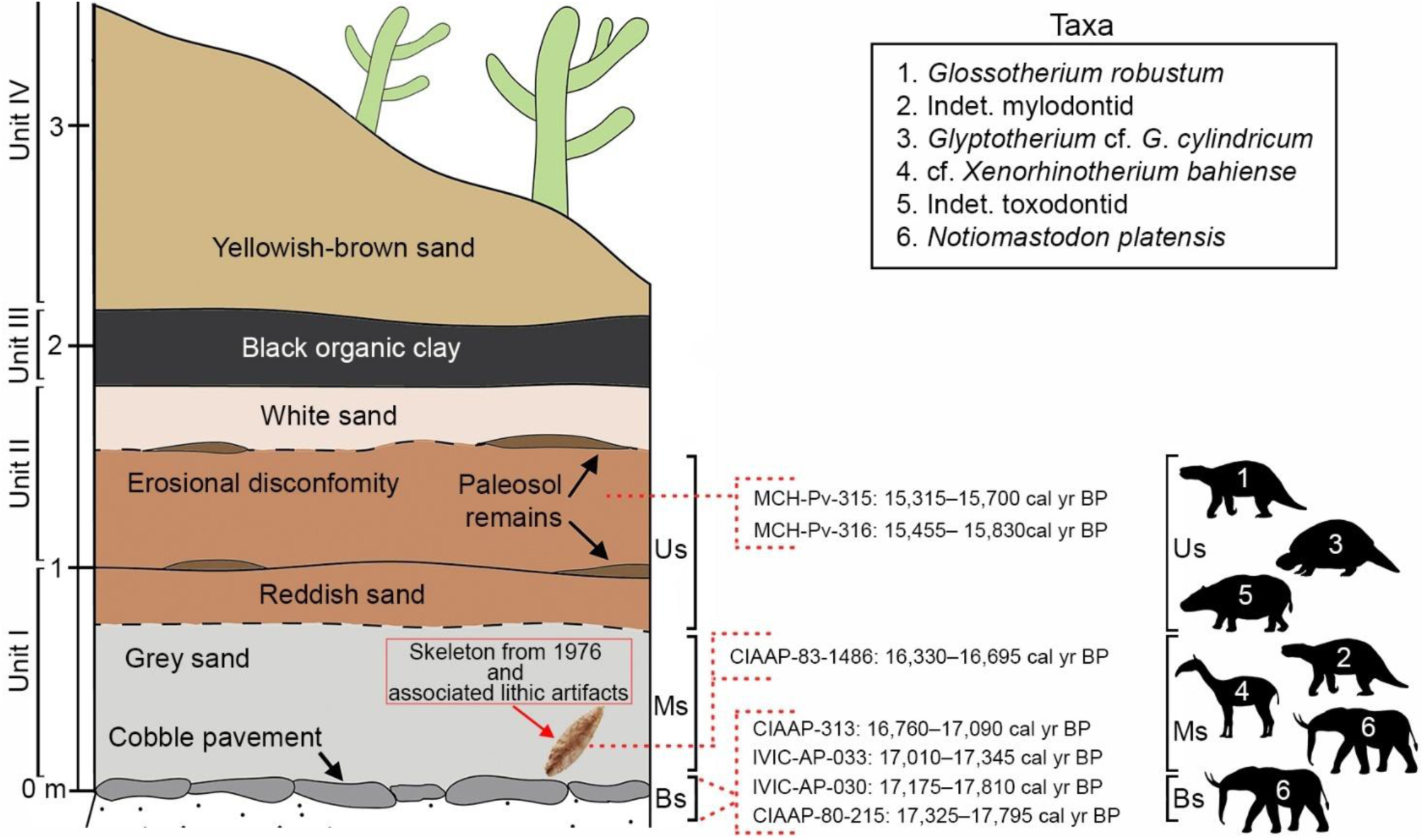
Schematic profile of the Taima-Taima site showing, Basal (Bs), Medium (Ms), and Upper (Us) fossiliferous strata, new dating (see Table 2), taxa reported here with bones preserving surface modifications, lithic artifacts, and their stratigraphic provenance. Profile modified after Carlini et al. (2022). The stratigraphic provenance of the taxa reported here is based on Casamiquela (1979), Carlini et al. (2022), historical archives of JM Cruxent, and authors personal observations. Abbreviations: calibrated years before present (ca yr BP).

**Table 2.** Chronometric data for the radiocarbon dated tooth enamel samples from Taima-Taima. Abbreviations: cal yr BP, calibrated years before Present [calibration using the IntCal20 curve (Reimer et al., 2020)]; Upper stratum (Us), Middle stratum (Ms), and Basal stratum (Bs).

| Sample /catalog<br>N° | Stratum | Taxonomy | Anatomical<br>element | Laboratory<br>N° | Date<br>yr (BP) | ± | cal yr BP<br>(95.4% CI) |
| --- | --- | --- | --- | --- | --- | --- | --- |
| MCH-Pv-315 | Us | *Equus* sp. | Incisor | ECHo6429 | 12,975 | 50 | 15,315–15,700 |
| MCH-Pv-316 | Us | *Equus* sp. | Lower molar | ECHo6430 | 13,060 | 50 | 15,455–15,830 |
| CIAAP-83-1486 | Ms | *Notiomastodon platensis* | Lower right m2 | ECHo7502 | 13,665 | 45 | 16,330–16,695 |
| CIAAP-313 | Bs | *Notiomastodon platensis* | Lower left m3 | ECHo7504 | 13,950 | 45 | 16,760–17,090 |
| IVIC-AP-033 | Ms/Bs | *Notiomastodon platensis* | Left M2 | ECHo6432 | 14,085 | 55 | 17,010–17,345 |
| IVIC-AP-030 | Ms/Bs | *Notiomastodon platensis* | Unerupted m3 | ECHo6431 | 14,330 | 60 | 17,175–17,810 |
| CIAAP-80-215 | Bs | *Notiomastodon platensis* | Lower left m2 | ECHo7503 | 14,365 | 50 | 17,325–17,795 |

#### 2.2.3 Radiocarbon calibrations

The calibration was undertaken using IntCal20 (Reimer et al., 2020) in OxCal 4.4 (Bronk Ramsey, 2009a). All calibrated estimates are noted at 94.5 % credible/confidence intervals.

### 2.3 *Notiomastodon platensis* tooth calculus analysis

Eight dental calculus samples were extracted from *Notiomastodon* specimens from Taima-Taima for phytolith and starch grain analysis, following an adjusted version of the protocol of Santiago-Marrero & Pagán-Jiménez (2023). We adjusted this procedure to exclude the addition of hydrogen peroxide, to prevent potential damage to or digestion of starch grains (electronic supplementary material S6). Following decalcification, slides were prepared for each sample. Phytoliths and starch grains were identified, counted, and photographed with a Leica DM500 light microscope at 200× and 400× magnification, following published reference material (e.g., Twiss et al., 1969; Brown, 1984; Piperno and Pearsall, 1998; Torrence et al., 2003; Ezell et al., 2006; Piperno, 2006; Iriarte and Paz, 2009; Gismondi et al., 2018; Pearsall, 2018). Results from the phytoliths and starch grains were qualitatively compared to published data on diet mesowear in *N*. *platensis* (Wilson et al., 2024) and other herbivores from the region (electronic supplementary material S7).

## 3. Archaeological context

Taima-Taima (11° 29′ 57′′ N, 69° 31′ 20′′ W; ∼ 60 m above sea level) is located about 18 km northeast of the city of Santa Ana de Coro (Fig. 1), near the town of Taratara, Colina municipality, Falcón State. The site is surrounded by hills, in a sedimentary environment influenced by the activity of resurgent springs (Cruxent, 1970; Ochsenius and Gruhn, 1979; Ochsenius, 1980; electronic supplementary material S1) with a continuous water supply that drains to the present coastline, located at approximately 550 m.

During the excavation season of 1976, four stratigraphic units, with a total thickness of almost four meters were described (Bryan, 1979). Casamiquela (1979: 59 pp.) categorised the fossiliferous assemblage from Taima-Taima according to its stratigraphic origin in three strata, using this term in a more temporal than geological (sedimentological) sense, with awareness of eventual differences in the time elapsed between one stratum and the other. These strata (Fig. 2) are the Basal stratum characterized by a cobble pavement of rocks of Miocene origin, covered by the Medium and Upper strata, these last two being characterized by more sandy sediments. These three strata correspond to units I and II described by Bryan (1979). According to Casamiquela (1979), the Unit I/II erosional disconformity (Fig. 2) represents the last evidence of megafauna in the Taima-Taima site. A series of radiocarbon dates were previously published for Taima-Taima (Tamers, 1971; Bryan et al., 1978; Bryan and Gruhn, 1979; electronic supplementary material S5). These range from 9,650±110 yr BP (or 10,975 cal yr BP) from the upper black organic clay (Unit III) to 14,200±300 yr BP (or 17,310-17,305 cal yr BP) from the Basal/Medium strata (Unit I). The four lithic artifacts (two El Jobo projectile points and two scrapers; see electronic supplementary material S1) found *in situ* and in close, physical association with remains of *N*. *platensis*, come from the same layers of the Medium stratum (Cruxent, 1978, 1979; Ochsenius and Gruhn, 1979) (Fig. 2).

## 4. Results

### 4.1 Surface modification of megaherbivore bones and other evidence of anthropic interactions

At least eleven fossil bones from the remains of at least five extinct megaherbivores (e.g., Figs. 3–5) exhibit signs of probable human intervention. The marks are classified in nine kinds (Table 1), and resemble those reported from historical collections (e.g., Chichkoyan et al., 2017; Chichkoyan, 2019; Toledo, 2023) and other sites (e.g., Fariña et al., 2014; Labarca et al., 2020, 2024; Nami et al., 2023; Del Papa et al., 2024) from the Late Pleistocene of South America.

**Figure 3.**
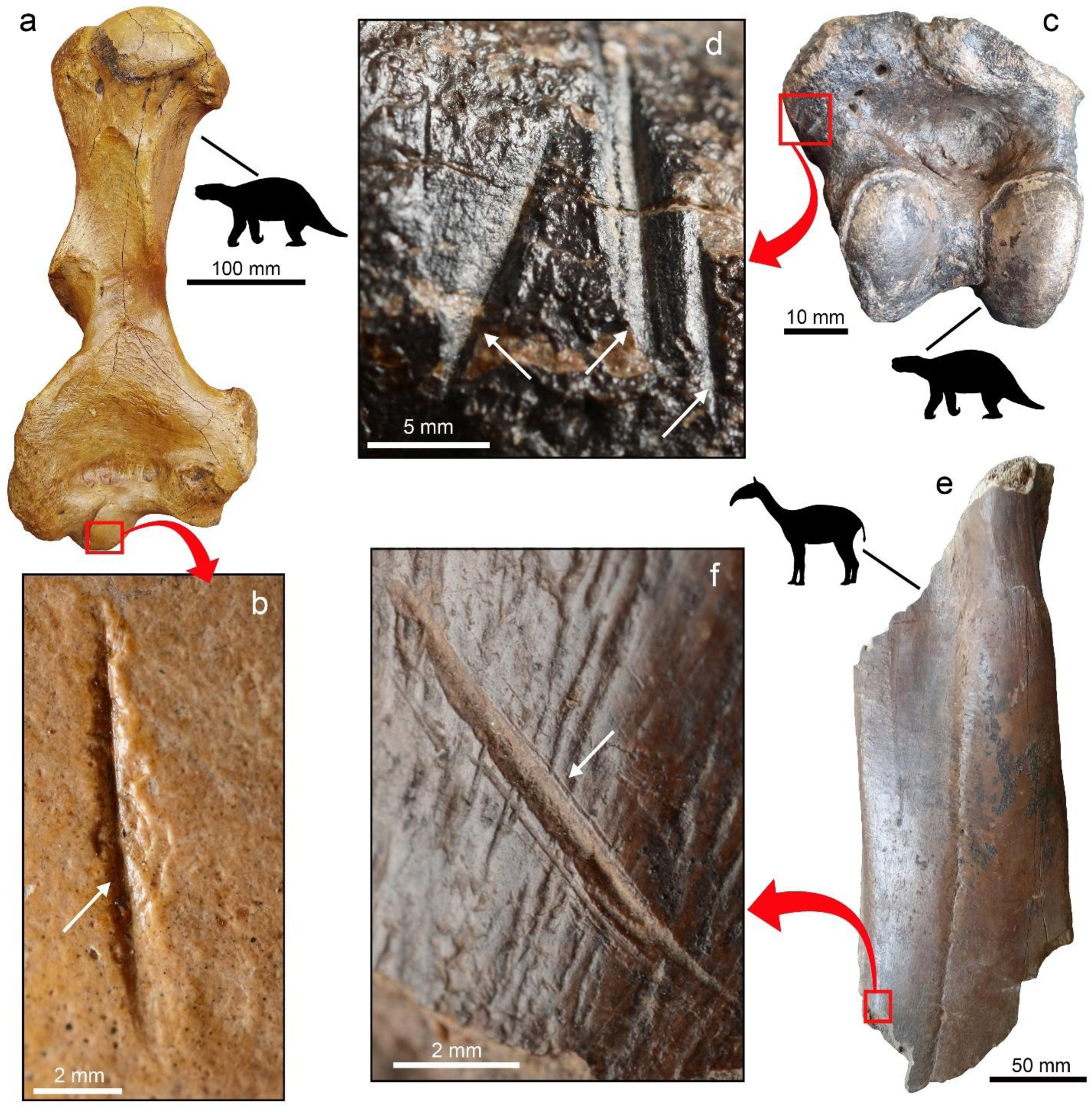
Bones with surface modification from the Taima-Taima site. Left humerus in posterior view (a: IVIC-AP-035) of *Glossotherium* cf. *robustum* with a chop mark on the capitulum (b). Indeterminate Mylodontidae phalange II in ventral view (c: MCNC-Pal-1831) with V-shaped cutting marks (d). Fragmentary right fused ulna-radius in anterior view (e: MCNC-Pal-1834) of cf. *Xenorhinotherium bahiense* showing a cutting mark (f). White arrows show the cut marks. For more detailed information and views on marks and modified surfaces see electronic supplementary material S4.

**Figure 4.**
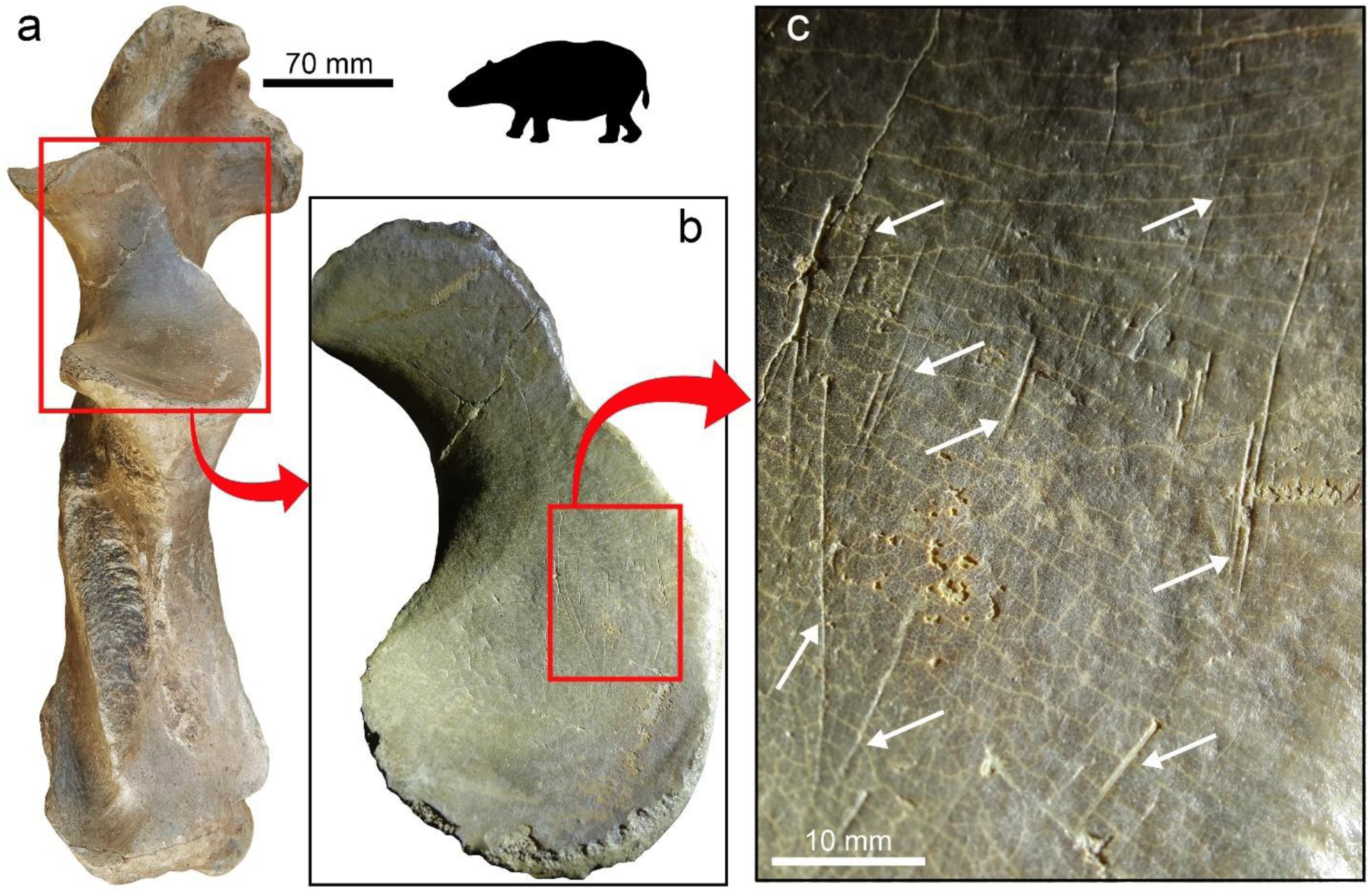
Left ulna (c: CIAAP-1533) of an indeterminate toxodontid with surface modifications from the Taima-Taima site. The ulna in lateral view (a) shows a set of parallel cut marks in the articular facet of the trochlear notch. White arrows show some of the marks. For more detailed information and views on marks and modified surfaces see electronic supplementary material S4.

**Figure 5.**
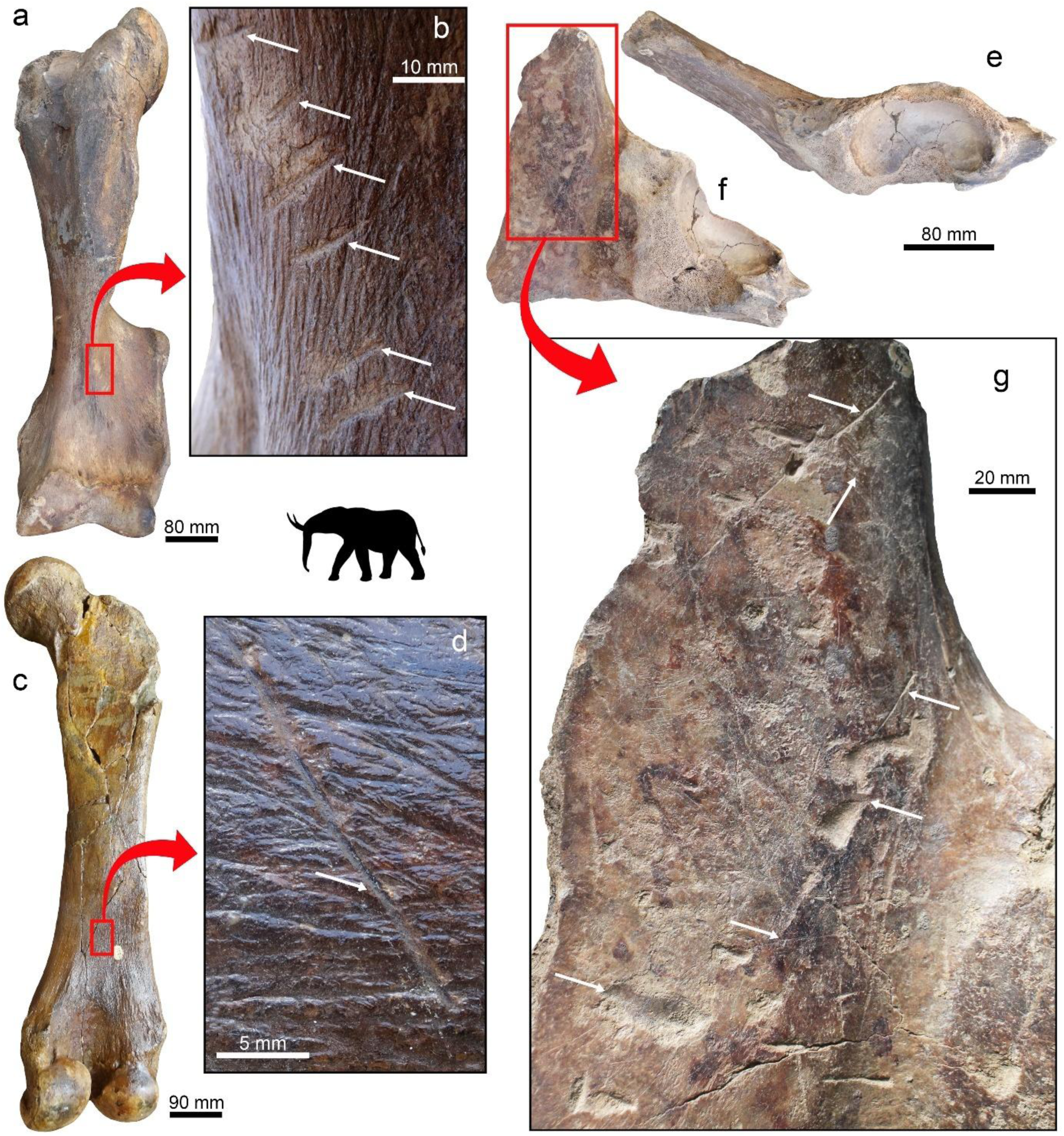
Modified postcranial remains of *Notiomastodon platensis* with surface modifications from the Taima-Taima site. Left humerus in anterior view (a: CIAAP-46) showing six cut marks on anterior surface above the base of the humeral crest (b). Right femur in posterior view (c: CIAAP-111) with a cut mark (d). Left ilium fragment (e–g: CIAAP-19-233) preserving a group of linear and depressions with loss of cortical material produced by likely by percussion. Specimens CIAAP-46 and –111, belong to the same semi-articulated adult individual collected during the 1976 excavation. White arrows show cutting marks and depressions. For more detailed information and views on marks and modified surfaces see electronic supplementary material S4.

Our examination suggests that bone modifications from Taima-Taima (electronic supplementary material 4) are present in four major groups of mammals extending from the Basal to the Upper stratum of the site (Fig. 2). These include a terrestrial sloth (*Glossotherium* cf. *robustum*) and other indeterminate mylodontid (based on the isolated phalanx, Fig. 3), a glyptodont (*Glyptotherium* cf. *G*. *cylindricum*; see Carlini et al., 2012), a macrauchenid (cf. *Xenorhinotherium bahiense*), a proboscidean (*Notiomastodon platensis*), and an indeterminate toxodontid (probably *Mixotoxodon*) (electronic supplementary material S3 and S4). The new evidence suggests that at least an individual of *G*. *robustum* (Fig. 3a) and an undetermined mylodontid (Fig. 3c), an individual of cf. *X. bahiense* (Fig. 3e), at least three of *N*. *platensis* (Fig. 5, and electronic supplementary material S4), and another of a toxodontid (Fig. 4) were likely exploited on the site. These are conservative hypotheses, as additional cases of anthropic marks are potentially recorded from the site (electronic supplementary material S4).

### 4.2 Chronology

The twenty-nine bone/dentine samples from Taima-Taima for radiocarbon dating on collagen, and analyzed at the ORAU, yielded negative results. Of the 29 samples prescreened for bone/dentine collagen preservation prior to radiocarbon dating, all had values well below 0.76% N, at <0.2%. This suggests poor preservation and likely insufficient collagen content for radiocarbon dating. Given the possibility of false negatives in the prescreening method used (Jacob et al., 2018), seven of the 29 samples that showed >0.1 %N were subjected to collagen extraction. Unfortunately, no one yielded any collagen following processing. Poor collagen preservation at Taima-Taima is unsurprising, however, given the environmental and burial conditions—high temperature and water action—which are known to have a negative impact (Hedges, 2002).

Seven radiocarbon dates were obtained on tooth enamel samples (Table 2; electronic supplementary material S2). These yield an age of 15,315–17,810 cal yr BP. Given the likelihood of young carbon uptake during the fossilization process (Zazzo and Saliège, 2011; Zazzo, 2014), particularly in an environment with considerable water action, these ages are likely minimum age estimates and, as such, imply a potential for greater antiquity. The dated specimens included two *Equus* teeth (MCH-Pv-315 and –316) coming from the middle to upper part of the Upper stratum (Fig. 2) were dated to 15,315–15,700 and 15,455–15,830cal yr BP. The anatomical features and proximity of these isolated teeth in the same layer and overlapping ages suggest they belong to the same individual, however. Additionally, five individuals of *N*. *platensis* from the Medium and Basal strata collected during the excavation campaigns of 1960s and 1970s (electronic supplementary material S3) show a wider range of ages (Table 2). CIAAP-83-1486—a molar of the mandible of the 1976 skeleton in which a fragment of the El Jobo projectile was found embedded in the pelvic area—was dated to 16,330–16,695 cal yr BP. This skeleton was excavated from the lower part (saturated grey sand) of the Middle stratum and the previously dated twigs were associated as potential stomach contents of this individual (Bryan et al., 1978; Bryan, 1979; electronic supplementary material S5). Molars CIAAP-313 and CIAAP-80-215 were collected in the Basal stratum during the 1976 excavation (Casamiquela, 1979) and yielded an age of 16,760–17,090 and 17,325–17,795 cal yr BP, respectively. Two isolated molars (IVIC-AP-030 and –033) from the 1960s–early 1970s campaigns were dated to 17175–17,810 and 17,010–17,345 cal yr BP. Although Cruxent’s records do not detail the stratum from which these two molars originate, the age obtained is like the other molars from the Basal and/or Medium strata (Table 2).

### 4.3 Dietary analysis of *Notiomastodon platensis* at Taima-Taima

In total, 19 phytoliths and eight starch grains were recovered from the dental calculus of *N. platensis* (electronic supplementary material S6). We identify five plant taxa from the phytoliths: arboreal dicotyledons, Poaceae, Marantaceae, Asteraceae and Arecaceae (Fig. 6), with the latter two only found in a single sample each. The non-diagnostic arboreal dicotyledons (six samples) are represented by spheroid granulates and sclereids, while Poaceae morphotypes (three samples) include Bulliforms and a single occurrence of a broken lobate phytolith, considered here as belonging to the Panicoideae. The Marantaceae phytoliths include conical to tabulate verrucate forms, consistent with the type typically produced in roots and rhizomes. The Asteraceae morphotype corresponds to those produced in the seeds or inflorescences of this clade. The most likely origin of the spheroid spinulose phytolith assigned to the Arecaceae from the IVIC-AP-034 molar sample is from a palm fruit or seed.

**Figure 6.**
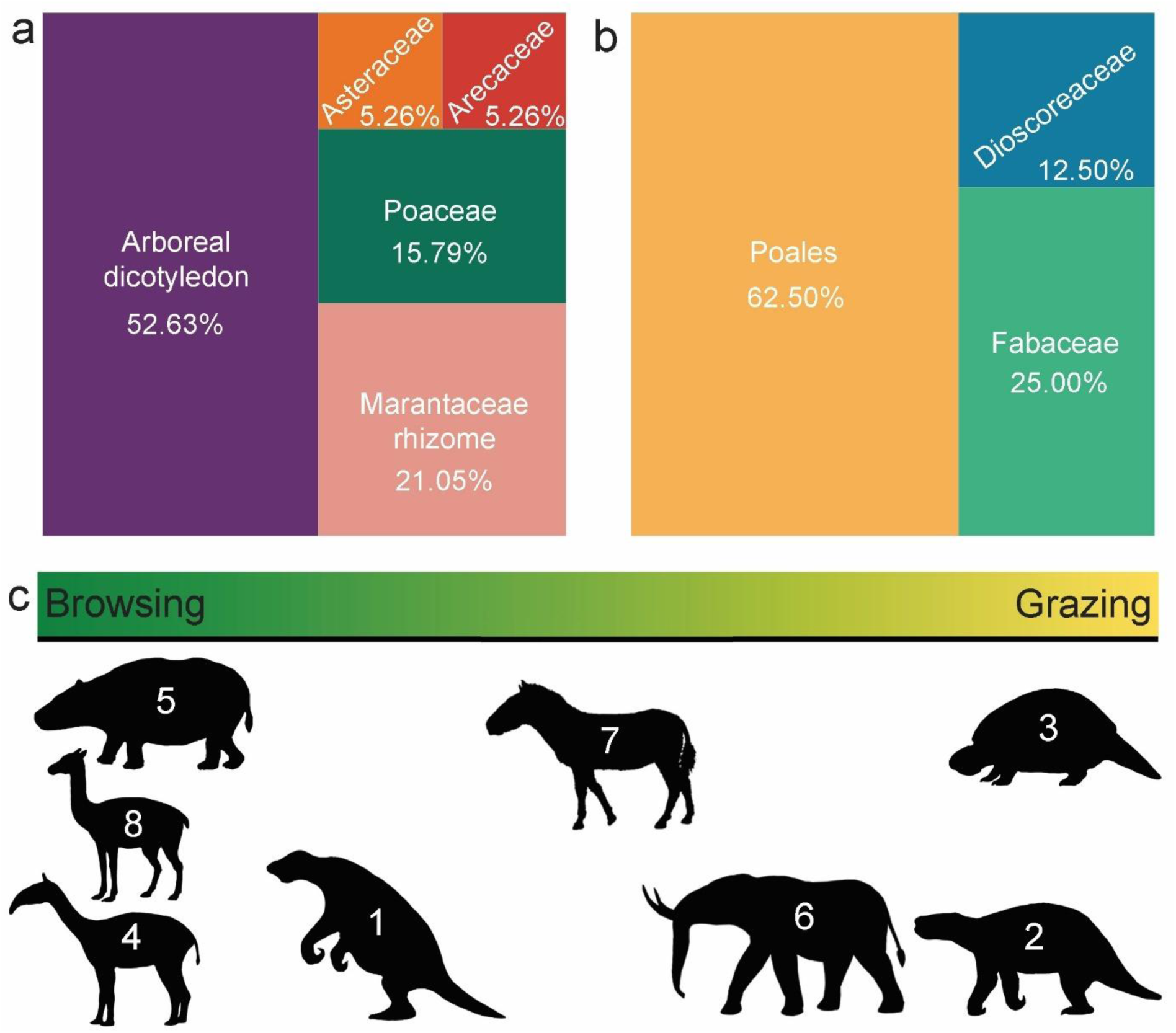
Dietary ecology of the herbivores of Taima-Taima. Tree maps showing the frequency of different plant taxa recovered from the dental calculus analysis of *Notiomastodon platensis* from Taima-Taima, considering phytoliths (a) and starch grains (b). The area of each colour block is proportional to the percentage of either phytoliths, or starch grains assigned to each taxon. Schematic representation of the inferred dietary ecology niche of the megaherbivores present in Taima-Taima (c), using dental calculus analysis, mesowear and reported diets from other localities (electronic supplementary material S6, electronic supplementary material S7). Taxa reported: *Eremotherium laurillardi* (1), *Glossotherium* cf. *robustum* (2), *Glyptotherium* cf. *G*. *cylindricum* (3), cf. *Xenorhinotherium bahiense* (4), *Mixotoxodon* (5), *Notiomastodon platensis* (6), *Equus* sp. (7), and *Palaeolama major* (8). Silhouette of *Equus* sp. from phylopic.org: by Zimices (CC BY-SA 3.0).

Three plant taxa were identifiable from the starch grains: cf. Poales, cf. Fabaceae and cf. Dioscoreaceae. Of these, the most common were the grains of the Poales (grasses and sedges), which were recovered from five of the eight samples. The starch grains were generally small to medium in size, predominantly spherical to polyhedral, and featured mostly centric hila. In some cases, they showed faint extinction crosses, characteristics consistent with those of grasses. Two samples include large (∼20 µm), ovoid to rounded starch grains with fissures around the hilum, assigned to the Fabaceae (beans and legumes). A single sample (CIAAP-1481) includes starch grains attributable to Dioscoreaceae (yams). Because we do not recognise any sedges or other Poales within the phytolith sample, we consider it most likely that the Poales starches derive from Poaceae. This suggests that six of the samples (75%) contain dental calculus being evidence of grass consumption.

## 5. Ancient palaeoenvironment

When the megafauna and humans were present at Taima-Taima, the site was presumably located between estimated 15 and 20 km from the coastal area, because of the low sea level at the Late Pleistocene time (Fig. 1). The region was characterised by a greater negative water balance with a predominance of savannah areas (Ochsenius, 1979, 1980). The watering hole, probably permanent throughout the year, likely offered an oasis that attracted animals during dry periods. The close association of extinct proboscideans with water resources is well-documented in the literature (Haynes, 1991; Shoshani and Tassy, 1996; Pasenko and Lucas, 2011), including *N. platensis* (Mothé et al., 2010; 2022; Lindsey and Lopez, 2015; Labarca et al., 2024). Due to their large body size and high-water requirements, these animals were naturally attracted to areas offering essential water and mineral sources, such as rivers, lakes, and groundwater springs. This ecological preference likely played a significant role in shaping their distribution and migration patterns, leading them to congregate around water holes and springs and, in many localities, encountering humans due to the same necessities; availability of freshwater springs during dry seasons or scarcity of surface water (Plint and Magill, 2021; Haynes, 2022). In the nearby watering holes of Muaco and Cucuruchú (Royo y Gomez, 1960; Cruxent, 1970; Aguilera, 2006; Carrillo-Briceño, 2015), the presence of *N*. *platensis* in association with other megaherbivores has also been reported (Carrillo-Briceño et al., 2024).

The megaherbivores from Taima-Taima contain representatives of a variety of dietary niches (Fig. 6), which reflect diversity in the available dietary vegetation. Wilson et al. (2024) used mesowear angle approaches to quantify diet in *N. platensis* and *E. laurillardi* from Taima-Taima, finding that the *N*. *platensis* specimens from this site possessed relatively shallow dentine valleys, with a mean mesowear angle of 128.4°, indicating a high level of dietary abrasion, consistent with grazing (Saarinen and Lister, 2023). Our dental calculus analysis provides some support for this interpretation, with 75% of samples showing evidence of grass consumption (Fig. 6). However, the phytoliths and starch grains reveal a diverse range of plants consumed by *N*. *platensis*, with 75% of the samples including unidentified arboreal dicotyledons. Dental calculus samples of *N*. *platensis* suggest a higher consumption of grasses at Taima-Taima than at other South American localities (e.g., Asevedo et al., 2012; González-Guarda et al., 2018, 2025), though with rare fruit consumption, as supported by these other studies. As for other herbivores, the diet of proboscideans tends to reflect their environment (Robinson et al., 2021; Abraham et al., 2024; González-Guarda et al., 2025), so proboscidean mesowear, as a proxy for diet, can be applied to reconstruct vegetation cover (Saarinen and Lister, 2016, 2023; Xafis et al., 2020; Foister et al., 2024; Wilson et al., 2024). In this case, the mesowear in *N. platensis* suggests high availability of grasses in the area, though with a diversity of other plants available, consistent with a savannah reconstruction. The other herbivores from Taima-Taima are consistent with this interpretation (electronic supplementary material S7). Wilson et al. (2024) found that the mean mesowear angle of the two *E. laurillardi* specimens from Taima-Taima was 106.5°, which suggested a browse-dominated mixed feeding dietary niche (Saarinen and Karme, 2017). This study also found a low-abrasion diet in the toxodontid *Mixotoxodon larensis* from the nearby Muaco locality, probably indicating a browsing niche, consistent with prior results from carbon isotopes (MacFadden, 2005). The single dental specimen of *G*. *robustum* has a mesowear angle of 152.5°, reflecting a grazing preference Wilson et al. (2024). Prior studies from other regions have produced different results regarding the diets of the other megaherbivores reported from Taima-Taima. De Oliveira et al. (2020) suggested a grazing niche for *X. bahiense* in Brazil based on microwear and occlusal enamel index analyses, whilst Omena et al. (2020) found more likely a higher consumption of C_3_ plants for *Xenorhinotherium*, suggesting a browsing niche, while reconstructing mixed-feeding and grazing dietary niches for *Palaeolama* and *Equus*, respectively, in the Brazilian Intertropical Region. Dental material from nearby Muaco suggests a browse-dominated diet for *Xenorhinotherium* and *Palaeolama*, while *Equus* specimens have a variable dietary signal, consistent with graze-dominated mixed-feeding (electronic supplementary material S7). Omena et al. (2020), Dantas et al. (2020) and Lessa et al. (2021) found a preference for C_4_ plants for *Glyptotherium*, suggesting that this glyptodont was likely a grazer. Dantas et al. (2020) predicted, for example, a diet of 77 % grasses, a value coincident with a savannah environment.

Direct quantification of the diet of these other herbivore taxa from Taima-Taima will help us to further understand the environment here at the time. However, combined, the megaherbivores in Taima-Taima (Fig. 6) suggest a palaeoenvironment with a range of vegetation types. Our results support a dominant grassy component in this region at the end of the Pleistocene based on the mesowear signal from *N*. *platensis*, though there were also several browsing herbivores (i.e., *Xenorhinotherium*, *Eremotherium*, *Palaeolama* and *Mixotoxodon*). We therefore suggest that the surroundings of the Taima-Taima site was an open savannah with low trees and shrubs available for browsing taxa.

The current vegetation at Taima-Taima and the greater region is dominated by xerophytic shrubland, characteristic of most of the northern Falcón State (Ochsenius, 1980). The early Spanish colonization, with the introduction of goats, may have environmentally affected the region, but the dominance of this type of vegetation is surely not an effect of it, as historical records indicate that this vegetation was predominant at the time of arrival of the first Europeans to the region (Jaramillo et al., 2020). Likewise, palaeobotanical records from Taima-Taima show the presence of some plant species (*Coccoloba uvifera*, *Portulaca oleracea*, and *Prosopis*, *Caesalpidium* or *Cercidium*) that are dominant in the current dry ecosystem of the Falcón coast, and which matches the expectations from the herbivores (Ochsenius, 1980). The vertebrate and geological record of the late Neogene Falcón Basin, as is the case of the Cocinetas Basin in neighbouring Colombia, suggest that aridity has been a feature only over the past ∼2 Ma (Pérez-Consuegra et al., 2018; Carrillo et al., 2018; Jaramillo et al., 2020; Scholz et al., 2020). Establishing when and how this change occurred and how it affected the biota is still a matter to be resolved (Wilson et al., 2024). Combined with the plant fossil record, the megafauna at Taima-Taima is an indicator of a relatively dry, though possibly more humid environment in the Late Pleistocene than today.

## 6. Taima-Taima, the hunting ground

Evidence of human-megafaunal interactions can be supported in the form of (1) physical associations, which simply refer to bones and tools found side by side, or in the same deposit; and (2) behavioural associations, represented by the presence of cut marks and other bone modifications (e.g., Fernández-Jalvo and Andrews, 2016), which represent direct interactions (Borrero, 2009).

Taima-Taima preserves both type of evidence in the human exploitation of different megaherbivore species. Previous assumptions documented the exploitation in the site of the proboscidean *N*. *platensis* (Bryan et al., 1978; Bampi et al., 2022; Haynes, 2022) and the giant armadillo *Glyptotherium* cf. *G*. *cylindricum* (Carlini et al., 2022). Our results support evidence of at least three other species of megaherbivores, including *Glossotherium* cf. *robustum*, cf. *X. bahiense*, and an indeterminate toxodontid (probably *Mixotoxodon*), that were likely exploited by humans in Taima-Taima. The evidence preserved at Taima-Taima also support the hypotheses that both killing and slaughtering activities were likely practised by humans at the site (Bryan et al., 1978; Carlini et al., 2022).

The physical associations of bones and tools from Taima-Taima are restricted so far only to three *N*. *platensis* individuals in close association with lithic artifacts. The first report comes from the excavation of 1968 and corresponds to a chert scraper found next to a hemimandible (Cruxent, 1978; electronic supplementary material S1). The other two *N*. *platensis* individual were potentially victims of projectile weapons, a hypothesis that has been substantiated by the presence of two El Jobo projectile points in association with their pelvic elements (electronic supplementary material S1). Both individuals were found at 18 m of each other, as reported from the 1974 and 1976 excavation campaigns (Cruxent, 1978; 1979; Bryan et al., 1978). The pelvis and associated artifact from the excavation of 1974, was displayed for many years in the museum Jose Maria Cruxent Exhibition Room of the IVIC (electronic supplementary material S1). Nevertheless, its current condition is uncertain, because its original display is covered by a diorama, which prevented our study and documentation of possible bone modifications. In contrast, the individual from the excavation of 1976 (dated here to 16,330–16,695 cal yr BP; Table 2) corresponds to a semi-articulated and incomplete skeleton, including a little more than 70 bone elements (Bryan et al., 1978; Casamiquela, 1979), with one chert scraper found adjacent to its left ulna and a second one with a rib (Cruxent, 1978, 1979; electronic supplementary material S1). Casamiquela (1979) noticed some behavioural associations represented in the form of six cut marks (reported here as chop mark) on left humerus (Fig. 5a), and other probable cut marks on two ribs. Here, for the first time, we report evidence of modifications of probable anthropic origin (Table 1; electronic supplementary material S2 and S4) also in the left scapula, right femur and tibia of the 1976 skeleton (Fig. 5c), with different types of modifications that could be related to chop, slicing and cutting marks, which likely suggest butchering and dismemberment of the individual.

Other three isolated postcranial elements of *N*. *platensis* with evidence of probable anthropogenic nature were also here observed. These are represented by two scapulae and an ilium fragment originating from the Basal and Medium strata (Table 1; electronic supplementary material S2 and S4). Probably the most representative of all these is the latter (electronic supplementary material S1 and S4), with a variety of modifications (Fig. 5e–g). Evidence appears to be chop, cutting and slicing marks (some of them parallel), including loss of cortical material likely produced by percussion (see Figs. S4.13–16). These modifications are not isolated evidence restricted only to that bone element. On the contrary, they seem to be present in other medium-large bones from Taima-Taima (Table 1). Casamiquela (1979: 71–72 pp.) presented a detailed discussion of the nature and taphonomic aspects of this kind of depressions and loss of cortical material preserved in some limb bones. For example, he suggested that femora CIAAP-90 and CIAAP-58-307 collected from the Basal stratum (figured in electronic supplementary material S4), were likely used as anvils, possibly for chopping meat the latter is based on the extreme modifications and loss of cortical material caused by percussion blows. We hypothesize that these two femurs could preserve evidence/modifications that are not of natural origin, however, they should be studied in more detail for confirmation. Percussion depressions or blows identified in some bones from Taima-Taima (see the ilium fragment illustrated in electronic supplementary material S4) may have been produced using a heavy tool, probably made of a hard material, coinciding with the presence of some potential artifacts found in Taima-Taima. Cruxent (1967, 1978, 1979) reported lithic artifacts referred to as expedient tools, which were probably used for a specific task once, or a few times and then discarded. These tools were somehow modified using the same rocks (pebbles) that make up the Basal stratum and could have functioned as striking objects (Cruxent, 1979).

*Notiomastodon platensis* is the most abundant species in Taima-Taima, with a minimum of 14 individuals, including 4 immatures (calves), 2 subadults, 3 adults, 2 mature adults, and 3 senile adults (electronic supplementary material S3). Despite these challenges, evidence suggests that *N. platensis* individuals of different ages (according to post-canine teeth type and wear stage of each tooth; Mothé et al., 2010) probably were exploited at Taima-Taima. For instance, the individual represented by the 1976 skeleton (associated with El Jobo projectile points in their pelvic region) appears to have been between 29 and 35 years old (Fig. 7; electronic supplementary material S3). Apart from the 1976 skeleton, our new evidence suggests that, at least, other individuals of *N*. *platensis* were probably exploited by humans at the site (electronic supplementary material S4). Taking as reference only the pelvic elements and the right scapula of the 1976 skeleton, together with the other two pelvic elements (left ilium CIAAP-19-233, and complete pelvic bone from the 1974 excavation with associated El Jobo projectile), or the other two right scapulas (CIAAP-119-110-210 and CIAAP-213-230; electronic supplementary material S2), all of them with potential evidence of anthropogenic modifications, we would be referring to a conservative approximation of at least three individuals. This figure may be underestimated, as assigning the modified bones elements to specific individuals from Taima-Taima is challenging due to their condition as isolated elements, poor preservation in some cases, the absence of other elements not recovered during different excavation campaigns, and absence of stratigraphic context in some cases. An example is the hemimandible excavated in 1968 and associated with a chert scraper (supplementary electronic material S1), whose whereabouts and stratigraphic context are unknown, not allowing us to determine if it is associated with a particular individual, like the pelvis with a El Jobo projectile excavated in 1974.

**Figure 7.**
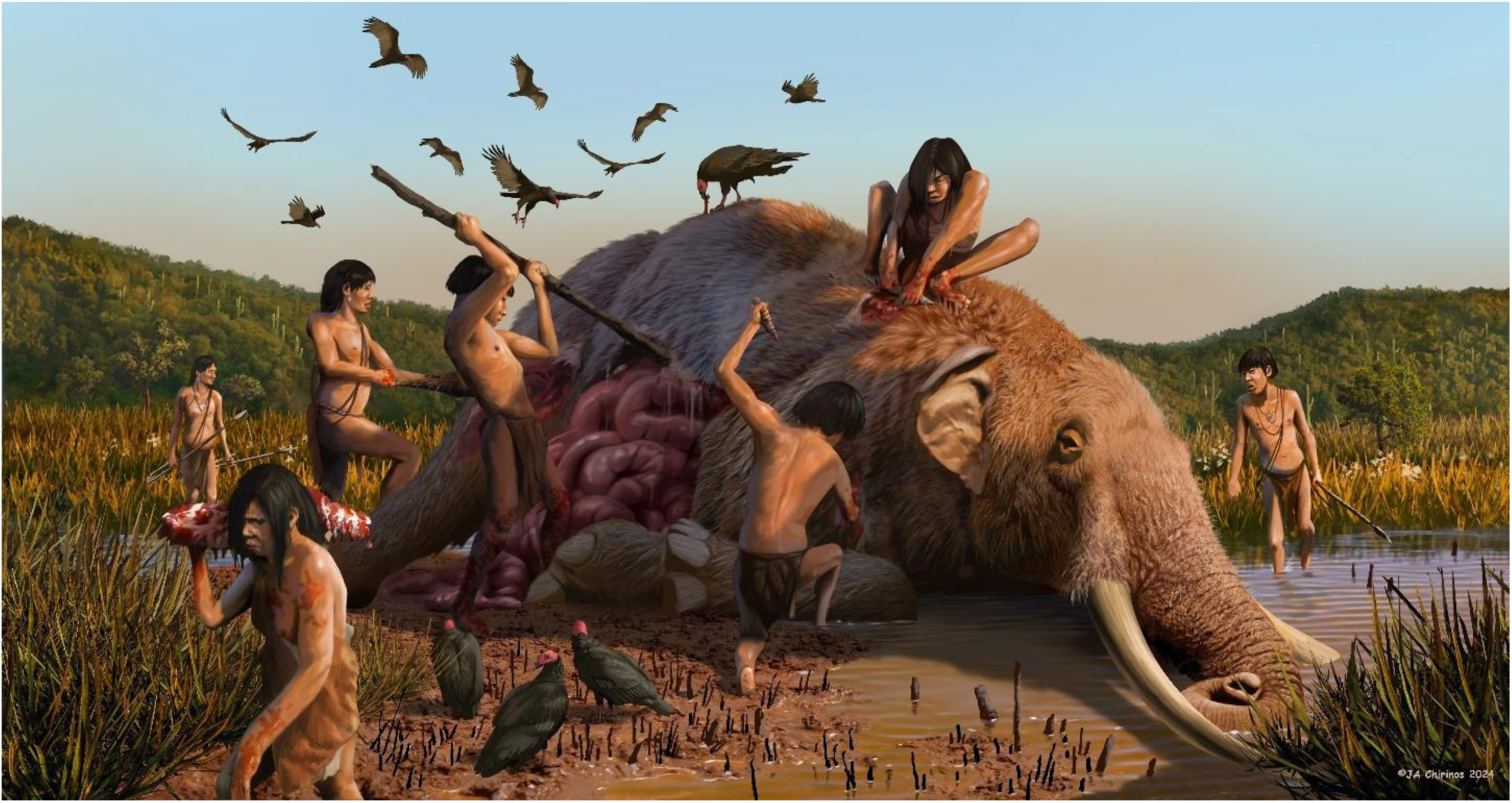
The butchering of *Notiomastodon platensis* in the ancient Taima-Taima watering hole at the end of the Pleistocene. Image credit: Jaime Chirinos 2024.

Proboscideans could be considered high-value prey due to their massive size, which could provide a substantial amount of food (meat, fat, organs and marrow) to sustain a large group of humans (Mothé et al., 2020); although some research ranked them as a prey with low values in calories per kilogram and high pursuit and handling costs (see Prates et al., 2025). Additionally, the proboscidean ivory and bones served as essential raw materials for crafting tools (e.g., Carrillo-Briceño et al., 2025). Even when targeting smaller or younger individuals, the rewards outweighed the risks and efforts involved in hunting them (Lupo and Schmitt, 2023). The successful capture of a gomphothere offered a crucial survival advantage, particularly in facing the climatic and environmental changes of the Late Pleistocene. In addition, due to the presence of water resources in the region, attracted *N. platensis* herds possibly increased encounter rates between proboscideans and humans, reducing prey search-time and increasing chances of resources procurement (Byers and Ugan, 2005). This may explain why *N. platensis* is the most common taxon found among the fossil remains from Taima-Taima site, although its presence is restricted to the period that encompassed the accumulation of the Basal and Medium strata (Casamiquela 1979; Cruxent, 1979).

Five skulls of *Glyptotherium* cf. *G*. *cylindricum* with patterns of breakage by intentional blows (Carlini et al., 2022) suggest that these glyptodonts were likely hunted and subsequently butchered at Taima-Taima, supported by the two inverted carapaces found at the site suggest (electronic supplementary material S1 and S3). Other megaherbivores remains from Taima-Taima with probable modifications of anthropogenic origin occur in two or more strata. (Fig. 2, electronic supplementary material S2). The limb of the adult individual of *Glossotherium* cf. *robustum* with cut marks (Fig. 3a, b) comes from the Upper stratum, while the phalanx with chop marks (Fig. 3c, d) was collected in the Medium stratum. The limb bone of cf. *Xenorhinotherium bahiense* (probably adult Fig. 3e, f; see electronic supplementary material S3), was collected in the Medium stratum, while the subadult individual of the indeterminate toxodontid (Fig. 4, probably *Mixotoxodon*) comes from the Upper stratum. There is no evidence that the bone elements of the taxa mentioned above have been found in association with any lithic artifact. However, the modifications identified in these elements are characterized by us as linear marks (Table 1, electronic supplementary material S4), which are likely associated with cutting activities. These types of modifications do not allow us to assert that these megaherbivores were hunted, but they were likely butchered at the site. The cut marks on most of these elements are preserved in areas of articulation and tendons (e.g., Figs. 3–5), which are likely to reflect a carcass dismemberment.

Other megaherbivores such *Equus* and *E. laurillardi* have been also reported for Taima-Taima (Cruxent, 1970; Casamiquela, 1979; Aguilera, 2006) and although in the first case abundant, we found no evidence of exploitation for these two taxa. However, there is evidence that *E*. *laurillardi* was exploited in the region. At the Late Pleistocene site of El Vano, in the Venezuelan Andes, about 190 kilometers south of Taima-Taima, an *E*. *laurillardi* individual was hunted and butchered by hunters carrying the El Jobo-type projectiles (Jaimes et al., 2024b), same technology reported for Taima-Taima.

## 7. Final Remarks

During the Late Pleistocene, Taima-Taima likely acted as an oasis attracting a diverse community of grazers and browsers. The discovery of middle-sized to megamammals species in sites in this region of the northern Neotropics indicates that resurgent springs were recurrent spots for native faunal communities, as seen currently in African savannahs (Ferry et al., 2016), and resource-procuring humans (Miotti, 2006). Our analyses suggests that, during this period, humans exploited proboscideans (Fig. 7) and other megamammals that congregated at the watering hole. These repeated encounters may have fostered the development of specific hunting and/or predation strategies. This is partly reflected in the diversity of El Jobo projectile points found in Taima-Taima and the region (Vargas et al., 2025), in some cases in direct association with the megafauna (Jaimes et al., 2024a). Combined evidence at the site suggests that the exploitation of megaherbivores was probably not limited to a single event or short period but likely lasted as long as the total estimated duration of Basal, Medium and Upper strata. Conservatively, however, the timing of human-megafauna interaction can only be attributed to CIAAP-83-1486— the only directly dated human-modified specimen at the site—with the enamel age of ECHo 7502 (Table 2) and associated twig measurements (electronic supplementary material S5) dating to 16,330–16,695 cal yr BP (see electronic supplementary material S1).

Data on dental mesowear analysis, phytoliths and starch grains recovered from *N. platensis* molars suggest a higher consumption of grasses than recorded for this species in other sites. This indicates high availability of grasses in the area, though with a diversity of other plants available, and is consistent with a savannah reconstruction as supported by other herbivores from Taima-Taima based on dental ecometrics (Wilson et al., 2024). The disappearance of megafauna at the site likely occurred sometimes after the deposition of the Upper stratum, since no other fossils were found in the overlying sediments (Bryan, 1979; Casamiquela, 1979; Gruhn and Bryan, 1984).

## Supporting information

Electronic supplementary material S1

Electronic supplementary material S2

Electronic supplementary material S3

Electronic supplementary material S4

Electronic supplementary material S5

Electronic supplementary material S6

Electronic supplementary material S7

## Acknowledgements

We thank Nohe Gilson, Rodolfo Sánchez, Jorge Rivas, Hyram Moreno, Edwin Chávez-Aponte, and Miguel Zabala for their valuable support in the field and collection activities, the Reyes family and community of Taratara for the valuable support, and the Instituto del Patrimonio Cultural de Venezuela for kindly providing permits for field activities and study of collections and for the temporary export of samples for dating studies. We also thank the Oxford Radiocarbon Accelerator Unit, Samreen Syeda (BUT Chimie Paris Saclay) and François Thil (Laboratoire des Sciences du Climat et de l’Environnement) for assistance and radiocarbon dating. We also thank the Latin American Center at the University of Zurich for its support with the GRC Travel Grant 2019 and Mobility Grant 2022, and the Georges und Antoine Claraz-Donation Grant (2024) granted to J.D.C-B. LBV thanks the Leverhulme Trust Fund (Early Career Research Fellowship ECF-2022-532) for funding her time on this research.

## Electronic supplementary material

**S1**. Notes on the history of studies of the Taima-Taima site.

**S2**. Fossil specimens from Taima-Taima.

**S3**. Notes on taxonomic assessment.

**S4**. Surface modification of bones and other evidence of anthropic origin in megaherbivore taxa from the Taima-Taima site (Late Pleistocene).

**S5**. Previously published radiocarbon dates for the Taima-Taima site.

**S6**. Phytolith and starch grain analysis of dental calculus from the Taima-Taima site.

**S7**. Dietary reconstruction of herbivores from Taima-Taima.

