## Supplementary material for "Diverse megamammals exploited by humans, chronology and palaeoecology at Taima-Taima, Late Pleistocene, South America": Electronic supplementary material S1

**Notes on the history of studies of the Taima-Taima site**

**1. The Taima-Taima site**

In the late 50s, Mr. Ramón Palencia, a local from the town of Taratara, was expanding the water sources in Taima-Taima to improve the drinking troughs for his animals (permanent springs are common in this area), when he found remains of fossil megafauna during the sediment removal process (Carrillo-Briceño, 2015). Upon learning of this discovery, the archaeologist Dr. José María Cruxent visited the site in 1961 and, together with Dr. José Royo y Gómez from the Universidad Central de Venezuela (UCV) in Caracas, observed the potential of the place and planned future excavations. Unfortunately, the delicate state of health of Royo y Gómez and his surprise death, on December 30, 1961, prevented the start of the excavations. Cruxent began the first excavation campaign at Taima-Taima in March of 1962, and due to the wetland condition of the site, he initially named the site Los Pozos de Royo y Gómez (Fig. S1.1) to honor the memory of his deceased colleague (Cruxent, 1967; Cruxent and Ochsenius, 1979; Ardila, 1987). The first campaign was carried out in different seasons, and according to excavation photos shown by Cruxent (1967: lam. 1), and the inverted glyptodon carapace, was recovered during this interval (Fig. S1.2).

A second excavation campaign at Taima-Taima began in 1968, where abundant fossil remains and some lithic artifacts, such as scrapers, were found (Cruxent, 1978, 1979). A chert scraper was found next to a *Notiomastodon platensis* hemimandible (Cruxent, 1978), and apart from the photograph of the specimens in situ (Fig. S1.3), we do not have any further details regarding probably bone modifications, catalogue number or repository location of the hemimandible.

According to Cruxent (1979: 78 p), in the excavation season of 1974 (which was a continuation of the second excavation campaign), a fragment of the El Jobo type projectile point was found in the pelvic cavity of a gomphothere. This specimen and associated artifact (Fig. S1.4a, b) are today housed in the “Jose Maria Cruxent Exhibition Room”, Centro de Antropología of the Instituto Venezolano de Investigaciones Científicas (IVIC), Miranda State, Venezuela, where it was exhibited for many years. However, its current condition is uncertain, because its original display was recently covered by a diorama. An estimated 150 square meters were excavated in Taima-Taima during the first and second campaigns seasons (Cruxent and Ochsenius, 1979; Oliver and Alexander, 2003).

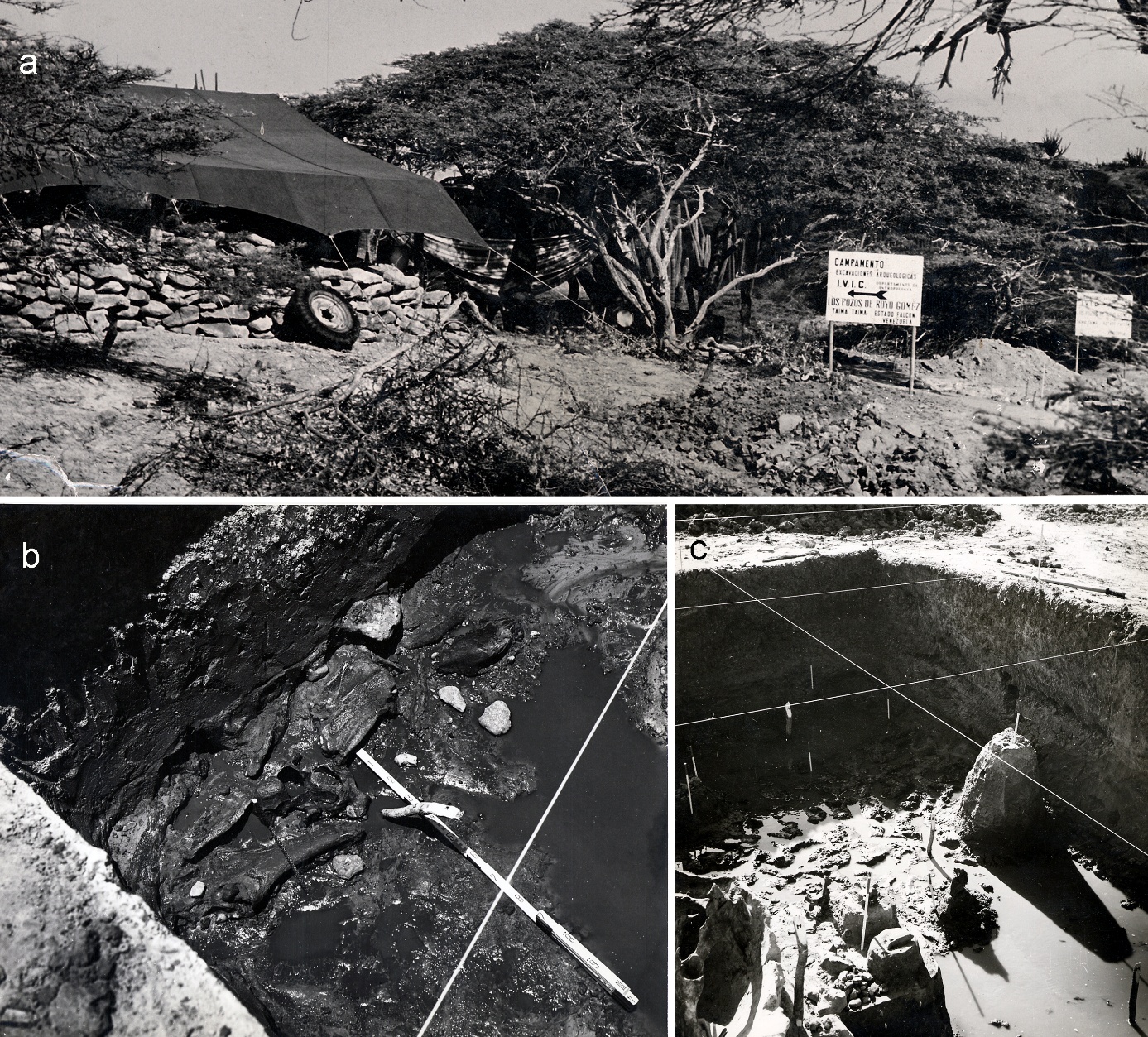

**Figure S1.1**. Excavations at the Taima-Taima “Los Pozos de Royo y Gómez” site during the first campaign. Camping (a) and excavation (b, c). In picture b are exhibited remains of *Eremotherium laurillardi*, including a mandible illustrated previously by Cruxent (1967). Images courtesy archive of the Universidad Experimental Francisco de Miranda (UNEFF).

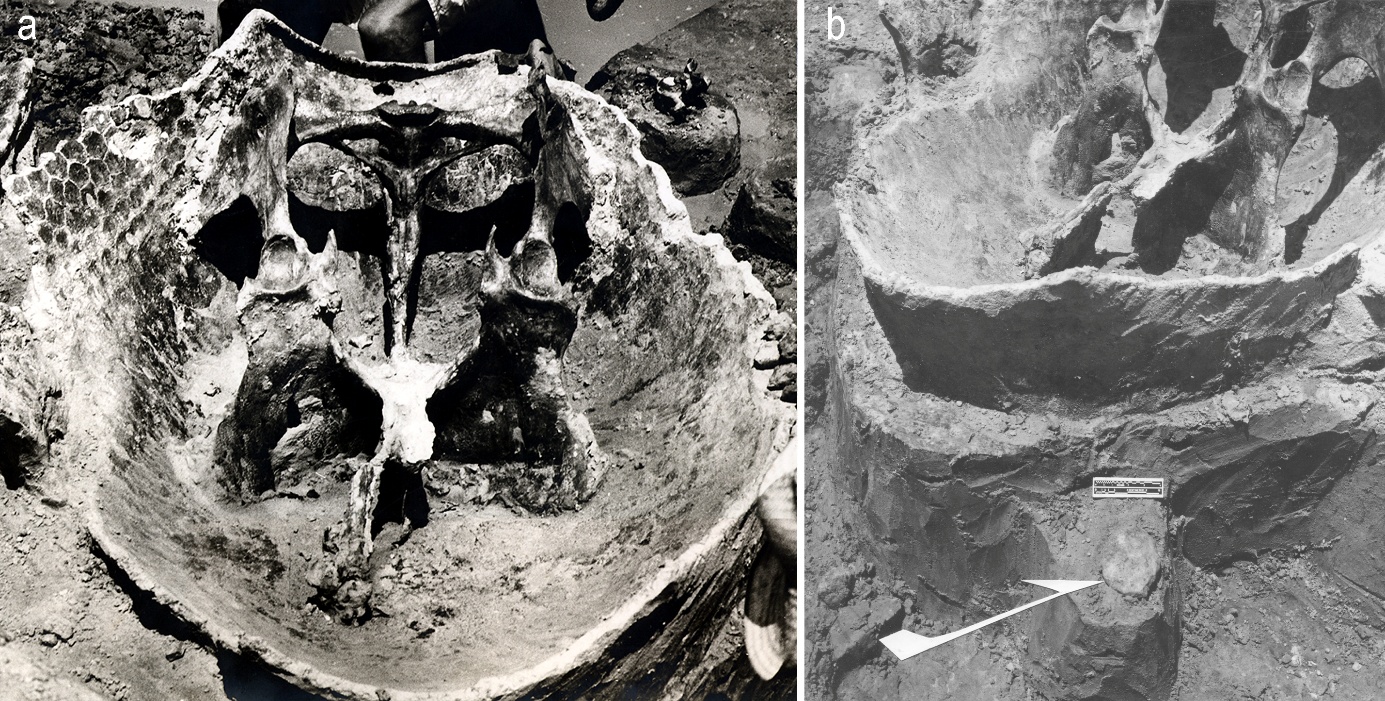

**Figure S1.2**. Excavations at the Taima-Taima site during the first campaign season (a, b). Inverted glyptodont carapace of *Glyptotherium* cf. *G*. *cylindricum*. The white arrow in image b shows a lithic artifact referred to “expedient tools” by Cruxent (1967, 1978, 1979). Images courtesy image archive of the Universidad Experimental Francisco de Miranda (UNEFF); image b modified after Cruxent (1978).

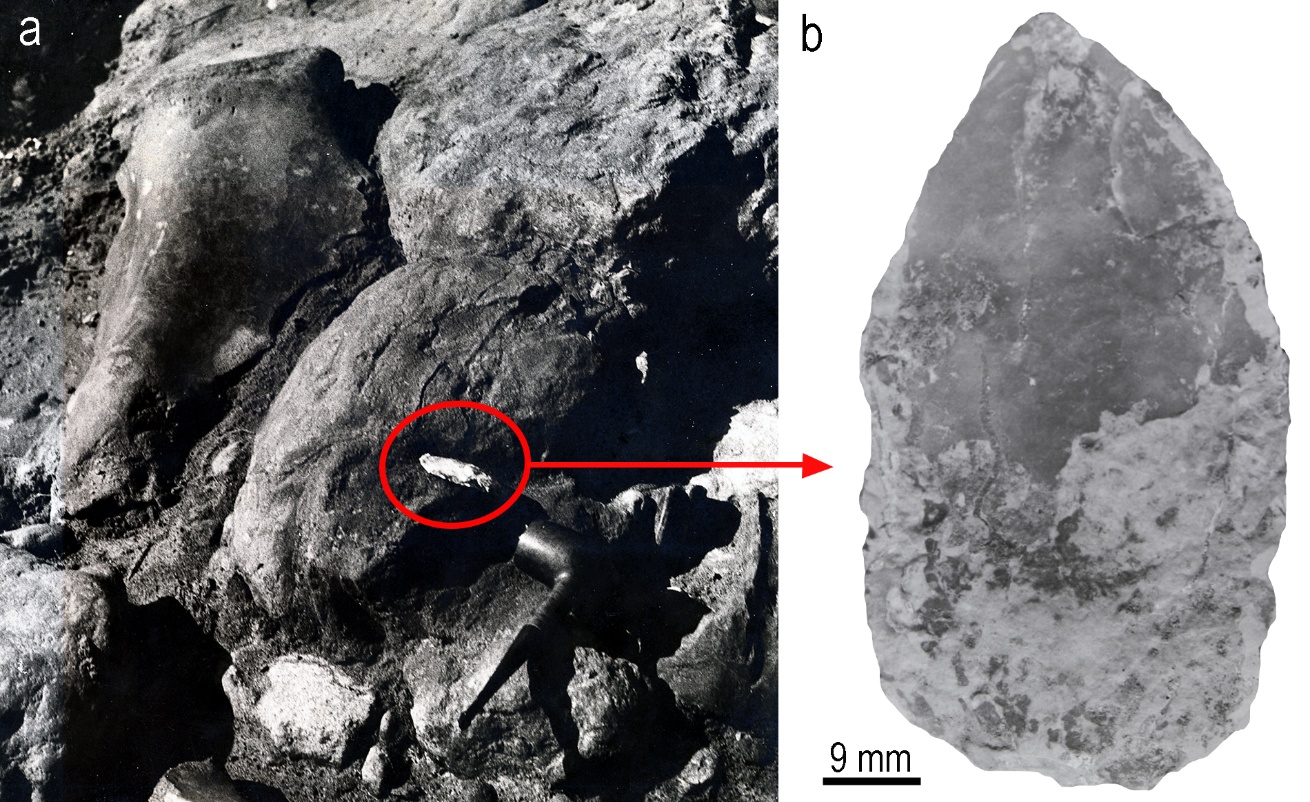

**Figure S1.3**. Hemimandible of *Notiomastodon platensis* (a) with scraper (field n° 3.012) produced in chert (b) *in situ* from the excavation of 1968. Image a, modified after Cruxent (1978), and image b courtesy of the Centro de Antropología del Instituto Venezolano de Investigaciones Científicas (IVIC).

The third excavation campaign started around July 1976, and this had the participation of a multidisciplinary team conformed by Alan Bryan, Claudio Ochsenius, Ruth Gruhn and JM Cruxent himself (Ochsenius and Gruhn, 1979). During this excavation, 80 square meters were added to the previously excavated area (Cruxent and Ochsenius, 1979). Four stratigraphic units were identified: 1) “Fine convoluted sand” (Unit I), 2) “Laminated fine sand” (Unit II), 3) “Black organic sandy clay” (Unit III), and 4) “Brown colluvium” (Unit IV) (Bryan, 1979). Many fossils of megafauna (Figs. S1.4c, 5) and other mammals and giant land tortoises were recovered from the Basal, Medium, and Upper fossiliferous strata, as part of units I and II. According to Casamiquela (1979), Unit I/II disconformity represents the last evidence of megafauna in the Taima-Taima section. During this excavation, a semi-articulated skeleton of a *Notiomastodon platensis* was also found with a fragment (medial) of an El Jobo projectile in its pelvic region (Fig. S1.4c) and a chert scraper associated with one of the limb bones (Fig. S1.6). The faunal assemblage reported for the Taima-Taima (Table S1.1) is characterized by the predominance of mammals of different body sizes, with a predominance of megafauna. This predominance in the site diversity may have resulted from a bias towards larger specimen collection in previous excavation campaigns. A detailed description of the two El Jobo projectiles and the two scrapers found in the different excavations of Taima-Taima is presented in detail by Cruxent (1978, 1979)

**Table S1.1**. Faunal list reported for the Taima-Taima site, and based on Casamiquela (1979), Ochsenius (1980), Bocquentin-Villanueva (1982), Aguilera (2006), Carrillo-Briceño (2015); Carlini et al. (2022); Reyes-Céspedes et al. (2023), and references therein.

| **TAXONOMY** | |
| --- | --- |
| XENARTHRA (Pilosa) |  |
| †Megatheriidae | †*Eremotherium laurillardi* |
| †Mylodontidae | †*Glossotherium* cf. *robustum* |
|  | Indet. |
| XENARTHRA (Cingulata) |  |
| †Glyptodontidae | †*Glyptotherium* cf. *G*. *cylindricum* |
| †Pachyarmatheriidae | †*Pachyarmatherium* cf. *P*. *brasiliense* |
| †LITOPTERNA |  |
| †Macraucheniidae | cf. †*Xenorhinotherium* *bahiense* |
| †NOTOUNGULATA |  |
| †Toxodontidae | cf. †*Mixotoxodon* *larensis* |
|  | Indet. |
| PROBOSCIDEA |  |
| †Gomphotheriidae | †*Notiomastodon* *platensis* |
| ARTIODACTYLA |  |
| Camelidae | †*Palaeolama major* |
| Tayassuidae | Indet. |
| Cervidae | Indet. |
| PERISSODACTYLA |  |
| Equidae | *Equus* cf. †*E*. *neogeus* |
| CARNIVORA |  |
| Felidae | Indet. |
| Ursidae | †*Arctotherium wingei* |
| CHIROPTERA |  |
| Phyllostomidae |  |
| TESTUDINES |  |
| Testudinidae | *Chelonoidis* sp. |

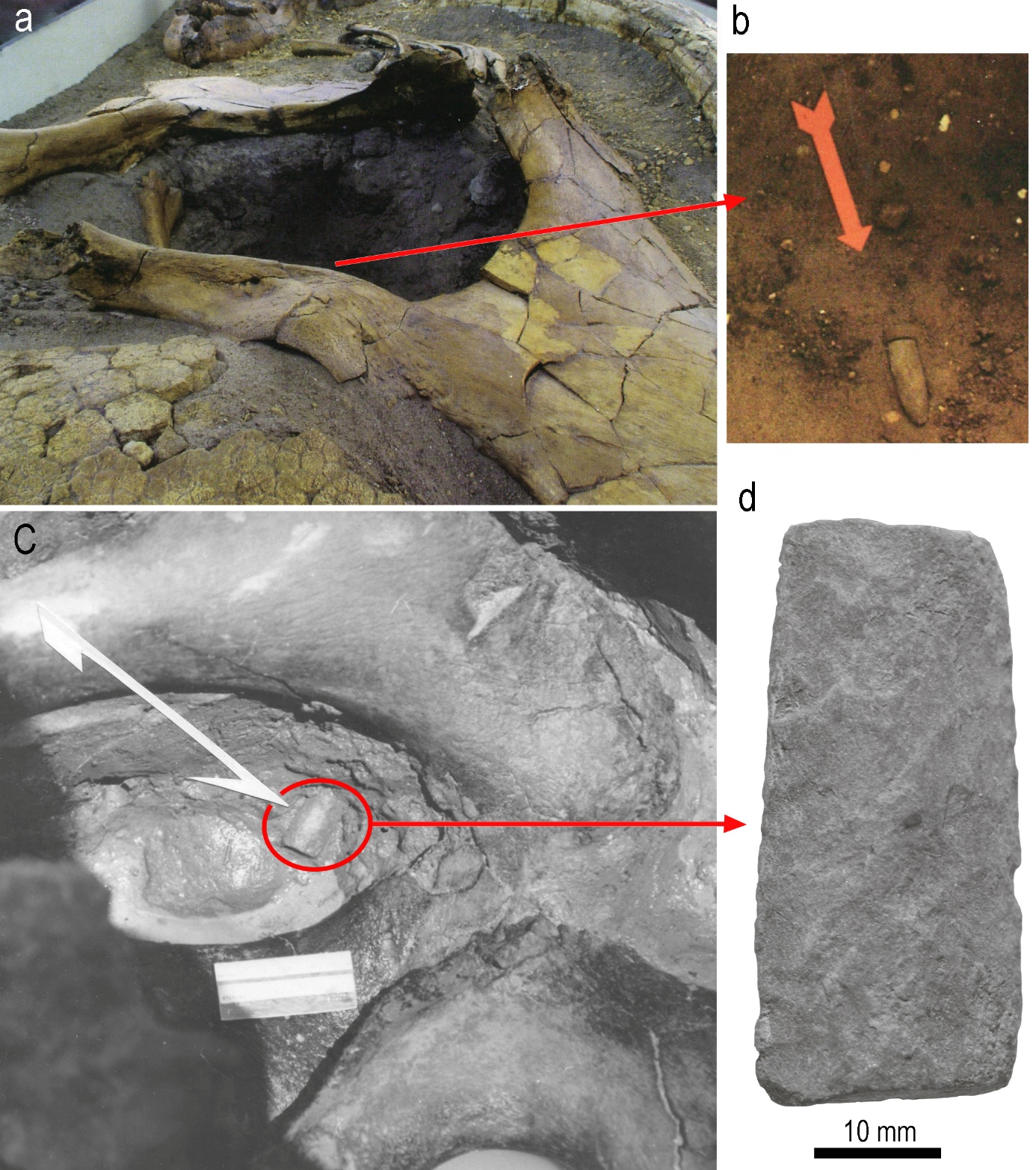

**Figure S1.4**. Pelvic bones of *Notiomastodon platensis* from Taima-Taima and associated with El Jobo projectile fragments from the excavation of 1974 (a, b) and 1976 (c, d; field n° 211/1), respectively. Images a and b modified after Aguilera (2006); image c modified after Cruxent (1978), and image d courtesy archive of the Centro de Antropología del Instituto Venezolano de Investigaciones Científicas (IVIC).

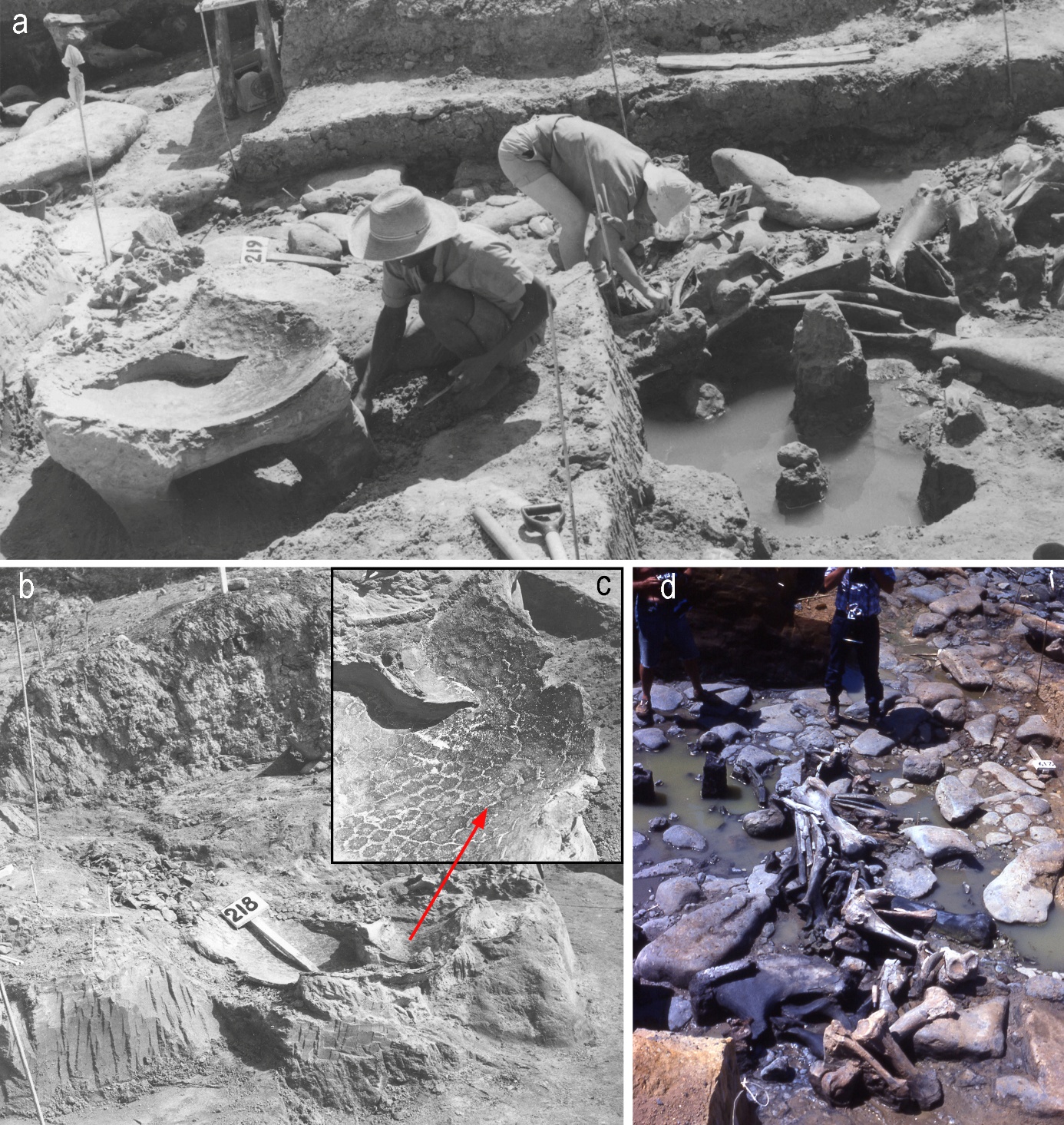

**Figure S1.5**. Excavations at the Taima-Taima site during the third campaign in 1976 (a–d). Second inverted glyptodont carapace (*Glyptotherium* cf. *G*. *cylindricum*) (a–c). In image a, Ruth Gruhn during the process of removing sediment from the semi-articulated skeleton of *Notiomastodon platensis* (see also image d), in whose remains a fragment of an El Jobo-type projectile point was found in situ inside the pelvic cavity (see Fig. S1.4). Images a–c modified after Cruxent (1978); image d courtesy archive of the Universidad Experimental Francisco de Miranda (UNEFF).

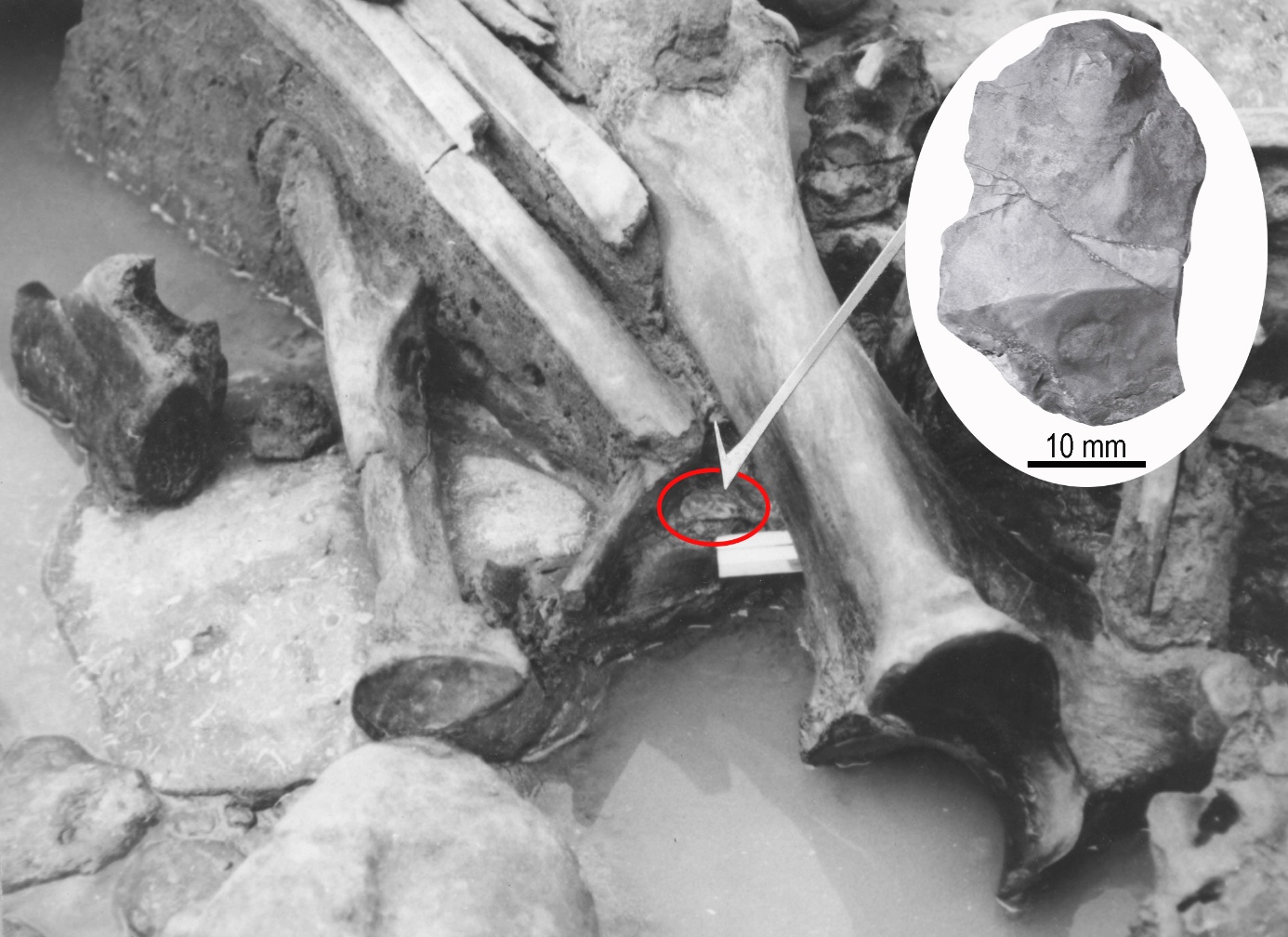

**Figure S1.6**. The white arrow shows a chert scraper (field n° 211/2), produced in chert and *in situ*, between the left ulna and a rib of *Notiomastodon platensis* from the excavation of 1976 (see Cruxent 1978, 1979). Image modified after Cruxent (1978).

After the excavation of 1976, there was a period of inactivity in the Taima-Taima site. It was not until the late 1980s when JM Cruxent resumed excavations, especially in the southern section of the area excavated in 1976. However, the results of this excavation season are unknown (Oliver and Alexander, 2003). Subsequent excavations between 1994 and 1996, led by the Universidad Nacional Experimental Francisco de Miranda (UNEFM) and the Instituto del Patrimonio Cultural de Venezuela (IPC), were carried out to condition the area for the construction of the *in situ* archaeological and paleontological park of Taima-Taima that opened in 2005 (Fig. S1.7). National institutions that with paleontological collections from the Taima-Taima site include: the Centro de Investigaciones Antropológicas, Arqueológicas y Paleontológicas (CIAAP) of the Universidad Experimental Francisco de Miranda (UNEFM), and Museo Comunitario de Taratara “Cristóbal Higuera” (MCH-Pv-) in Falcón State, Laboratorio de Arqueología del IVIC, Miranda State, and the Museo de Ciencias Naturales de
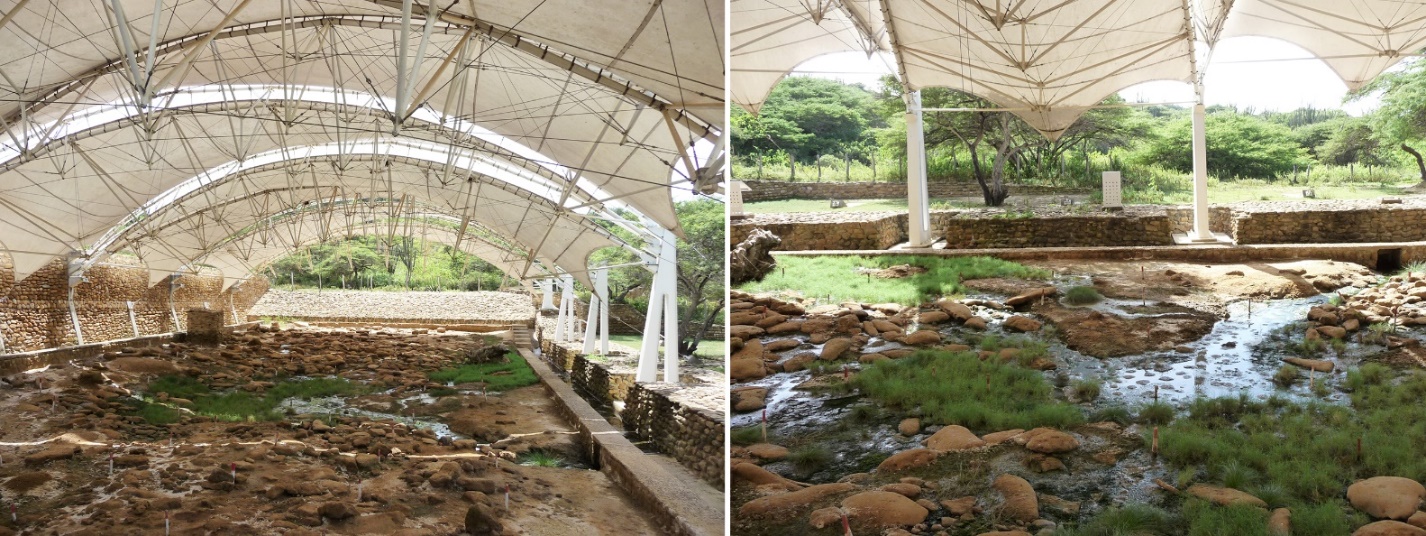
Caracas (MCNC), all in Venezuela.

**Figure S1.7**. *In situ*, the archaeological and palaeontological park of Taima-Taima. In both images, it is possible to observe what corresponds to the Basal stratum of the site, and how the spring remains active at present. Images from Jorge Carrillo-Briceño.

**2.** **Bone/dentine collagen dating**

Twenty-nine bone/dentine samples from Taima-Taima were sent to the Oxford Radiocarbon Accelerator Unit (ORAU) in 2023 for radiocarbon dating and were analysed by one of the authors (Lorena Becerra-Valdivia). These specimens were collected at national collections (Table S1.2) and transported with authorization from the Instituto del Patrimonio Cultural de Venezuela (IPC) using permissions: VE-IPC-CEBC-PP-06/2022-1 and VE-IPC-CEBC-PP-01/2023.

**Table S1.2**. Fossil specimens from Taima-Taima that were sent to the Oxford Radiocarbon Accelerator Unit (ORAU) in 2023. Abbreviations: frag., fragment; m, lower molar; M, upper molar.

| **Sample ID** | **Weight (mg)** | **Sample collection-catalogue number** | **Sample type** | | **Bone element/s** | **Taxonomy** | **Collection** |
| --- | --- | --- | --- | --- | --- | --- | --- |
|  |  |  | **Bone** | **Tooth** |  |  |  |
| **Taima-Ta-01** | **2.04** | CIAAP-91 | X |  | Humerus | *Notiomastodon* sp. | UNEFM-CIAAP |
| **Taima-Ta-02** | **2.00** | CIAAP-58-307 | X |  | Femur | *Notiomastodon* sp. | UNEFM-CIAAP |
| **Taima-Ta-03** | **2.04** | CIAAP-46 | X |  | Humerus | *Notiomastodon* sp. | UNEFM-CIAAP |
| **Taima-Ta-04** | **1.94** | CIAAP-218 | X |  | Ilium frag. | *Notiomastodon* sp. | UNEFM-CIAAP |
| **Taima-Ta-05** | **2.04** | CIAPP-164 | X |  | Rib | *Notiomastodon* sp. | UNEFM-CIAAP |
| **Taima-Ta-06** | **2.02** | CIAPP-86 or 89 | X |  | Rib | *Notiomastodon* sp. | UNEFM-CIAAP |
| **Taima-Ta-07** | **1.93** | CIAAP- s/n° | X |  | Rib | *Notiomastodon* sp. | UNEFM-CIAAP |
| **Taima-Ta-08** | **2.14** | CIAAP-309 | X |  | Femur | *Notiomastodon* sp. | UNEFM-CIAAP |
| **Taima-Ta-09-1** | **1.95** | CIAAP-19-233 | X |  | Ilium frag. | *Notiomastodon* sp. | UNEFM-CIAAP |
| **Taima-Ta-09-2** | **2.11** | CIAAP-19-233 | X |  | Ilium frag. | *Notiomastodon* sp. | UNEFM-CIAAP |
| **Taima-Ta-10-1** | **2.00** | IVIC-AP-020 | X |  | Humerus | *Eremotherium* sp. | IVIC-Arqueología |
| **Taima-Ta-10-2** | **2.05** | IVIC-AP-020 | X |  | Humerus | *Eremotherium* sp. | IVIC-Arqueología |
| **Taima-Ta-10-3** | **2.04** | IVIC-AP-020 | X |  | Humerus | *Eremotherium* sp. | IVIC-Arqueología |
| **Taima-Ta-11-1** | **-** | IVIC-AP-023 | X |  | Tibia | *Eremotherium* sp. | IVIC-Arqueología |
| **Taima-Ta-11-2** | **-** | IVIC-AP-023 | X |  | Tibia | *Eremotherium* sp. | IVIC-Arqueología |
| **Taima-Ta-12** | **1.96** | IVIC-AP-? |  | X | Dentine-tusk | *Notiomastodon* sp. | IVIC-Arqueología |
| **Taima-Ta-13** | **1.93** | CIAAP-1482 |  | X | M 3-dentine | *Notiomastodon* sp. | UNEFM-CIAAP |
| **Taima-Ta-14** | **2.05** | CIAAP-1481 |  | X | M2-dentine | *Notiomastodon* sp. | UNEFM-CIAAP |
| **Taima-Ta-15** | **2.04** | CIAAP-1483 |  | X | m3-dentine | *Notiomastodon* sp. | UNEFM-CIAAP |
| **Taima-Ta-16** | **2.05** | CIAAP-1485 |  | X | M3-dentine | *Notiomastodon* sp. | UNEFM-CIAAP |
| **Taima-Ta-17** | **2.05** | CIAAP-67 |  | X | M2-dentine | *Notiomastodon* sp. | UNEFM-CIAAP |
| **Taima-Ta-18** | **2.06** | CIAAP-83 |  | X | Molar-dentine | *Notiomastodon* sp. | UNEFM-CIAAP |
| **Taima-Ta-19** | **1.97** | MCNC-Pal-1838 | X |  | Skull bone frag. | *Glyptotherium* sp. | MCNC |
| **Taima-Ta-20** | **2.01** | MCNC-Pal-1839 | X |  | Skull bone frag. | *Glyptotherium* sp. | MCNC |
| **Taima-Ta-21** | **1.95** | MCNC-Pal-1839 | | X | Tooth-dentine | *Glyptotherium* sp. | MCNC |
| **Taima-Ta-22** | **2.01** | MCNC-Pal-s/n° | X |  | Rib | Indet. Mammal | MCNC |
| **Taima-Ta-23** | **2.05** | MCNC-Pal-s/n° | X |  | Rib | Indet. Mammal | MCNC |
| **Taima-Ta-24** | **1.94** | MCNC-Pal-s/n° | X |  | Osteoderm | *Glyptotherium* sp. | MCNC |
| **Taima-Ta-25** | **2.03** | MCNC-Pal-s/n° | X |  | Osteoderm | *Glyptotherium* sp. | MCNC |
