## Supplementary material for "Diverse megamammals exploited by humans, chronology and palaeoecology at Taima-Taima, Late Pleistocene, South America": Electronic supplementary material S3

**Notes on taxonomic assessment**

**1. Taxa from the Taima-Taima site whose fossil remains present evidence of modifications of anthropic origin.**

**XENARTHRA (Pilosa)**

†Megatheriidae

†Mylodontidae

*Glossotherium* cf. *robustum*

*Specimen*. Left humerus (IVIC-AP-035)

*Remarks*. Several features support the identification to the genus *Glossotherium*, notably the size, the shape of the groove separating the two proximal tuberosities, the absence of the entepicondylar foramen, a deltopectoral plate oriented anterolaterally, and lesser tuberosity less protruded than the greater tuberosity (Boscaini et al., 2022; De Iuliis et al., 2017; McAfee, 2007). Among related species within the genus, doubt remains between *Glossotherium robustum* and *Glossotherium phoenesis*. The specimen appears relatively gracile like *G. phoenesis* but lacks the stronger projection of the greater tubercle proximally and the stronger distal extension of the ectepicondyle and entepicondyle (Cartell et al., 2019). The strong deltopectoral crest and medial curvature of the diaphyseal margin invalidate *G. tropicorum* (De Iuliis et al., 2017). Identification for *G*. cf. *robustum* is therefore strongly favoured.

Mylodontidae indet.

*Specimen*. Phalange II (MCNC-Pal-1831).

*Remarks*. The overall shape, joint regions and size are reminiscent of an intermediate phalange II of a ground sloth. However, almost no more diagnostic elements are known for this part of the skeleton of this clade. The most precise identification would correspond likely to a phalange II of a digit II or IV of a mylodont (e.g., Haro et al., 2016; Püschel et al., 2017). However, in the absence of more certain arguments, we prefer to propose a cautious and open identification. Consequently, for now, we assign this phalanx to Mylodontidae Indet.

**XENARTHRA (Cingulata)**

†Glyptodontidae

*Glyptotherium* cf. *G. cylindricum*

*Specimens*. Skulls and carapaces reported for the Taima-Taima site (Carlini et al., 2022).

*Remarks*. The remains known from Falcón state (western Venezuela) are practically all attributed to *Glyptotherium*, in accordance with a recent revision (Zurita et al., 2018). The skulls, with fracture patterns interpreted as possible intentional percussion blows and two inverted carapaces (see electronic supplementary material S1) discovered at Taima-Taima were identified as *Glyptotherium* cf. *G*. *cylindricum* by Carlini et al. (2022). To date, no lithic artifacts have been reported in direct association with the remains of glyptodonts in Taima-Taima, which makes it difficult to infer the technique and tools used to hunt these armadillos, especially with blows to the head. Heavy artifacts, such as hafted rocks, could probably have served this purpose (Carlini et al., 2022), and a potential tool (Cruxent, 1967, 1979) was found very close to one of the inverted carapaces in Taima-Taima (electronic supplementary material S1). The inverted carapaces, skulls and other postcranial bones of Glyptotherium cf. G. cylindricum were reported for the Unit I/II erosional disconformity of the Upper stratum (Casamiquela, 1979; Carlini et al., 2022). Only a few isolated osteoderms (Casamiquela 1979) and the humerus are coming from the Medium stratum.

**†LITOPTERNA**

†Macraucheniidae

cf. *Xenorhinotherium bahiense*

*Specimen*s. A fragmentary limb bone represented by a right fused ulna-radius fragment (MCNC-Pal-1834).

*Remarks*. One of the problems presented by the Taima-Taima specimens studied here is their fragmentary state of preservation. Nevertheless, MCNC-Pal-1834 may belong to *Xenorhinotherium bahiense*, the only macraucheniid so far identified (based on cranial and dental elements) for the Taima-Taima and Muaco sites (Aguilera, 2006). The fossil record of *X*. *bahiense* suggests that this was the only macrauchenid species that inhabited northern South America at the end of the Pleistocene (Scherer et al., 2009), although disagreements exist over the validity of this taxon, and some authors consider it synonymous with *Macrauchenia patachonica* (Guerin and Faure, 2004). We tentatively assign the materials under study from Taima-Taima to cf. *Xenorhinotherium bahiense*.

**†NOTOUNGULATA**

†Toxodontidae

Toxodontidae indet.

*Specimen*. Left ulna (CIAAP-1533).

*Remarks*. This ulna is the only long bone so far recognized as a toxodontid from the Taima-Taima site. The incomplete fusion between the metaphysis and distal epiphysis suggests that this is likely to belong to a subadult individual. The only toxodontid reported for the Late Pleistocene of Venezuela (Rincón, 2011), and other localities in northern South America and Central America, is *Mixotoxodon larensis*, and the determinations of these specimens are based mainly on dental elements (Bocquentin-Villanueva, 1979; Hernández Jasso and Blanco Piñon, 2020). Aguilera (2006) referred to the isolated dental elements and a foot bone from Taima-Taima as *Mixotoxodon* cf. *larensis*. Dental elements assigned to *Mixotoxodon larensis* have also been reported from the nearby site of Muaco (Bocquentin-Villanueva, 1979; Rincón, 2011). It is conceivable that the ulna CIAAP-1533 may also belong to *M*. *larensis*. However, we remain prudent and keep the specimen identified for now as Toxodontidae indet., since postcranial elements of *Mixotoxodon* have been little studied and their diagnosis is unknown.

**PROBOSCIDEA**

†Gomphotheriidae

*Notiomastodon platensis*

*Specimen*. Eight bone elements are reported here with evidence of modifications of anthropic origin (electronic supplementary material S2). However, the number of cranial and postcranial elements from the Taima-Taima site and deposited in different collections in Venezuela exceeds more than one hundred specimens.

*Remarks*. Two Proboscidea species, *Notiomastodon platensis* and *Cuvieronius hyodon*, occurred in South America during the Late Pleistocene–Early Holocene (Alberdi and Prado, 2021; Mothé et al., 2017a); most of their diagnostic features come from dental specimens, including upper tusks and last molars, as well as lower jaw symphysis and tusks (Mothé et al., 2016; 2017b). *Notiomastodon* and *Cuvieronius* share a similar lower jaw structure with a short horizontal ramus and reduced symphysis; however, while *Notiomastodon* has no trace of lower tusks, *Cuvieronius* presents a pair of lower incisors or its corresponding vestigial alveoli. *Cuvieronius* has elongated, twisted, and slightly upcurved upper tusks with a longitudinal enamel band, while *Notiomastodon*'s upper tusks have a great variation in length, robustness, shape, and enamel presence, and are never twisted (Mothé and Avilla, 2015). The post-canine teeth of both proboscideans are bunodont and quite similar in morphology, differing only in the complexity of the last molars (number of main and accessory cusps), in which *Notiomastodon* had a range of 35–82 cusps and *Cuvieronius* 33–60 cusps. Considering these morphological traits, the specimens from Taima-Taima, as well as those from the nearby sites of Muaco and Cucuruchú (Carrillo-Briceño, 2015), present features that fit *Notiomastodon platensis* diagnostic features, such as lower jaws with downturned and plain symphysis (i.e., without lower tusks and/or their vestigial alveoli), linear upper tusks (untwisted), which vary from straight to slightly upcurved, and third molars with cusps ranging from at least 38 to 81 (the lower score might be higher due to fragmentation in some last molars). Thus, we confirm that the proboscidean remains from this site belong to the species *Notiomastodon platensis*, taxon that has also been reported in other localities from the Late Pleistocene of the Falcón state (Carrillo-Briceño et al., 2024) and elsewhere in Venezuela (Carrillo-Briceño, 2015).

**2. An approximation of the number of individuals of *Notiomastodon platensis* present in Taima-Taima.**

With the exemption of the semi-articulated skeleton collected in 1976 (Bryan et al., 1978; Casamiquela, 1979), the assignment of other cranial and post-cranial elements (collected in Taima-Taima in different excavation campaigns) to individuals is a difficult task. The latter is mainly due to the absence of detailed stratigraphic information for many of the specimens and the state of disarticulation and fragmentation in most of these. However, using dental elements, their type of tooth and wear stage facilitates the estimation of the “Minimum Number of Individuals (MNI) (see Simpson and Paula-Couto, 1957; Mothé et al., 2010).

According only to the type and wear stage of the post-canine teeth from Taima-Taima (Table S3.1), the MNI corresponds to at least 14 *N*. *platensis* individuals, including 4 immatures (calves), 2 subadults, 3 adults, 2 mature adults, and 3 senile adults (Fig. S3.1). When the same type of tooth with the same wear stage was identified, we paired it with a corresponding tooth in the dental arch, e.g., a left lower last molar in wear stage 3 (CIAAP-1483) with a right lower last molar in wear stage 3 (CIAAP-61-224). However, if we consider the combination of these traits with the taphonomic aspects (e.g., colour, fragmentation, presence of dental calculus) and general morphology of the specimens (e.g., number and complexity of lophs/lophids, robustness of tooth, and morphology of distal cingulum), the number of individuals increases to 23, being 4 immatures (calves), 3 subadults, 6 adults, 3 mature adults, and 7 senile adults. We believe that a minimum number of 14 individuals can be recognized as a conservative number until more detailed studies are carried out in the future.

*Notiomastodon platensis* individuals from Taima-Taima may have reached almost 5 tons in body mass, according to the body mass estimative using adult femur length (Larramendi, 2015). The complete femur CIAAP-111 (corresponding to the semi articulated skeleton of 1976 and its respective mandible and molars CIAAP-83/-1486) might have weighed approximately 2.5 tons, being possibly a female adult from 29 to 35 years (estimated age by associated teeth and wear stage, see Table S3.1), while the individuals represented by the specimens CIAAP-311 and CIAPP 312 might have reached between 4 and 2.3 tons, respectively. Unfortunately, these last specimens were not associated with dental materials; therefore, it is not possible to infer a more precise age class beyond adults.

**Table S3.1**. Type and wear stage of the post-canine teeth of *Notiomastodon platensis* from Taima-Taima. Abbreviations: dp, deciduous premolar; M/m, molar.

| **Specimen number** | **Type of teeth** | **Wear stage** | **Age in years** | **Age class** |
| --- | --- | --- | --- | --- |
| CIAAP-78-226 | Right lower dp2 | Stage 1 | 1 or less | Immature |
| CIAAP-67 | Left lower dp4 | Stage 2 | 6 | Immature |
| CIAAP-84 | Right lower dp3 | Stage 4+ | 6 | Immature |
|  | Right lower dp4 | Stage 2 | 6 |  |
| CIAAP-1480 | Lower left dp3 | Stage 2 | 2-3 | Immature |
| MCNC-Pal-s/n | Upper left dp2 | Stage 2 | 1-2 | Immature |
|  | Upper right dp2 | Stage 2 |  |  |
|  | Upper left dp3 | Stage 1 |  |  |
|  | Upper right dp3 | Stage 1 |  |  |
| IVIC-AP-030 (mandible) | Lower right m1 | Stage 3 | 17-20 | Subadult |
|  | Lower left m1 | Stage 3 |  |  |
|  | Lower right m2 | Stage 1 | 19-24 |  |
|  | Lower left m2 | Stage 1 |  |  |
| CIAAP-223 | Lower right m2 | Stage 0 | 19 | Subadult |
| IVIC-AP-032 | Lower left m2 | Stage 1 | 19-24 | Subadult |
| CIAAP-1479 | Lower right m2 | Stage 2 | 25-29 | Adult |
| CIAAP-1481 | Upper left M2 | Stage 2 | 25-29 | Adult |
| CIAAP-1484 | Lower left m2 | Stage 2 | 25-29 | Adult |
| IVIC-AP-033 | Upper left M2 | Stage 3 | 29-35 | Adult |
| CIAAP-83-1486 | Lower right m2 | Stage 3 | 29-35 | Adult |
| CIAAP-1485 | Lower right m3 | Stage 0 | 28-34 | Adult |
| CIAAP-80-215 | Lower left m2 | Stage 3 | 34-41 | Mature adult |
| IVIC-AO-031 | Lower right m3 | Stage 1 | 37-41 | Mature adult |
| IVIC-AP-034 | Lower right m3 | Stage 1 | 37-41 | Mature adult |
| CIAAP-1483 | Lower left m3 | Stage 3 | 49-53 | Senile adult |
| CIAAP-313 | Lower left m3 | Stage 3 | 49-53 | Senile adult |
| CIAAP-70-215 | Lower right m3 | Stage 3 | 49-53 | Senile adult |
| CIAAP-61-224 | Lower right m3 | Stage 3 | 49-53 | Senile adult |
| CIAAP-12-286 | Lower right m3 | Stage 3 | 49-53 | Senile adult |
| CIAAP-214 | Upper right M3 | Stage 3 | 49-53 | Senile adult |

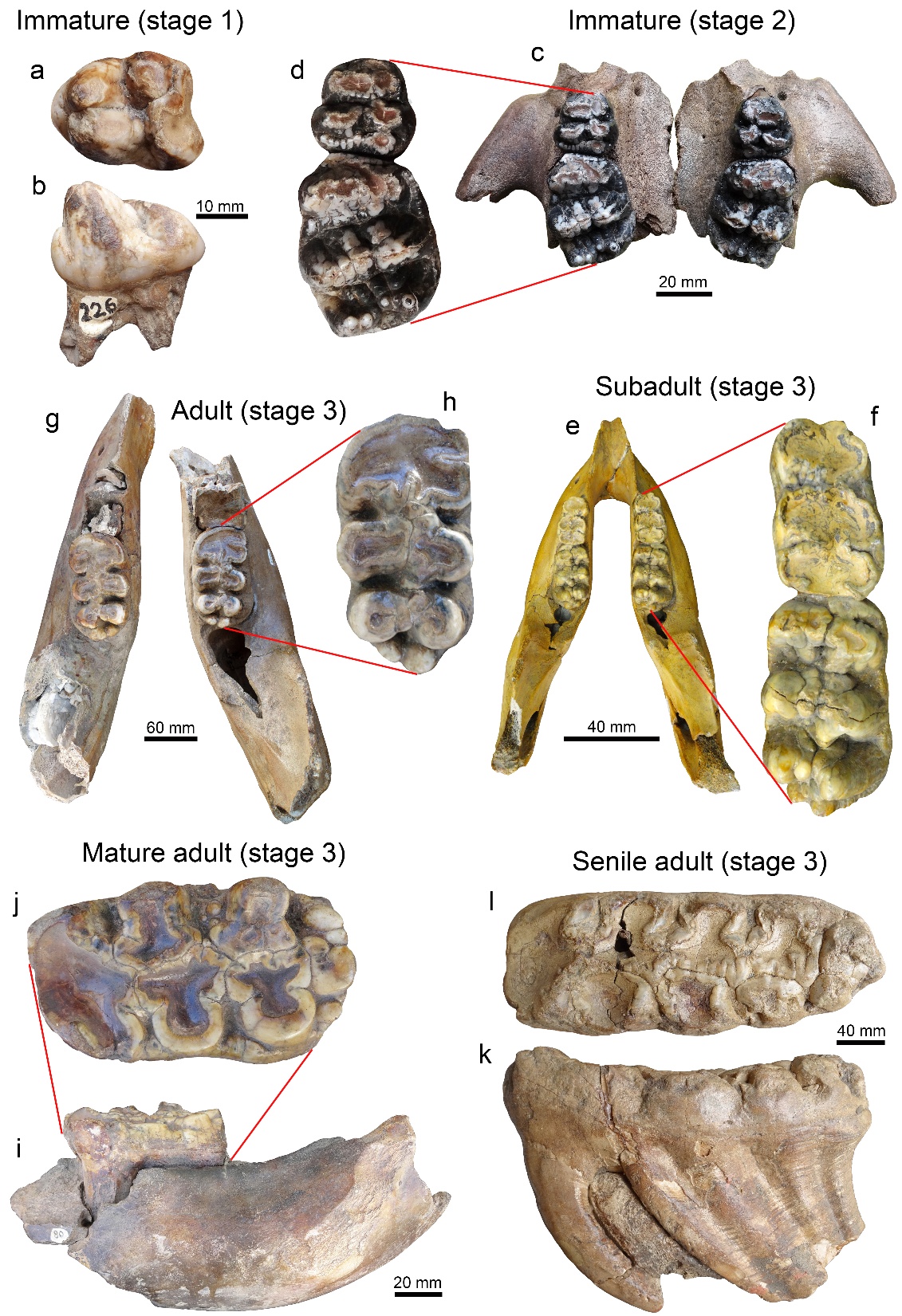

**Figure S3.1**. Examples of age classes and wear stage in some post-canine teeth of *Notiomastodon platensis* from Taima-Taima. Right isolated deciduous premolar 2 (a, b: CIAAP-78-226), maxilla with upper right and left deciduous premolar 2 and 3 (c, d: MCNC-Pal-s/n), mandible with lower right and left molar 1 and 2 (e, f: IVIC-AP-030), mandible with right left molar 1 (broken) and 2 (g, h: CIAAP-83 and -1486), left hemimandible with molar 2 (i, j: CIAAP-80-215), and isolated lower right molar 3 (k, l: CIAAP-61-224).
