## Supplementary material for "Diverse megamammals exploited by humans, chronology and palaeoecology at Taima-Taima, Late Pleistocene, South America": Electronic supplementary material S4

**Surface modification of bones and other evidence of anthropic origin in megaherbivore taxa from the Taima-Taima site (Late Pleistocene)**

Different variables are involved in the processes of bone surface modifications produced by humans, and their type, size/shape, distribution and frequency, is possible to be inferred due to certain patterns (e.g., Binford, 1981; Courtenay et al., 2018; Fernández-Jalvo and Andrews, 2016; Pizarro-Monzon et al., 2021; among others). This type of evidence, which in most cases was produced unintentionally (Pizarro-Monzon et al., 2021), would reflect human behavior associated with butchering activities. Interpreting these unintentionally marks in the present for inferring past human behavior regarding carcass manipulation has always been a challenge. There is an ongoing work which involves experimental archaeology, next-generation technology, taphonomic contextualization and profound academic antecedents to define an anthropic cut-mark. For that, there is always a continual debate to define and differentiate a butchering mark from other causes. Considering these issues, for this work, a matrix for identifying the classical attributes that define anthropic marks (electronic supplementary material 2), considering the information developed below, was performed. In this regard, we present the most conspicuous marks we found in some fossil bones from the Taima-Taima site. Nevertheless, as the research to define the most characteristic cut-marks is in continual renewal, there will always be doubt in the agent behind traces on bone surface.

From the Taima-Taima fossil assemblage, we have preliminarily identified 11 bone remains (cranial and postcranial elements) of at least that four extinct megaherbivores with evidence of probable modifications of human nature. We have categorized these modifications into nine types (Table 1; electronic supplementary material S2), based on different classical morphological attributes and criteria (e.g., Lyman, 1994; Domínguez-Rodrigo et al., 2009; Fernández-Jalvo and Andrews, 2016). The modifications described in this contribution are the following:

*Chop marks*. These are created by sharp wedge-shaped tools that strike the bone with force and are designed to bite into (Okaluk and Greenﬁeld, 2022). These types of marks usually penetrate deeply into the bone with a smooth sloping side showing the direction of the cut, and its origin can only be caused by tool or falling blocks (Fernández-Jalvo and Andrews, 2016). In the Taima-Taima deposit, the presence of falling blocks is ruled out, since the geology and morphology of the terrain do not allow their presence or existence.

*Cut and slicing marks*. Both are produced by linear movements on the surface of the bone. Usually, cutting marks produce a deep V-shaped cross-section (wide or narrow), while slicing marks tend to be less deep, which contrasts with linear marks left by the teeth of carnivores develop a U-shaped cross-section (Lyman, 1994). Cut marks can be expressed with both sides having a symmetrical cross-section, but in some cases an asymmetric cross-section may also be present (Domínguez-Rodrigo et al., 2009). Cut and slicing marks show a marked difference with trampling marks, because former can usually be related to parts of bones where muscle or tendon insertions are present or located at concave protected areas that are not reachable by trampling unless the salient bone angles are damaged; in general, trampling marks are shallower than cuts (see Fernández-Jalvo and Andrews, 2016). According to Lyman (1994), sedimentary abrasion will not produce marks that mimic slicing marks. According to Greenfield (2006), different types of raw materials and tools (e.g., blades, flakes, and side scrapers) used in the butchering process produce slicing cut marks.

*Hertzian cones*. These are of triangular morphology and are caused by the stress produced on the bone by the differential pressure exerted on the surface due to the bone resistance when cutting with a tool (Bromage and Boyde, 1984; Fernández-Jalvo and Andrews, 2016).

*Internal microsteps*. In the form of steps are produced during the movement of the tool and are characterized by break up the smooth outline of the cut mark (Fernández-Jalvo and Andrews, 2016).

*Internal micro-strations*. They are attributed to the movement of rocks on the bone produced by their displacement (e.g., mechanical action of trampling, transport) or by retouched edges of lithic instruments (Shipman and Rose, 1983).

*Shoulder effect*. These are pseudo cut marks and parallel to the primary cut, and they are produced by irregularities of the stone cutting edge (Fernández-Jalvo and Andrews, 2016).

*Scraping marks*. These can be very long, straight and produced by holding the stone tool edge transversally to the direction of motion (Andrews and Fernández-Jalvo, 2012).

*Percussion marks*. These can be notches or depressions on the bone surface, of variable sizes and depths, produced when the bone smashed with a stone or other hard artifact (Fernández-Jalvo and Andrews, 2016). Bone fractures can also be caused by percussion agents.

**1. Modified bones from Taima-Taima**

1.1 Mylodontids

A humerus and a phalange of likely different individuals with evidence of probable anthropic marks were identified. The left humerus (IVIC-AP-035), assigned here to an adult individual of *Glossotherium robustum* (electronic supplementary material S3), is complete and well-preserved (Fig. S4.1), exhibiting abundant cracks and some manganese spots (e.g., Fig. S4.1c–e). We preliminarily assume that these are not weathering cracks, and they, likely were produced in this limb bone post-excavation, probably generated by changes from a humid and water-saturated environment to a drier one (e.g., collection deposits). These cracks are common in many of all the materials we have studied from Taima-Taima, and they do not present a sediment filling, which justifies our hypothesis. Even though some of these cracks seem to have occurred after the cleaning and restoration of the fossil specimens, since older layers of consolidants were also affected. Future taphonomic work may shed more light on the origin of these cracks.

In posterior view, IVIC-AP-035 shows a set of linear cut marks (Fig. S4.1f–h, S4.2) on the capitulum. The largest of these marks, with about 12 mm long, correspond to a chop mark, which is deep and has a typical “symmetric” V-shape (Fig. S4.1j, S4.2b). Well-defined hertzian cones are preserved on both sides of the mark (Fig. S4.1j), and what appears to be an internal micro-striation is vaguely observed. A second mark appears parallel to the main mark with a separation of 14 mm between them (Fig. S4.1i). This mark is smaller, about seven millimeters long, it is less deep, it seems to be somewhat asymmetrical in cross-sections, and it is also characterized by the presence of hertzian cones (Fig. S4.1i). A set of a little more than 14 shallow and parallel marks are observed diagonally to the main mark (Fig. S4.1g, h). These marks are up to about 8 or 9 millimeters long, and due to their parallel condition, they may correspond to potential slicing marks. No other relevant surface modifications are observed in IVIC-AP-035, which suggests that the marks described here can be related to human action and not to other pre/post-burial agents. Their location in the distal epiphysis could be associated with the cutting and dismembering process of the forearm.

The intermediate phalanx II (MCNC-Pal-1831) of an indeterminate mylodontid (electronic supplementary material S3) presents a set of three well-defined short and deep marks on the ventro-lateral left portion (Fig. S4.3b), the longest being about 8 mm long. These marks are very close to each other, two of which are parallel, and one is diagonal. A probable fourth mark, very small of only two millimeters long, is located close to and parallel to the diagonal mark mentioned above. The three main marks are characterized by deep V-shaped, and they seem to have an asymmetrical profile. In the middle one, internal striations and what appear to be internal microsteps can be observed. These characteristics and the location and orientation of the marks indicate an anthropic origin, likely a chop mark. The phalanx is covered by manganese, including the marks, confirming they were performed before the bone blackening. A long crack appears to run through the marks and other parts of the phalanx, and this is filled in (Fig. S4.4), and must have occurred subsequently to the modifications of anthropic origin.

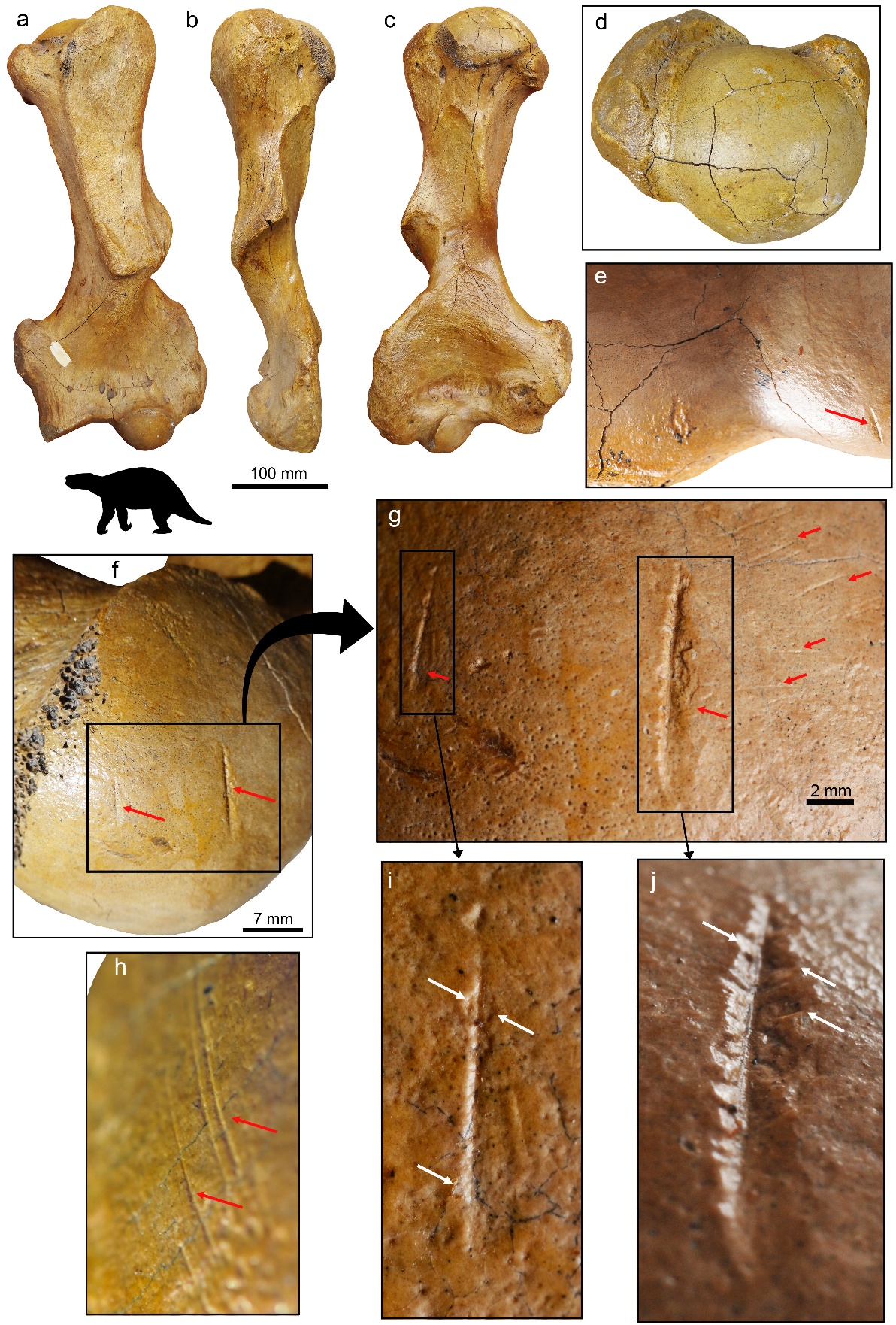

**Figure S4.1**. Left humerus (a–c: IVIC-AP-035) of *Glossotherium robustum* from Taima-Taima with a series of linear cut marks on the capitulum (f–j) cracks are observed in figures d and e. Red arrows show the marks, and white arrows hertzian cones. Views: anterior (a), lateral (b), posterior (c), proximal (d).

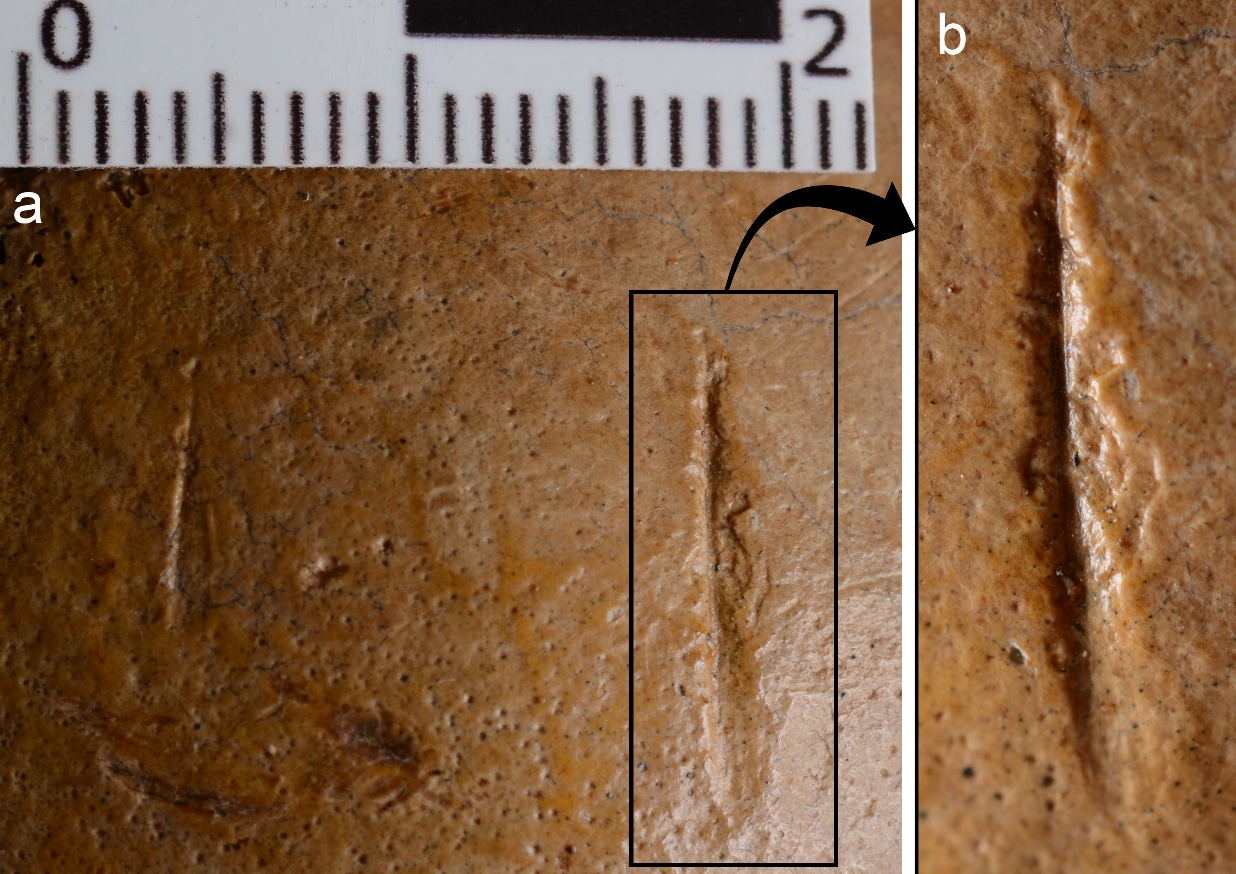

**Figure S4.2**. Detail of the linear cut marks on the capitulum of the left humerus (IVIC-AP-035) of *Glossotherium robustum* from Taima-Taima. The longest (b) is about 12 mm long.

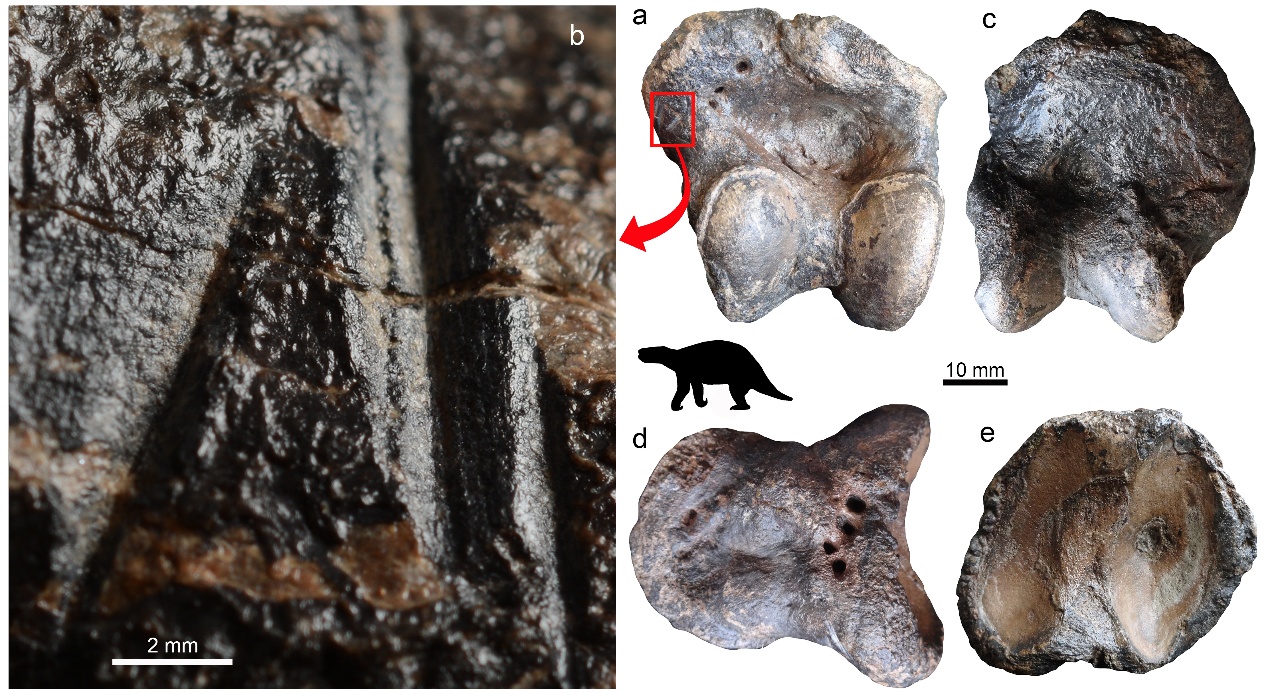

**Figure S4.3**. Indeterminate Mylodontidae phalange II (a, c–e: MCNC-Pal-1831) from Taima-Taima with V-shaped cutting marks (*b*). Views: dorsal (c), lateral (d), proximal (e), ventral (a).

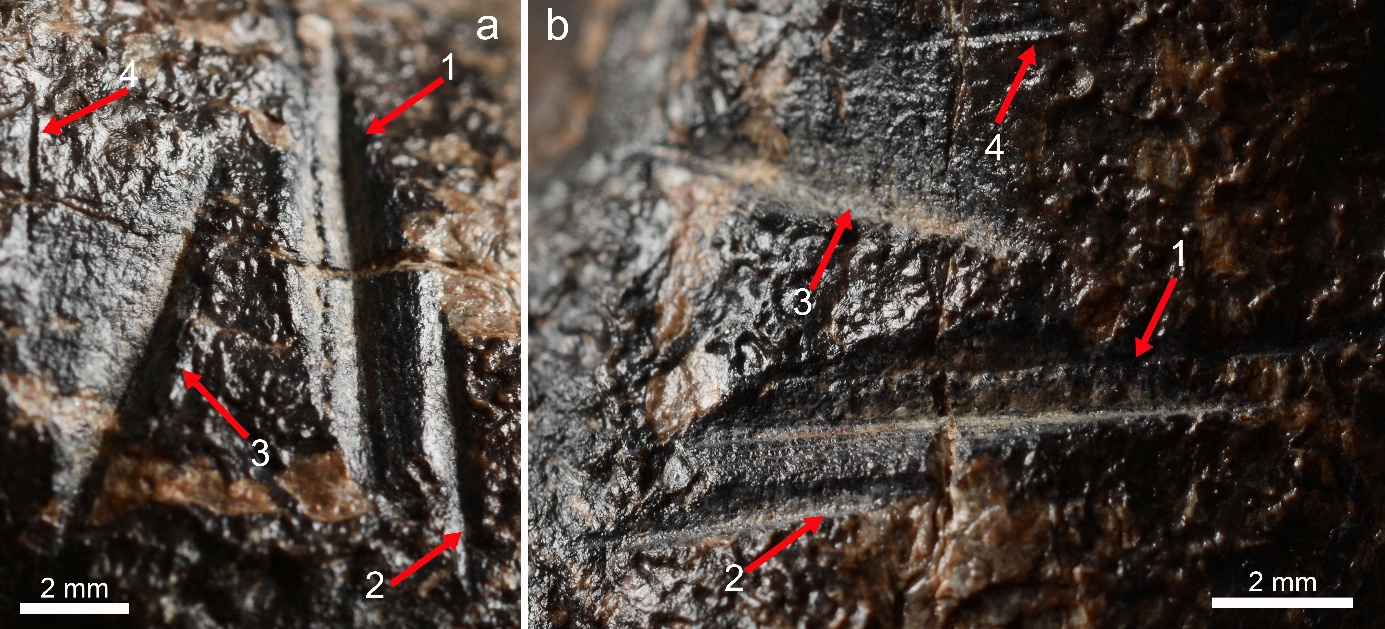

**Figure S4.4**. Detail of the linear V-shaped cut marks (in different orientation: a, b) on indeterminate Mylodontidae phalange II (MCNC-Pal-1831) from Taima-Taima. Red arrows show the marks.

1.2 Macraucheniids

We have identified a modified bone fragment of cf. *Xenorhinotherium* *bahiense* (electronic supplementary material S3), represented by a fragmentary and isolated right fused ulna-radius (MCNC-Pal-1834). Specimen MCNC-Pal-1834 corresponds to the diaphysis of a right fused ulna-radius, preserving part of the aliform expansion of the radius, and missing both epiphyses (Fig. S4.5a, c, d). The bone is preserved in dark brown color, and some sediment spots and bone cracks were observed. We believe that these cracks probably correspond to a desiccation cracking that occurred after the extraction of the fossil from the deposit. In the distal section of the diaphysis, on the anterior face, in a transverse position, a well-defined and deep V-shaped cut mark was identified (Fig. S4.5b, e). This mark has an approximate length of about 8 millimeters, it has a symmetrical cross-section, and on both sides preserves shoulder effects (Fig. S4.5e).

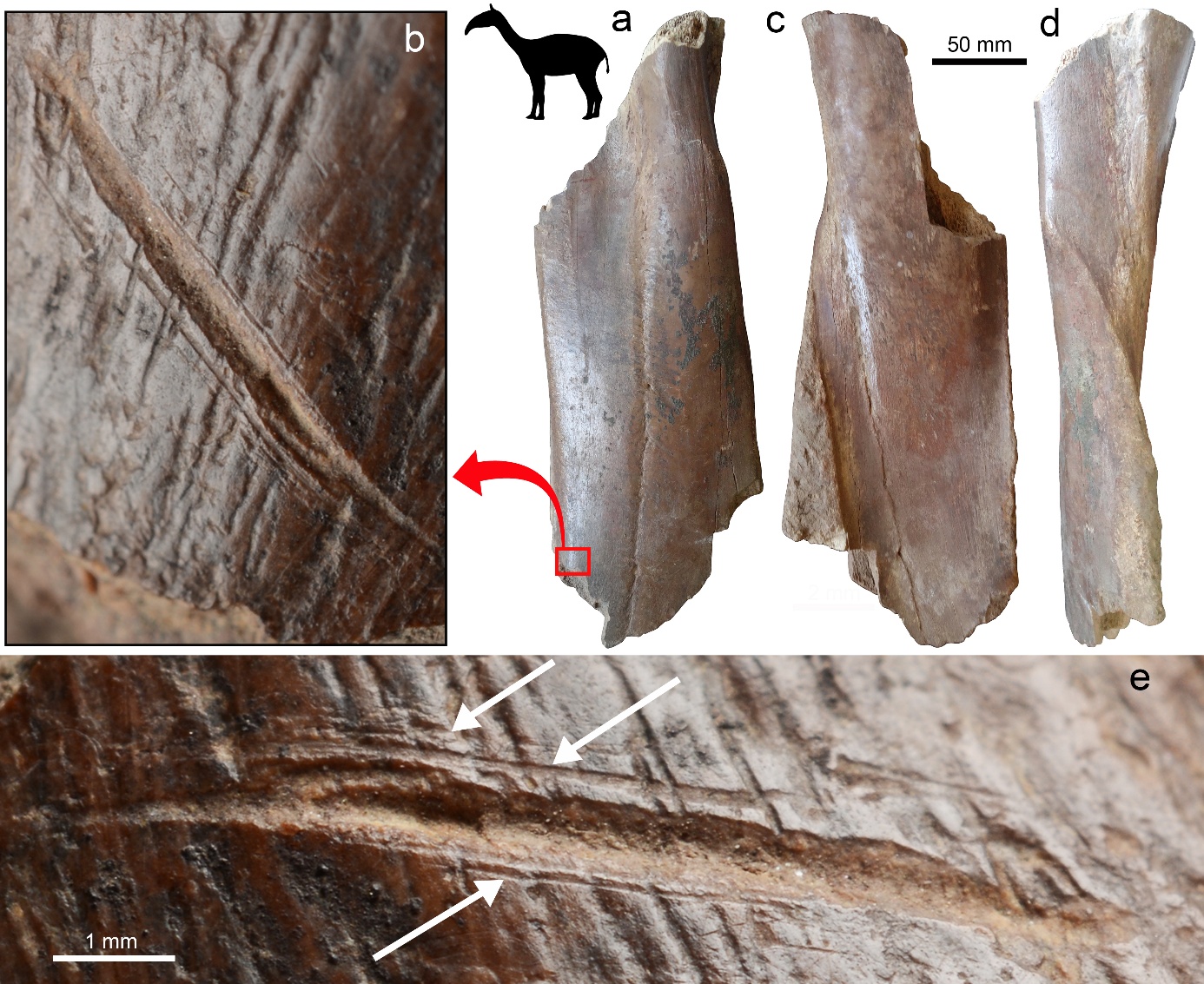

**Figure S4.5**. Fragmentary right fused ulna-radius of cf. *Xenorhinotherium* *bahiense* (MCNC-Pal-1834), in anterior (a), posterior (c) and lateral (d) views showing a cutting mark (b, e). White arrows show shoulder effects.

1.3 Toxodontids

An isolated and complete left ulna (Fig. S4.6) of an indeterminate toxodontid (CIAPP-1533; probably *Mixotoxodon*, electronic supplementary material S3) also shows modifications of probable anthropic origin (Figs. S4.6–8). This ulna likely belonged to a subadult, since metaphysis and distal epiphysis are not completely fused. The cortical surface is well preserved, with some manganese staining and cracks, the latter produced probably by desiccation after the extraction of the fossil from the deposit (Fig. S4.9). A probable cut mark was identified, being transverse and located on the lateral side of the diaphysis in the coronoid process (Fig. S4.6a, c). This one is about 11 mm long, not very deep, has a slightly V-shaped and seems symmetrical in cross-section. On one of its edges, what appears to be hertzian cones are preserved. Unfortunately, this part of the bone is thickly covered with lacquer, which prevents clearer details from being observed. In the articular facet of the trochlear notch, near the lateral edge, a set of at least 24 shallow relatively parallel marks are observed (Fig. S4.7, 8). The longest of these marks reaches 22 millimeters in length. Due to their parallel condition, they may correspond likely to slicing marks, probably produced during the dismemberment and separation of the ulna and humerus. It is unlikely that the articular facet of the trochlear notch, due to its morphology and position, has been affected by trampling or another erosive process. This latter aspect is supported because no other region of the cortical surface of the ulna preserves erosive and trampling marks. We identified the presence of some recent modifications, especially caused by damage during handling of the fossil, and these were observed in the articular facet of the trochlear notch. These modifications look fresh (Fig. S4.9b) in comparison with the cut marks, they even seem to have been produced by friction with other specimens or harder materials. Loss of cortical material is observable on the lateral side of the ulna between metaphysis and distal epiphysis (Fig. S4.6d). However, we do not have evidence that the latter was generated intentionally, but it seems to have been produced before burial. No other evidence which we think could be of human origin was identified.

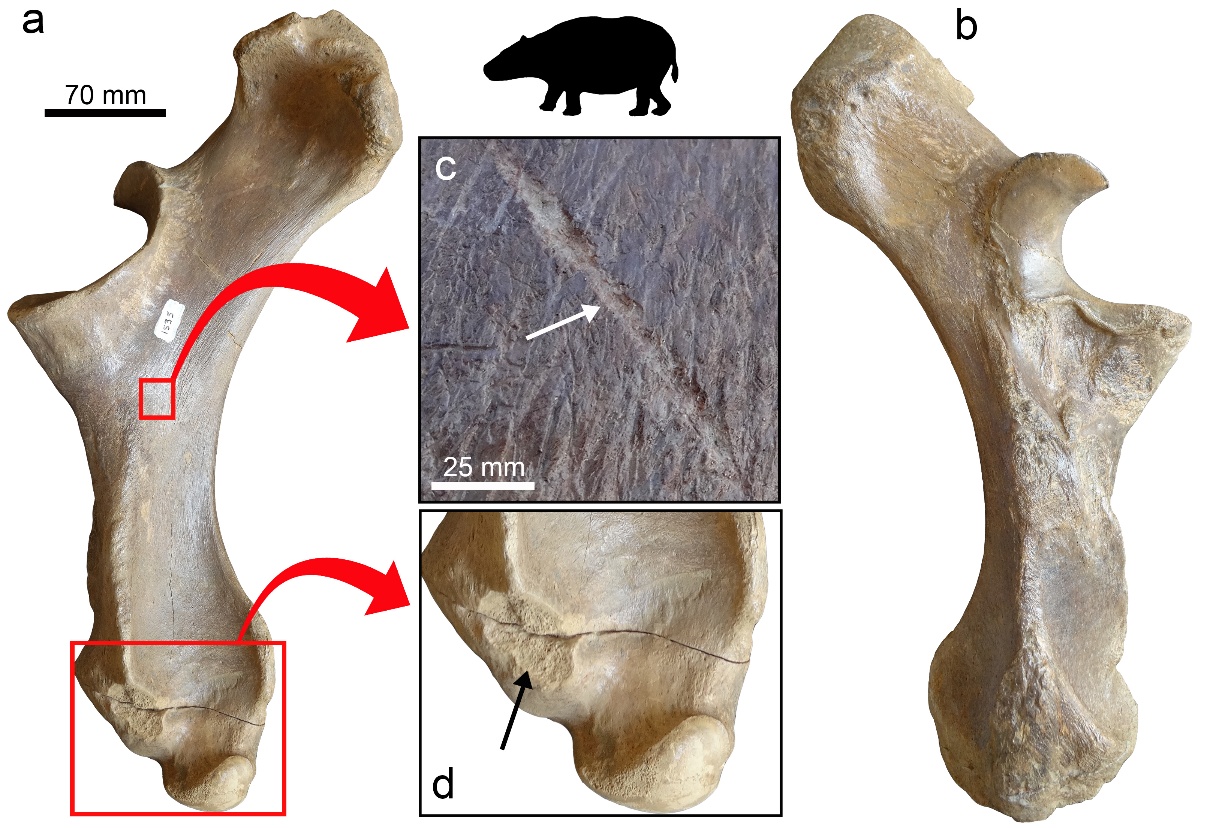

**Figure S4.6**. Lef ulna (CIAAP-1533) in lateral views (a, b) of an indeterminate toxodontid (probably *Mixotoxodon*) from Taima-Taima. Cut mark (b) and loss of cortical material on the lateral side of indeterminate origin nature (d)

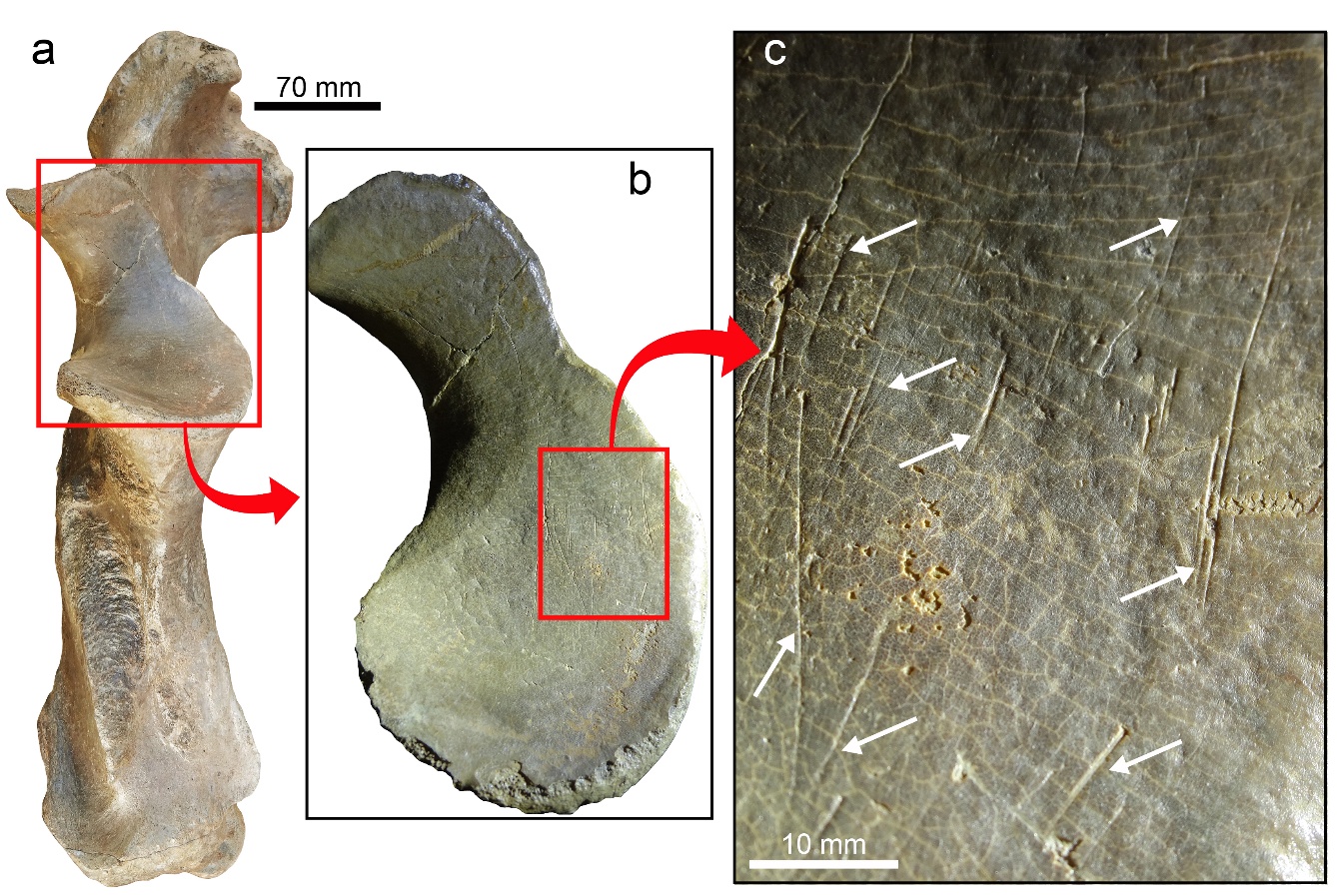

**Figure S4.7**. Lef ulna CIAAP-1533 of an indeterminate toxodontid (probably Mixotoxodon) from Taima-Taima in anterior (a) and proximal (b) views, showing a set of slicing marks (white arrows) on the articular facet of the trochlear notch (c).

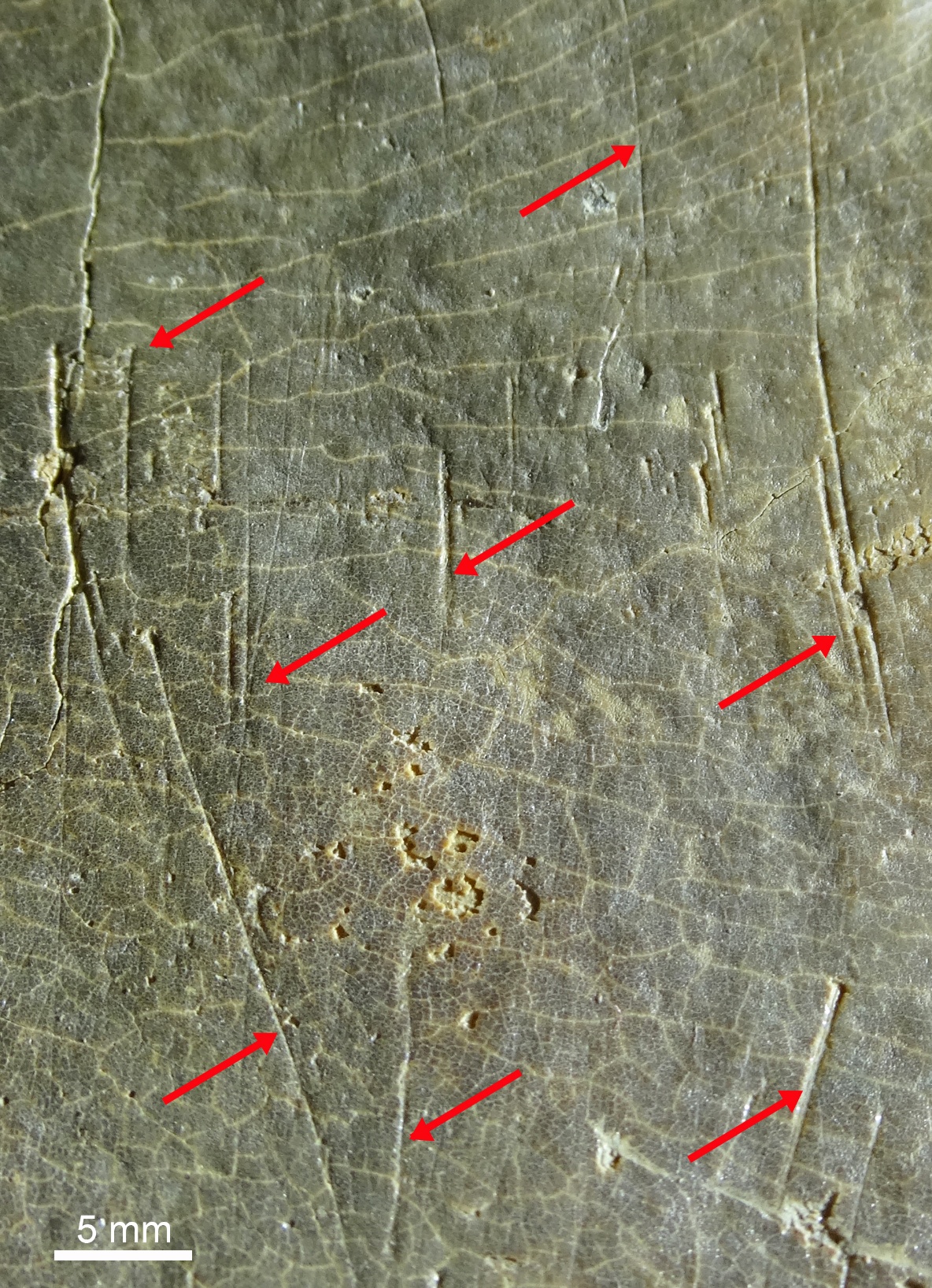

**Figure S4.8**. Set of slicing marks (red arrows) on the articular facet of the trochlear notch of the left ulna CIAAP-1533 of an indeterminate toxodontid (probably Mixotoxodon) from Taima-Taima.

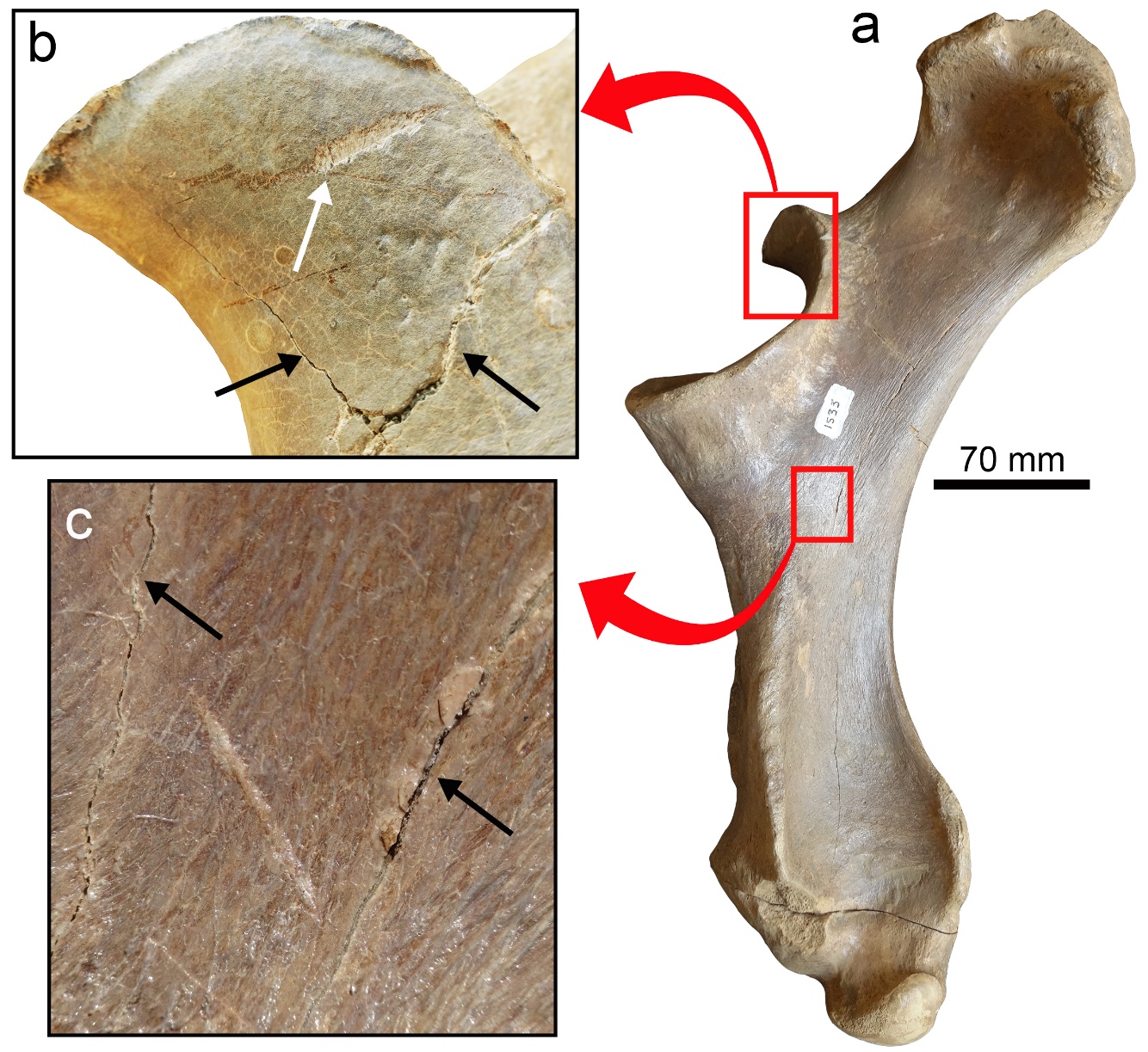

**Figure S4.9**. Lef ulna CIAAP-1533 of an indeterminate toxodontid (probably Mixotoxodon) from Taima-Taima in anterior in lateral view (a), showing some examples of cracks (black arrows) and recent (post-excavation) damage (white arrow) caused by handling the fossil (b).

1.4 Gomphotheres

The remains of *Notiomastodon platensis* are the most abundant at the Taima-Taima site (electronic supplementary material 3). Several bones were associated with the carcass of an individual excavated in 1976, in a semi-articulated condition, and with an El Jobo projectile point in its pelvic region (Bryan et al., 1978; Casamiquela, 1979), as well as with many other isolated elements collected during different campaigns. The elements that form part of the skeleton or carcass of the1976 do not have a single catalogue number. On the contrary, almost all the elements referred to this taxon have a particular number previously assigned in the collection where they are deposited (see electronic supplementary material S2).

A detailed anatomical and taphonomic description of the bone elements of the 1976 skeleton, and other findings was presented by Casamiquela (1979). This author recognised modifications of anthropic origin (*i.e.*, cut marks) only in the left humerus (CIAAP-46) and two ribs (CIAAP-43-154 and CIAAP-145) of the above-mentioned skeleton. Our documentation confirms the presence of at least six well-defined V-shaped cut marks in humerus CIAAP-46, located on the anterior surface above the base of the humeral crest (Fig. S4.10b, c). These marks are parallels between them, short, with lengths not exceeding 14 mm for the longest of the group, and at least five of them preserve well-defined hertzian cones (e.g., Fig. S4.10d). An apparent slicing mark of about 50 mm seems to be the continuation of the mark that is in the lowest position of the group (Fig. S4.10c). We have also identified other modifications of anthropic origin in elements of the 1976 skeleton that were not previously reported, including a left scapula (CIAPP-47-229), a right femur (CIAAP-111), and a right tibia (CIAAP-109) (Fig. S4.11, 12). A series of at least three linear marks, which we think could be associated with slicing marks, were observed on both sides of the scapula CIAPP-47-229 (Fig. S4.12c). A circular depression was also observed in the scapula, with an approximate diameter of 38 mm in its widest part (Fig. S4.12b). Lack of collapsed cortical bone inside or any other evidence does not allow us to distinguish whether the origin of this depression was due to pathology or was caused by another agent, e.g. carnivores. Other probable modifications present in the scapula should be studied in more detail in the future to elucidate their origin.

The right femur, CIAAP-111 (Fig. S4.11a), preserves a transversal deep V-shaped cut of about 30 millimeters in length in its anterior face, at the level of the distal diaphysis (Fig. S4.11b). It preserves on one of its edges what appears to be hertzian cones, and it is covered with the same patina coloration as the bone. No marks related to trampling have been observed on the specimen. In the articular surface of the tibia, CIAAP-109 (Figs. S4.11d, e), we have also identified at least 10 linear marks that could be associated with cut marks. Two of these marks are parallel to each other, and these appear to be the longest of the group, at just over 40 mm in length and with a typical V-shaped profile. The presence of these marks in the concavity of the articulation surface suggests that they cannot have a trampling origin, since evidence of the latter is also not observed in other areas of the bone. In addition, around tibial tuberosity, a large depression probably of a pathological nature, was observed, however, this assumption deserves a more detailed taphonomic study.

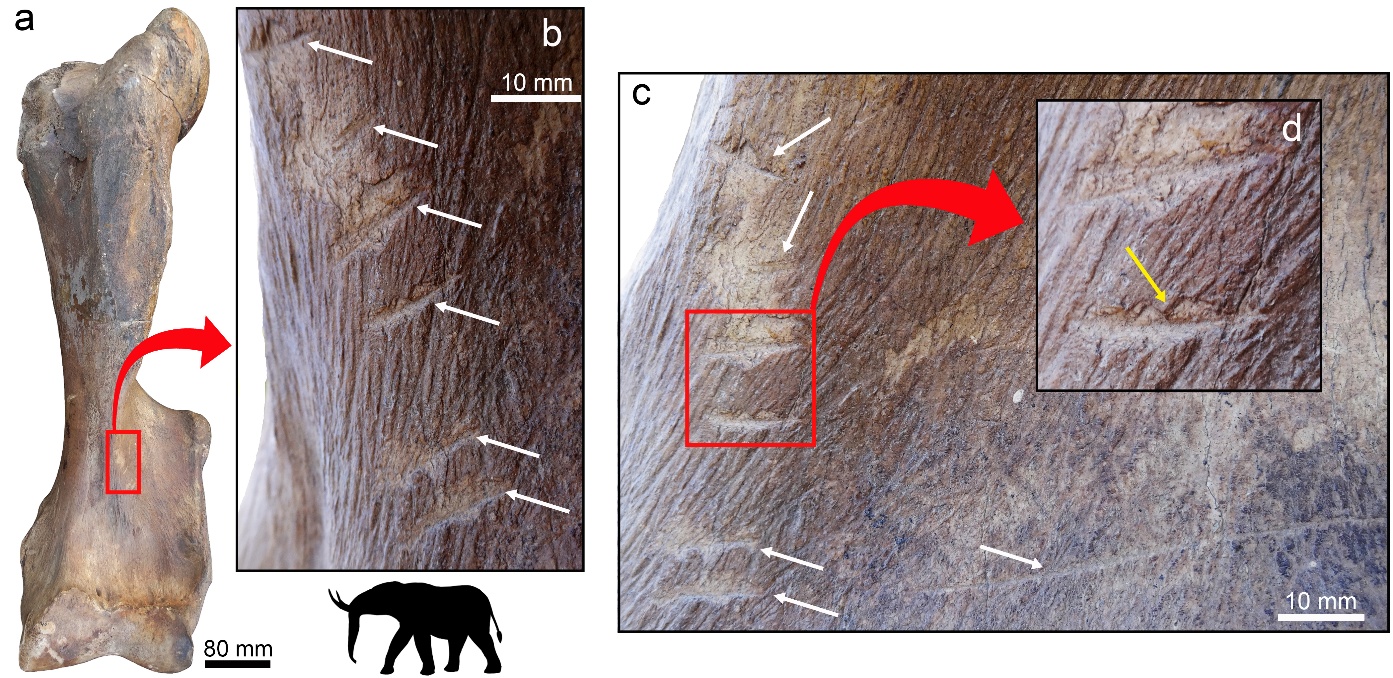

**Figure S4.10**. Left humerus (CIAAP-46) of *Notiomastodon platensis* in anterior view (a) with cut marks on anterior surface above the base of the humeral crest (b–d). This specimen belongs to the semi-articulated adult individual collected during the 1976 excavation. Marks are indicated by white arrows. Yellow arrow (d) exemplifies a hertzian cone in one of the cut marks.

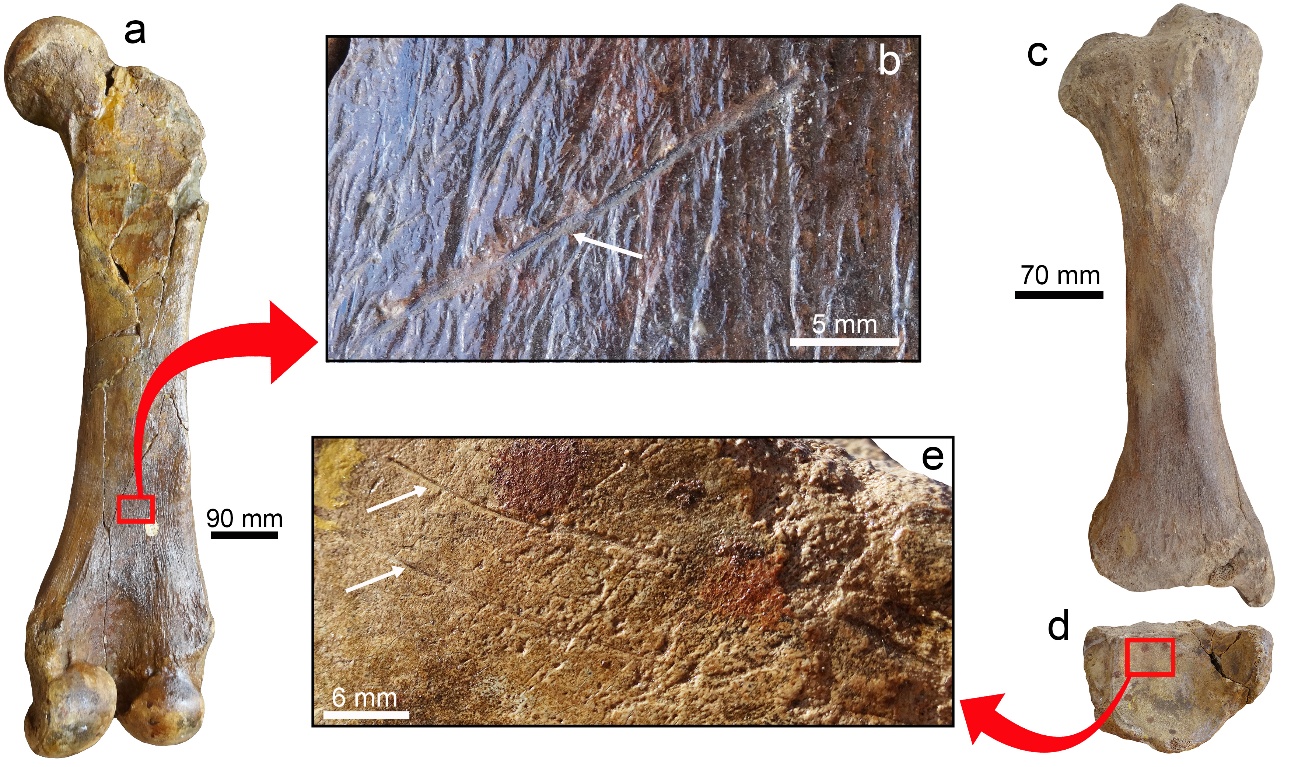

**Figure S4.11**. Right femur (CIAAP-111) and right tibia (CIAAP-109-211) of *Notiomastodon platensis* in posterior (a), anterior (c) and distal (d) views, showing marks (b, e). These specimens belong to the semi-articulated adult individual collected during the 1976 excavation. Marks are indicated by withe arrows.

Other 4 specimens of *N*. *platensis* reported here (Table 1, electronic supplementary material S2) as isolated elements and likely to belong to individuals other than the 1976 skeleton, preserve clear evidence of probable modifications of human nature. These specimens are represented by two right scapulae (CIAAP-119-110-210 and CIAAP-213-230) and a left ilium (CIAAP-19-233).

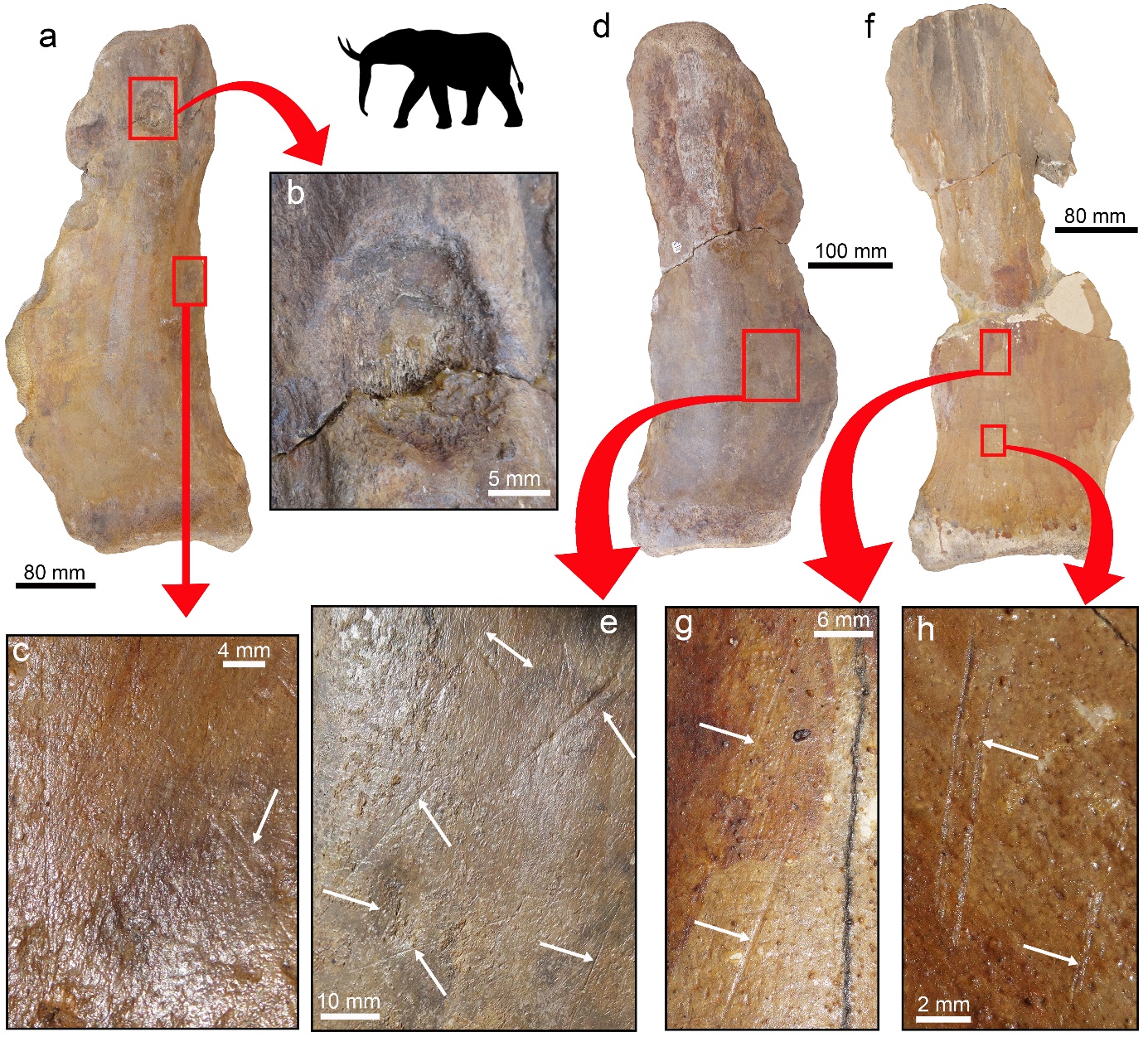

**Figure S4.12**. Left (a: CIAAP-47-229), and right (d: CIAAP-213-230; f: CIAAP-119-110-210) scapulae of *Notiomastodon platensis* from Taima-Taima, showing different linear cut marks (c, e, g, h) and possible pathology (b). White arrows show the marks. The specimen CIAAP-47-229 belongs to the semi-articulated adult individual collected during the 1976 excavation.

Of the two right scapulae, one (CIAAP-119-110-210) preserve two groups of three to four deep and slicing marks below the rim of the glenoid cavity in the anterior side (Fig. S4.12f–h). The most representative and best preserved are three marks, parallel to each other, with a maximum length of about 15 mm, being deep with a V-shaped section (Fig. S4.12h). In the other right scapula CIAAP-213-230 (Fig. S4.12d, e), we have documented what appears to be a depression, probably originating by percussion, and many linear marks, which could have different textures. We do not rule that many of these linear marks could be related to erosive processes (e.g., trampling). However, some of them are deep, with V-shaped profiles and we believe that they were produced during the dismemberment process. CIAAP-213-230, like many other specimens studied here, and others that were not documented from Taima-Taima, should be subjected to more detailed taphonomic studies in the future. One of the problems when trying to observe the details in specimens such as the two scapulae mentioned here, is that these were formerly prepared and consolidated with varnishes that generated a thick layer, which somehow covers delicate details of the marks, preventing even their observation during the photographic record.

The left ilium (CIAAP-19-233) is incomplete but preserves the acetabulum (Figs. S4.13 to S4.14). This ilium was mentioned by Casamiquela (1979) as n° 19, referring to it as a larger ilium than those preserved in the pelvic elements of the 1976 skeleton. This ilium is one of the most complex bone elements that we have documented in the sample studied, due to the large quantity and variety of modifications that are preserved. One side of the ilium is more affected than the other, and the coloration between the two sides is different, with the most affected side preserved in a dark red-brown color (Fig. S4.14). The most affected side stands out for the large number of depressions with loss of cortical material that seem to have been produced likely by impacts (Fig. S4.13d, e). Some fissures and cracks are observed on this side, and they seem to have originated during the percussion process to which this flat bone was subjected. Clearly, what appears to be chop marks can be observed deep and with V-shaped profiles. The longest (e.g., Fig. S4.15) of these appears to be about 120 mm long, and on the edges of one of its ends preserves what appears to be hertzian cones. Other smaller marks have similar patterns as described above. On the side that we call “less affected”, the evidence of modifications is also no less significant, and some areas of this side appear to be scraping marks (Fig. S4.16). The latter are very elongated which does not match similar patterns produced by trampling (see Fernández-Jalvo and Andrews, 2016). If these marks were indeed made by humans, they must have been produced by holding the stone tool edge transversely to the direction of the motion. At least two of the fissures/cracks observed on this side of the ilium appear to have been produced before the ilium was buried.

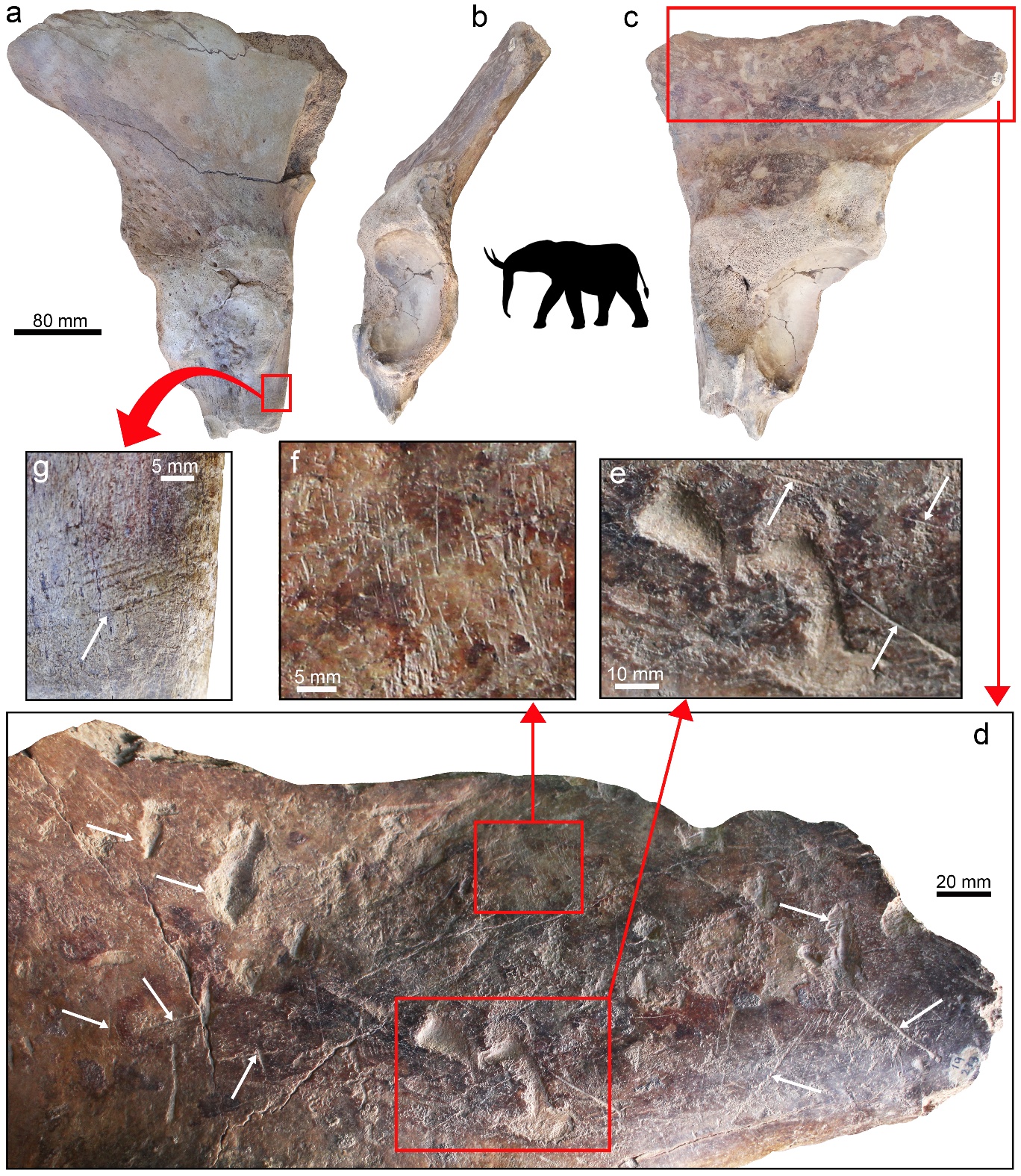

**Figure S4.13**. Left ilium fragment (a–d: CIAAP-19-233) of *Notiomastodon platensis* from Taima-Taima preserving a group of linear marks (d, some of them parallel: f and g) and depressions (e.g., d, e) with loss of cortical material produced by likely by percussion. White arrows show cutting marks and depressions.

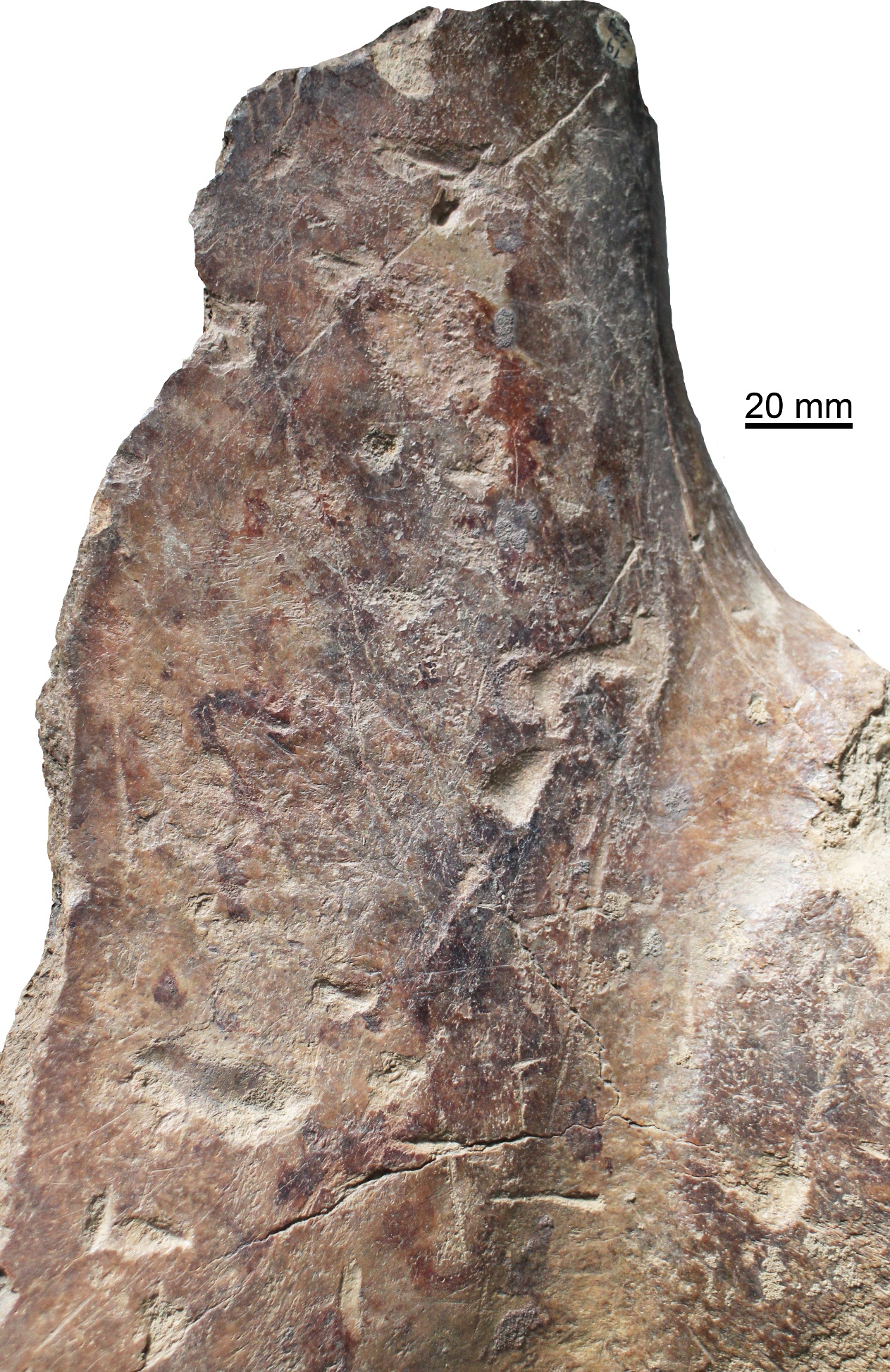

**Figure S4.14**. Left ilium fragment CIAAP-19-233 of *Notiomastodon platensis* from Taima-Taima preserving linear marks and depressions with loss of cortical material produced by likely by percussion.

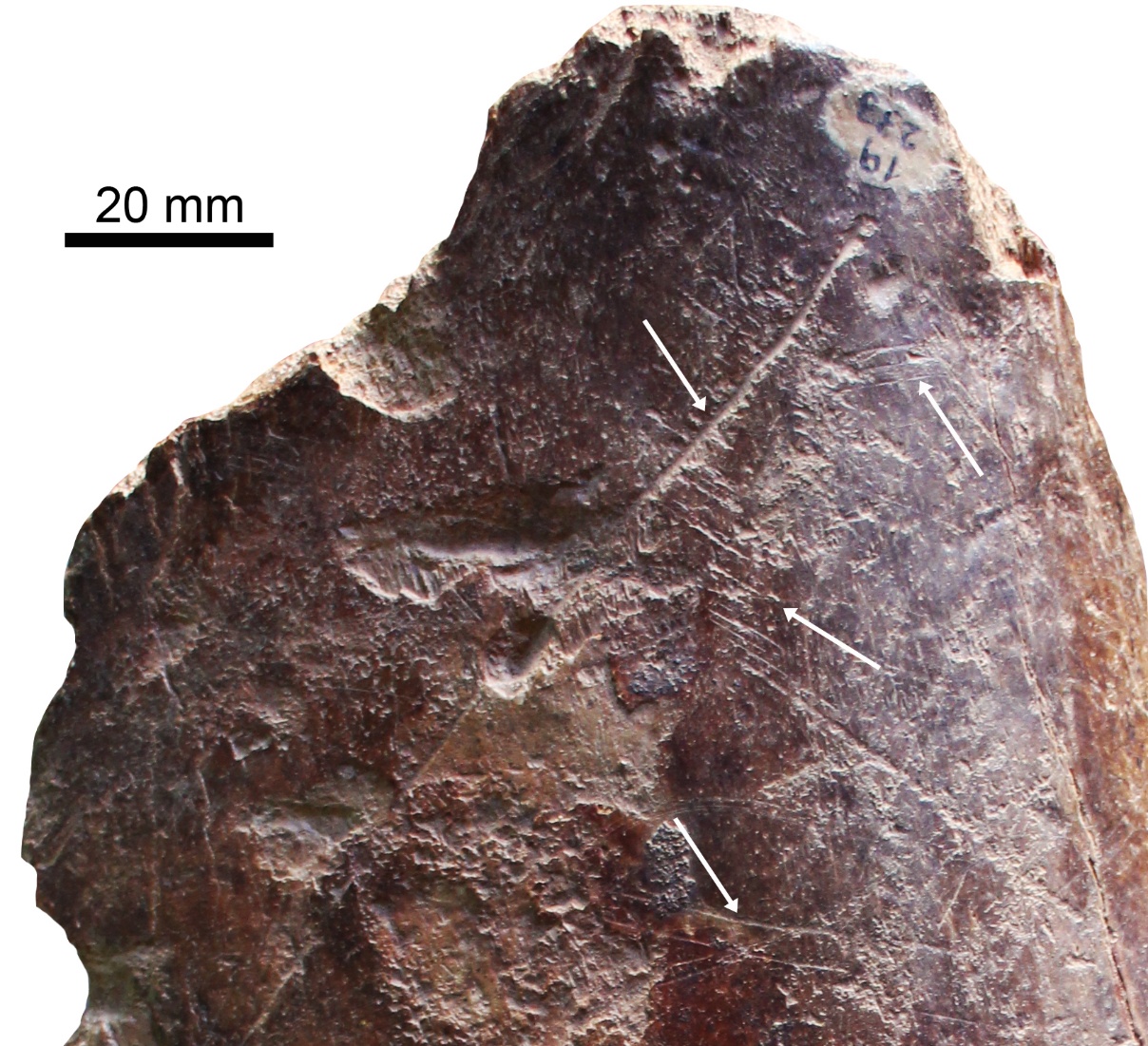

**Figure S4.15**. Left ilium fragment CIAAP-19-233 of *Notiomastodon platensis* from Taima-Taima preserving linear marks and depressions with loss of cortical material. White and red show cutting marks.

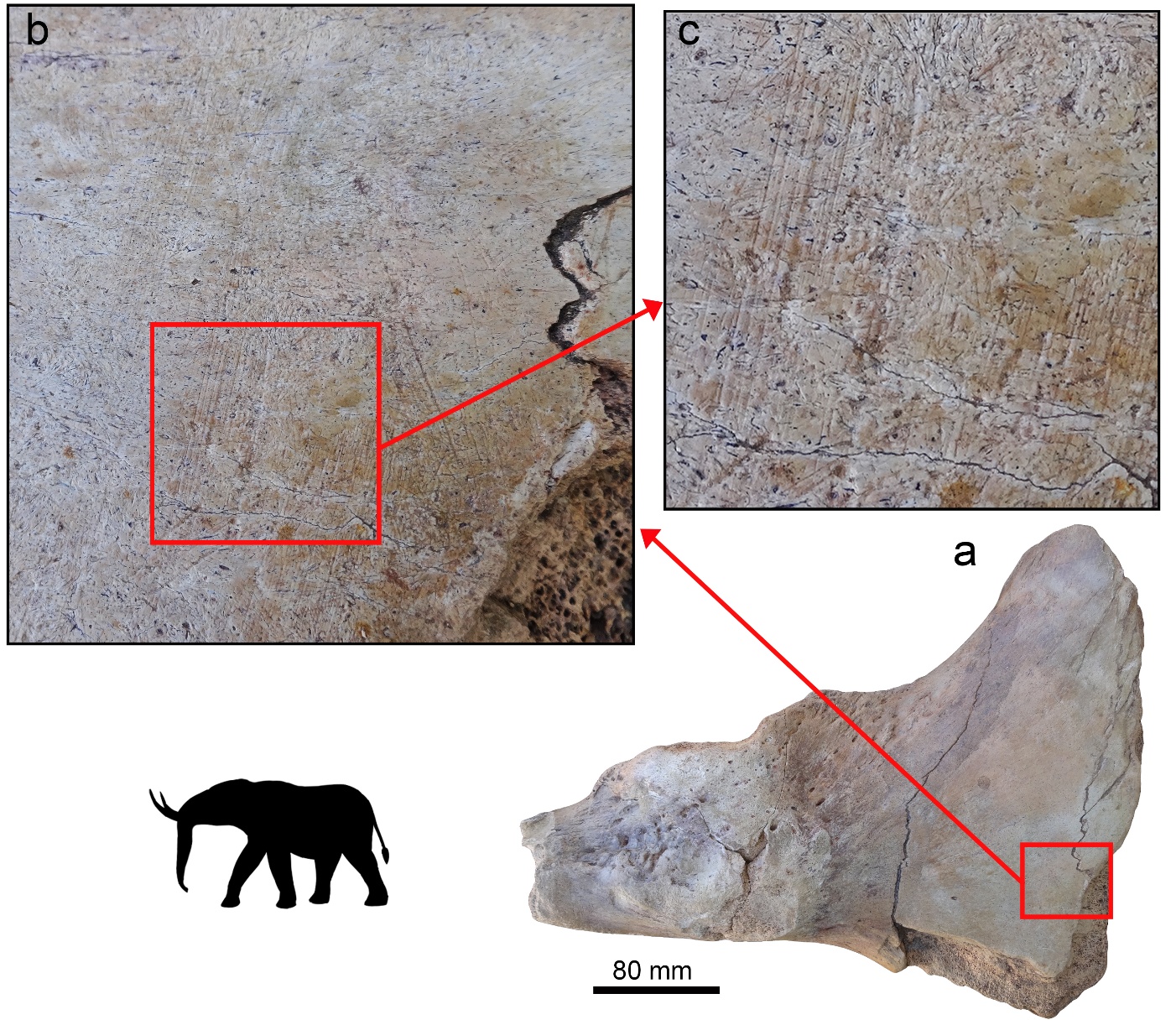

**Figure S4.16**. Left ilium fragment (a: CIAAP-19-233) of *Notiomastodon platensis* from Taima-Taima preserving group of scraping marks (b, c).

2. Further evidence of possible modification of human origin in gomphotheres bones

In the CIAAP collection in Coro, there are some *N*. *platensis* bones characterized by modifications and other alterations that could be of anthropogenic origin. However, although we mention and offer a brief description of the specimens here, they should be studied in more detail in the future to confirm that assumption. These specimens are represented by the two femora CIAAP-58-307 and CIAAP-90. These femora have something in common, and it is the large number of modifications that they preserve on their surface, especially those associated with probable percussion impacts and loss of cortical material (Fig. S4.18). These femora are incomplete, each missing one of the epiphyses. Both bones come from the Basal stratum (Cobble pavement), where a large part of the bones found in the layer are also in an incomplete state, especially the long ones (e.g., other femora) where the epiphyses are missing. Casamiquela (1979) presented a detailed discussion of the interpretation of fractures of intentional origin in long bones from this layer. The depressions observed in the cortical area of CIAAP-58-307, and CIAAP-90 (Fig. S4.18e, f, i, j) have a pattern like the percussion marks observed in the ilium (CIAAP-19-233 (Fig. S4.14). Some extreme cases of modification are observed, such as in CIAAP-90, the most modified bone so far found at the Taima-Taima site. These modifications probably have an anthropic origin and not a natural/taphonomic one (see Casamiquela, 1979). Casamiquela (1979: 71–72 pp.), suggested that femora CIAAP-58-307 and CIAAP-90 were likely used as anvils, possibly for chopping meat. CIAAP-90 has a rounded depression in its distal part, even some marked lines that could be more associated with cut marks than trampling (Fig. S4.18c). Some of these depressions observed in both femora allow us to more clearly appreciate the loss of cortical material. But most of the depressions appear to be altered or were worn out by an erosive agent that may have modified their surface and edges, giving the impression of having a smoother surface. Possibly these femurs were somewhat more exposed to erosive agents at the site than the ilium described above. We believe that many shallow and short linear marks also preserved in both bones could have a non-anthropic origin (e.g., trampling). A taphonomic study of how the rocks found in the basal stratum and the site dynamics may have affected these two femora is essential for future analysis and interpretation.

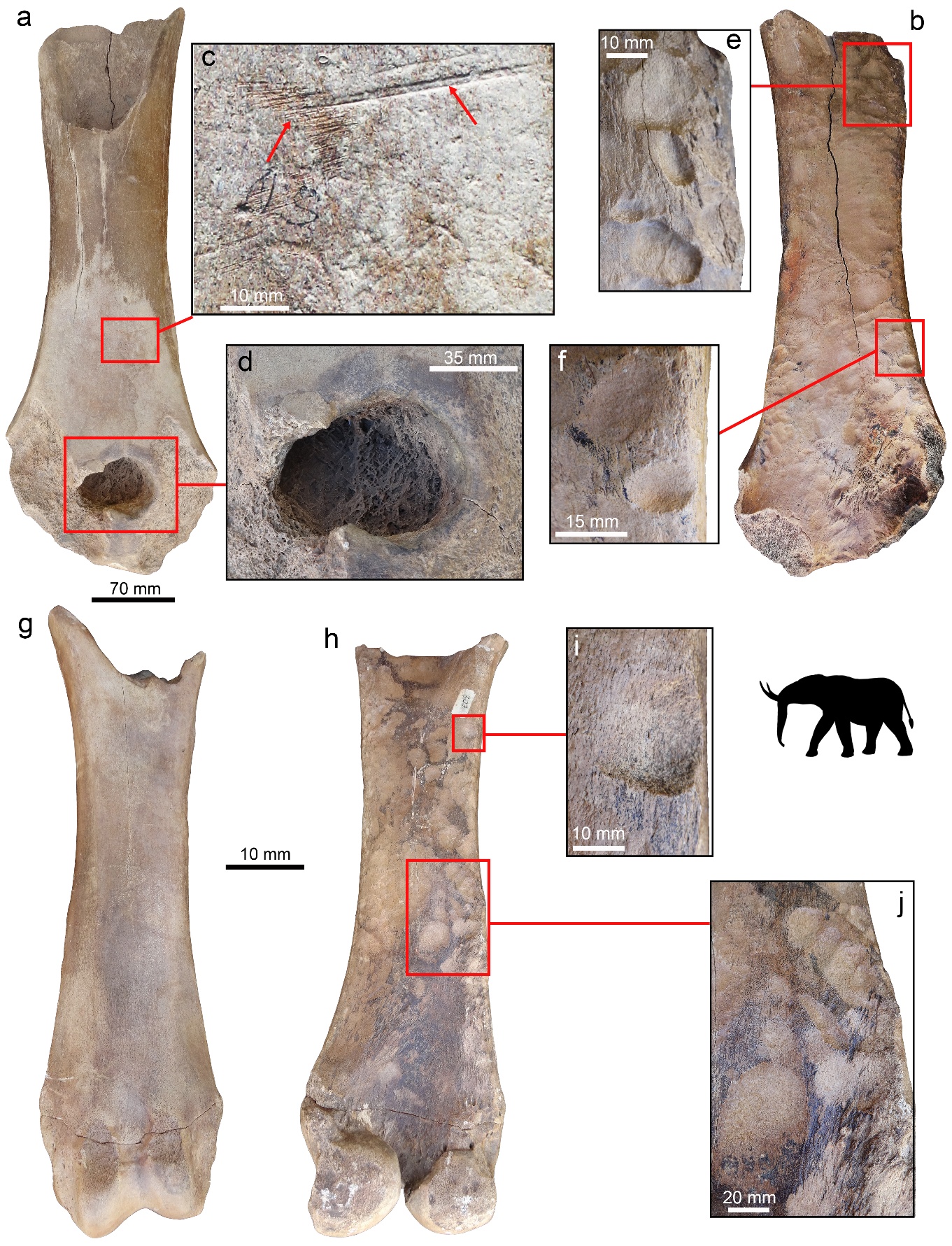

**Figure S4.18**. Right (a, b: CIAAP-90) and left (g, h: CIAAP-58-307) femora of *Notiomastodon platensis* from Taima-Taima preserving probably cutting marks (e.g., c), depressions and loss of cortical bone (e, f, i, j) and a hole with smooth edges (d). Femora CIAAP-90 and -58-307 were interpreted as being used as anvils (see Ochsenius and Gruhn, 1979).

3. Number of individuals exploited at the Taima-Taima site.

An exact determination of the number of *N*. *platensis* individuals that were exploited at the Taima-Taima site, and based only on the specimens studied here, is difficult to determine. The latter is due to the lack of association between most of the bone elements that have been reported as isolated in different excavation campaigns. For example, if we take as reference the pelvic elements and the right scapula of the 1976 skeleton, together with the other two pelvic elements (left ilium CIAAP-19-233, and complete pelvic bone from the 1974 excavation, Fig. S1.4a), or the other two right scapulas (CIAAP-119-110-210 and CIAAP-213-230; electronic supplementary material S2) of two other individuals, all of them with evidence of anthropogenic modifications, we would be referring to at least three individuals. This is a conservative approximation, especially considering the number of *N*. *platensis* individuals we have totaled for Taima-Taima is around 14 individuals (electronic supplementary material S1).
