## Supplementary material for "Diverse megamammals exploited by humans, chronology and palaeoecology at Taima-Taima, Late Pleistocene, South America": Electronic supplementary material S6

**Phytolith and starch grain analysis of dental calculus from the Taima-Taima site.**

**1. Specimens analyzed**

The analysis was carried out based on 8 dental calculus samples collected from molars of the proboscidean *Notiomastodon platensis* from the Late Pleistocene site of Taima-Taima, Falcón State, northwestern Venezuela (Table S6.1). The studied specimens are housed in the Centro de Investigaciones Antropológicas, Arqueológicas y Paleontológicas (CIAAP) of the Universidad Experimental Francisco de Miranda (UNEFM), Falcón State, and Laboratorio de Arqueología del Instituto Venezolano de Investigaciones Científicas (IVIC) in Miranda State, both in Venezuela.

Dental calculus was removed using dental tools and placed in airtight plastic containers to prevent contamination. Subsequently, the extraction process was carried out in the Laboratório de Microarqueologia – LABMICRO of the Museu de Arqueologia e Etnologia of the Universidade de São Paulo, Brazil, between July 29 and August 2, 2025. Followed standard procedures, with the hydrogen peroxide (H₂O₂) phase omitted to prevent potential damage to or digestion of the starch grains. All 8 samples exhibited little to no reaction to hydrochloric acid (HCl). Notably, the decalcification of sample IVIC-AP-032 (Table S6.1) was slower, taking approximately 10 hours, while the other samples were completed within 6 to 7 hours. The acid reaction in all cases was minimal, producing only light bubbling and releasing colourless gas. The phytoliths extracted from the samples showed significant taphonomic degradation Two slides were prepared from each sample.

**Table S6.1**. Dental pieces from the Taima-Taima site in which dental calculus was sampled. Abbreviations: dp, deciduous premolar; m/M, molar.

| **Catalog N°** | **Samplend element** | **Taxonomy** |
| --- | --- | --- |
| CIAAP-67 | Left lower dp4, from mandible | *Notiomastodon platensis* |
| CIAAP-1479 | Isolated lower right m2 | *Notiomastodon platensis* |
| CIAAP-1481 | Isolated upper left M2 | *Notiomastodon platensis* |
| CIAAP-1485 | Isolated lower right m3 | *Notiomastodon platensis* |
| IVIC-AP-030 | Lower right m2 from Mandible | *Notiomastodon platensis* |
| IVIC-AP-031 | Isolated lower right m3 | *Notiomastodon platensis* |
| IVIC-AP-032 | Isolated lower left m2 | *Notiomastodon platensis* |
| IVIC-AP-034 | Isolated upper left M2 | *Notiomastodon platensis* |

**2. Methods**

Sample extraction was conducted using an adjusted version of the protocol proposed by Santiago-Marrero and Pagán-Jiménez (2023). Phytoliths and starch grains were identified, counted, and photographed with a Leica DM500 light microscope at 200× and 400× magnification, following published reference material (e.g. Twiss et al., 1969; Brown, 1984; Piperno and Pearsall, 1998; Torrence et al., 2004; Ezell et al., 2006; Piperno, 2006; Iriarte and Paz, 2009; Gismondi et al., 2018; Pearsall, 2018). A complete scan of the slides was conducted to maximize the result from each sample. The data obtained was plotted using C2 software (Juggins, 2010) and MATLAB Version: 9.13.0 (MathWorks, 2022). Phytolith nomenclature follows the International Code for Phytolith Nomenclature 2.0 (ICPN 2.0; Neumann et al., 2019).

The dental calculus extraction proceeded as follows:

2.1 Sample preparation.

Before starting the procedures, the work area was disinfected with a 5% bleach solution. Reusable tools were washed with bleach and subsequently sterilized in an autoclave. Standard laboratory safety practices were followed, including the use of masks and powder-free gloves. Sterile 15 ml tubes were prepared and clearly labelled. To monitor for potential contamination, blank controls (an air blank and a reagent blank) were included in each batch.

2.2 Sample washing.

After photographing each sample and assigning it a specific number (Fig. S6.1a), the samples were weighed (Fig. S6.1b) and transferred into properly sterilized 15ml test tubes (Fig. S6.1c, d). The tubes were then filled with a 5% sodium hexametaphosphate ((NaPO₃)₆) solution (Fig. S6.1e) and subjected to a 15-minute cycle in an ultrasonic bath at room temperature (Fig. S6.1f). At the end of this stage, the tubes were centrifuged three times at 2,500 rpm for 2 minutes each, with the supernatant discarded after every cycle and the pellet rinsed with distilled water.

2.3 Decalcification.

Approximately 5 ml of 10% hydrochloric acid (HCl) was added to each test tube (Fig. S6.1g). The tubes were then left at room temperature until bubbling ceased, (6–12 hours). During this time, the liquid changed colour from light yellow (Fig. S6.1h) to dark yellow. Afterward, the samples were centrifuged at 2,500 rpm for 2 minutes, and the acid was carefully removed. The residues were rinsed three times with distilled water to eliminate any remaining acid. Following a final centrifugation step, the supernatant was discarded, and the concentrated residue was transferred to a labelled tube (Fig. S6.1i).

2.4 Slide assembly.

Microscope slides and coverslips were autoclaved for 40 min and subsequently cleaned with 5% bleach, rinsed with distilled water, and washed with ethanol before drying. Inside an exhaust hood, one to two drops of a 50:50 glycerin–water solution were added directly to the slide together with the sample and mixed. Coverslips were then applied and sealed with resin.

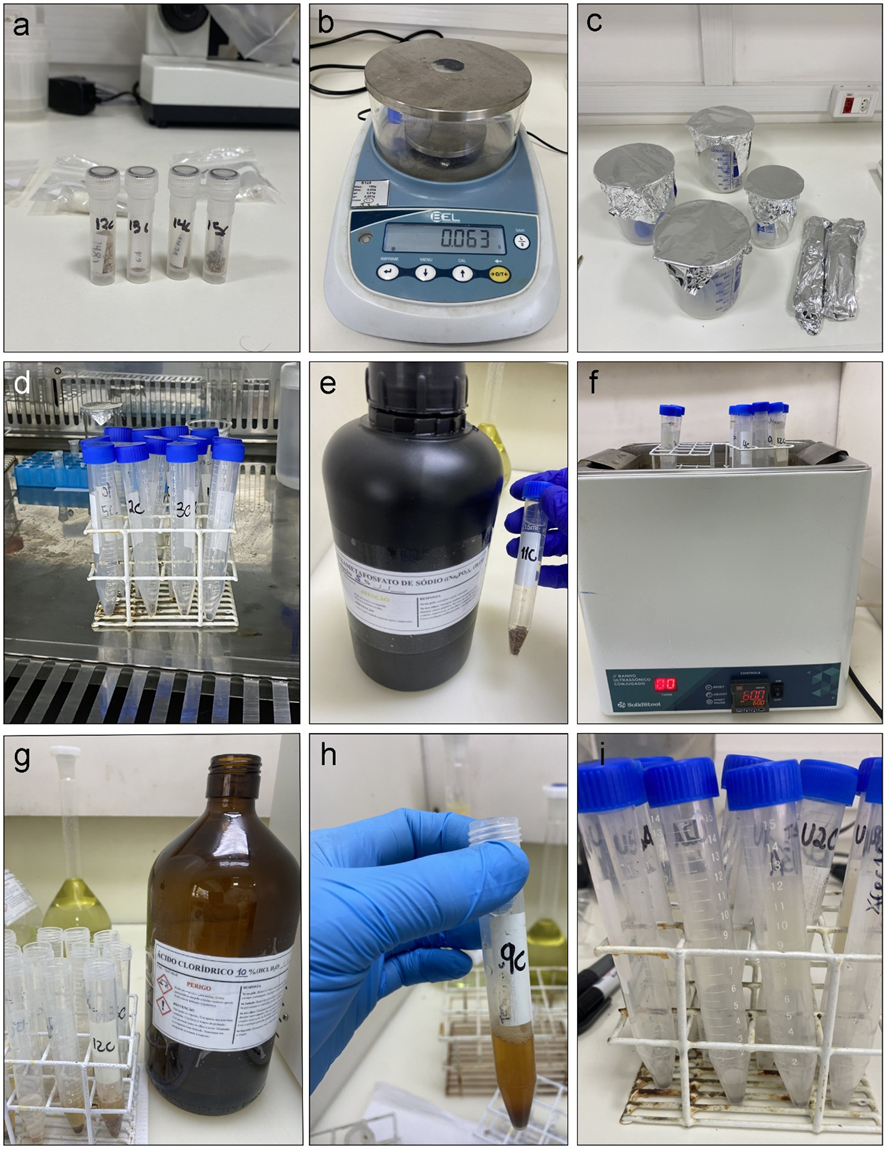

**Figure S6.1**. Steps in the dental calculus phytolith/starch grain extraction method. Photographing and assignment of numbers to samples (a); samples are weighed (b); tools are cleaned, wrapped in aluminium foil and sterilized in an autoclave (c); samples are added to 15ml test tubes (d); added 5% sodium hexametaphosphate ((NaPO₃)₆) solution to samples (e); ultrasonic bath phase (f); decalcification phase with 10% hydrochloric acid (HCl) (g); end of decalcification phase (h); end of extraction (i).

**3. Results**

A total of 8 distinct samples were analysed, resulting in the identification of 19 phytoliths and 8 starch grains. Results are presented by site in the following subsections, beginning with the findings from Taima-Taima, allowing for a detailed examination of the microbotanical evidence recovered from each location.

Phytoliths were identified from five plant taxa, listed alphabetically: arboreal dicotyledons, Arecaceae, Asteraceae, Marantaceae and Poaceae. Starch grains were associated with three taxa: Dioscoreaceae, Fabaceae, and Poales.

The main taxa identified in the samples include non-diagnostic arboreal dicotyledons, represented by SPHEROID GRANULATES (Fig. S6.2a) and SCLEREIDS (Fig. S6.2b). Poaceae (grasses) morphotypes were classified as BULLIFORMS (Fig. S6.2c), and sample CIAAP-1481 (Table S6.1) contained a broken lobate phytolith, likely belonging to the Panicoideae subfamily (Fig. S6.2d). Marantaceae (arrowroot family, which includes rhizomatous herbs) phytoliths were exclusively conical to TABULAR VERRUCATE forms (Fig. S6.2e, f), typically produced in roots and rhizomes. Perforated opaque bodies from the Asteraceae (e.g., sunflower and daisy family) taxa (Fig. S6.2g) were present in only one sample (IVIC-AP-030), corresponding to morphotypes typically produced in the seeds or inflorescences. Arecaceae (palms) SPHEROID SPINULOSE phytoliths (Fig. S6.2h) were also identified in a single sample (IVIC-AP-034); while this morphotype is produced in all parts of the plant, given the dental calculus context, it is likely derived from a palm fruit or seed. Full microremain data for all samples are presented in Table S6.2.

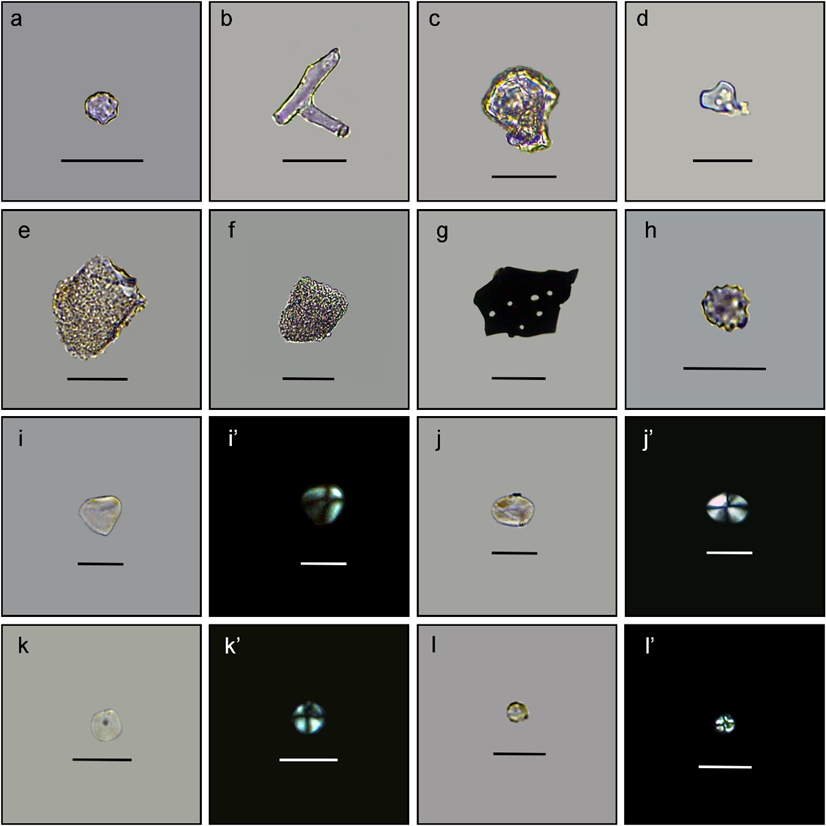

**Figure S6.2**. Phytoliths and starch grains recovered from the Taima-Taima site samples. SPHEROID GRANULATE produced in all parts of Arboreal dicotyledons (CIAAP-1485) (a). SCLEREID produced in all parts of Arboreal dicotyledons (IVIC-AP-032) (b). BULLIFORM FLABELLATE produced in the leaf/stem/inflorescence of Poaceae (IVIC-AP-030) (c). Damaged/broken likely BILOBATE produced in the leaf/stem of Panicoideae (Poaceae) (CIAAP-1481) (d). CONICAL VERRUCATE produced in the roots/rhizome of Marantaceae (e: CIAAP-1481, and f: IVIC-AP-031). Opaque PERFORATED-SHEET produced in the inflorescence of Asteraceae (IVIC-AP-030) (g). SPHEROID SPINULOSE produced in all parts of Arecaceae (IVIC-AP-034) (h). Broadly triangular with rounded edges starch grain likely from cf. Dioscoreaceae (CIAAP-1481) (i: brightfield, and i’: polarized). Rounded–ovoid starch grain likely from cf. Fabaceae (CIAAP-1479) (j: brightfield, j’: polarized). Ovoid -angular starch grain likely from cf. Poales (k, k’: IVIC-AP-031, and l, l’: IVIC-AP-032) (k, l: brightfield, k’, l’ polarized). Scales=20 µm.

The most common starch grains identified in the samples likely belonged to the order Poales (Fig. S6.2k, l), which includes grasses and sedges. These starch grains were recovered from five of the eight samples analysed (IVIC-AP-031, IVIC-AP-032, CIAAP-1481, -67, and -1485). The starch grains were generally small to medium in size, predominantly spherical to polyhedral, and featured mostly centric hila. In some cases, they showed faint extinction crosses, characteristics consistent with those of grasses. Two samples (IVIC-AP-030 and CIAAp-1479) contained starch grains that can likely be attributed to the Fabaceae family, which includes beans and other legumes. These starch grains were larger (~20 µm), ovoid to rounded in shape, and sometimes exhibited fissures around the hilum.

Finally, sample CIAAP-1481 showed starch grains consistent with Dioscoreaceae, the yam family, which is known for its starchy underground tubers. The presence of these three groups indicates a dietary contribution from grasses, legumes, and tuber-producing plants.

Overall, the results from Taima-Taima reveal evidence of diverse plant consumption by proboscideans (Fig. S6.3 and Fig. S6.4). Phytoliths and starch grains were recovered from grasses (Poaceae), arboreal dicotyledons, palms (Arecaceae), Asteraceae, and Marantaceae rhizomes, as well as starches tentatively identified as Fabaceae and Dioscoreaceae. As no sedges or other Poales phytoliths were identified, it is likely that Poales starches derive from Poaceae plants. This indicates that grasses, in addition to arboreal dicotyledons, are the main microbotanical remains found in the Taima-Taima proboscidean dental calculus.

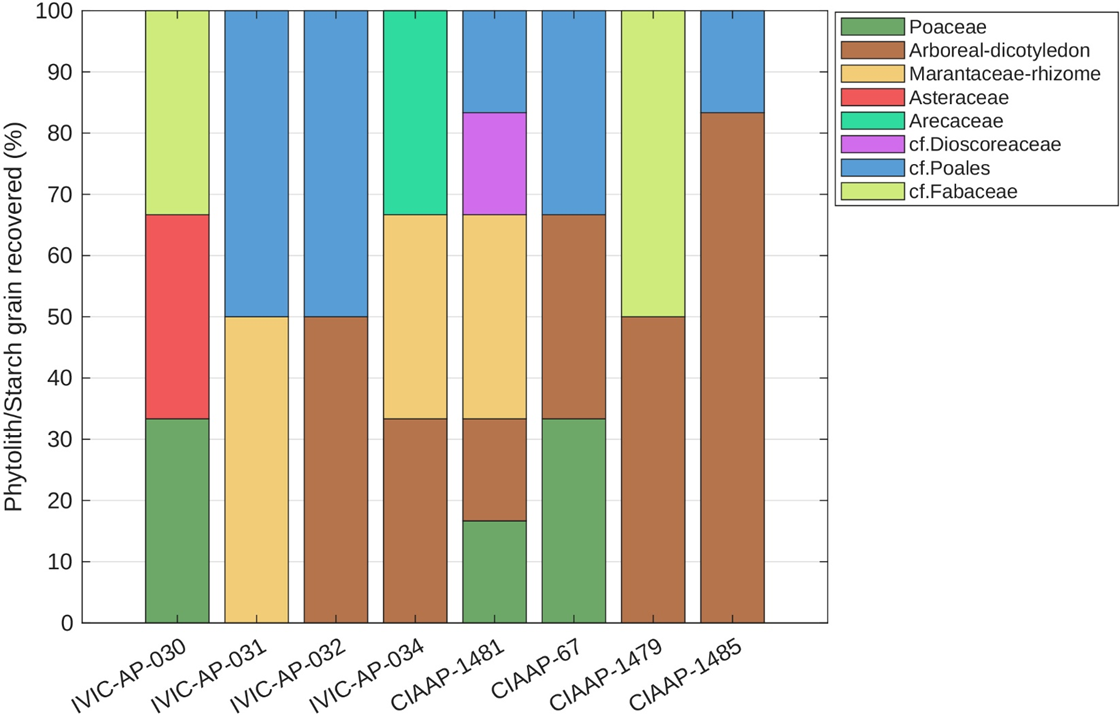

**Figure S6.3**. Stacked bar chart of phytoliths and starch grains recovered from the Taima-Taima samples.

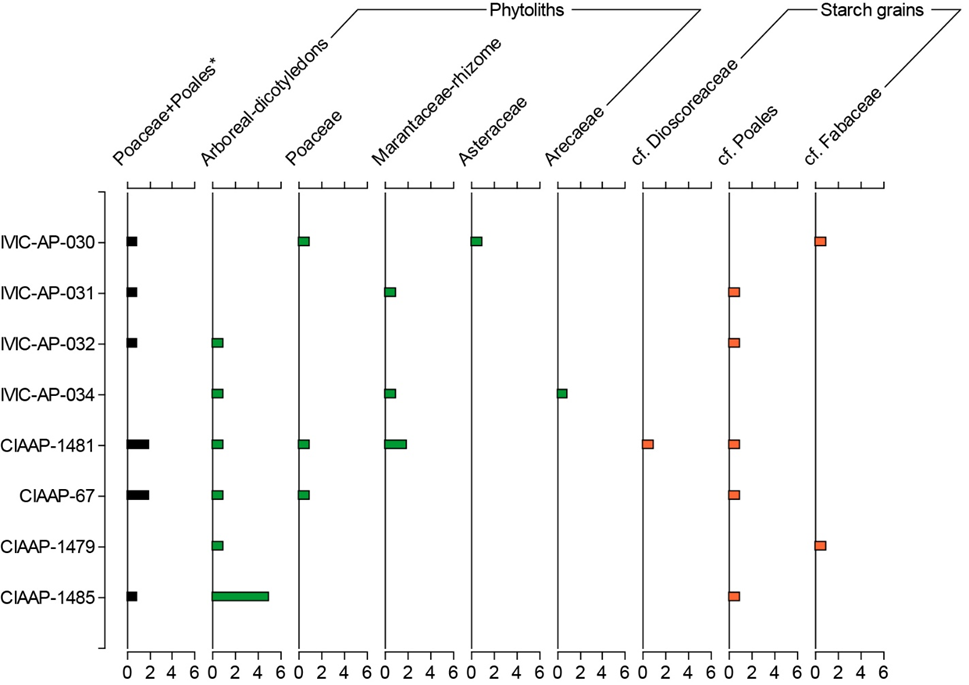

### **Figure S6.4**. Phytoliths and starch grains recovered from the Taima-Taima samples. Horizontal bars represent quantity of microremains recovered. * Poaceae + Poales category combines all grass and grass-like starches with securely identified Poaceae phytoliths to provide a likely complete representation of grasses in the assemblage. Colours are used only for visual differentiation: green = phytoliths, orange = starch grains, and black = total Poales (all phytoliths and starches identified).

**Table S6.2**. Microremains identified in the Taima-Taima samples.

| **Catalog N°** | **Poaceae** | **Arboreal-dicotyledons** | **Marantaceae rhizome** | **Asteraceae** | **Arecaeae** | **cf. Dioscoreaceae** | **cf. Poales** | **cf. Fabaceae** |
| --- | --- | --- | --- | --- | --- | --- | --- | --- |
| IVIC-AP-030 | 1 | 0 | 0 | 1 | 0 | 0 | 0 | 1 |
| IVIC-AP-031 | 0 | 0 | 1 | 0 | 0 | 0 | 1 | 0 |
| IVIC-AP-032 | 0 | 1 | 0 | 0 | 0 | 0 | 1 | 0 |
| IVIC-AP-034 | 0 | 1 | 1 | 0 | 1 | 0 | 0 | 0 |
| CIAAP-1481 | 1 | 1 | 2 | 0 | 0 | 1 | 1 | 0 |
| CIAAP-67 | 1 | 1 | 0 | 0 | 0 | 0 | 1 | 0 |
| CIAAP-1479 | 0 | 1 | 0 | 0 | 0 | 0 | 0 | 1 |
| CIAAP-1485 | 0 | 5 | 0 | 0 | 0 | 0 | 1 | 0 |
