## Supplementary material for "Diverse megamammals exploited by humans, chronology and palaeoecology at Taima-Taima, Late Pleistocene, South America": Electronic supplementary material S7

**Dietary reconstruction of herbivores from Taima-Taima**

Mesowear angles (MAs) for *Notiomastodon platensis* and *Eremotherium laurillardi* from Taima-Taima come from Wilson et al. (2024). The MA data for *Mixotoxodon larensis* and *Glossotherium tropicorum* from Muaco also comes from Wilson et al. (2024). For an estimate of diet in *Equus* sp*.*, *Palaeolama major* and *Xenorhinotherium bahiense* from Muaco, we used the classical mesowear approach of Fortelius and Solounias (2000), scoring the penultimate upper molar for both cusp shape and relief. Cusp shape was scored as either sharp (1), rounded (2) or blunt (3). Relief was coded as either high (1) or low (3), using selenodont/plagiolophodont threshold angles (Wilson and Saarinen, 2024). Average mesowear scores are an average of the cusp shape and relief scores for a given specimen (or across a whole species; Saarinen et al., 2016). While the numbers of specimens are low for each species, so we do not quantitatively reconstruct diet from these mesowear scores, in general, lower mesowear scores (sharp cusps and high relief) are consistent with a low abrasion browsing diet and higher mesowear scores (blunt cusps and low relief) are consistent with an abrasive, grazing diet.

For taxa where mesowear angles are used instead (e.g. proboscideans, xenarthrans and notoungulates; Saarinen et al, 2015; Saarinen and Karme, 2017; Wilson et al., 2024; Wilson and Saarinen, 2024), lower angles indicate a less abrasive diet, and higher angles a more abrasive diet, indicating grazing. For sloths, the threshold angle between browsing and mixed feeding is 100˚ and the corresponding mixed-feeding-grazing angle is 133˚ (Saarinen and Karme, 2017). For ungulates, threshold values between browsing and mixed feeding and between mixed feeding and grazing are 106˚ and 120˚ respectively (Saarinen and Lister, 2023; Wilson and Saarinen, 2024). In addition to the mesowear angles measured by Wilson et al. (2024), we measured additional mesowear angles from *Xenorhinotherium bahiense* from surface scans made using the Polycam 3D scanning application, following the procedure of Wilson et al. (2024). Angles in *Xenorhinotherium* were measured from the ectoloph wear facet and converted to mesowear angles, as has been done previously for notoungulates (Wilson et al., 2024; Wilson and Saarinen, 2024), following the equation: mesowear angle = 180 – 2×facet angle. These angles represent the first attempt to apply mesowear angles to lophodont litopterns, and the results from the angles match those from the classical mesowear, supporting their utility in future studies of litopterns. We could not obtain accurate estimates of mesowear angles from the scans of *Glyptotherium* cf. *cylindricum*, so rely on estimates from elsewhere in the continent for our dietary reconstruction in the main text.

It is possible that diet differs between Muaco and Taima-Taima, though they are located close together (separated by ~3 km). We consider that these values are likely to be more representative than reconstructions of palaeodiet of the same species from other localities elsewhere in South America, given that there can be significant differences in reconstructions across space (e.g. MacFadden, 2005; González-Guarda et al., 2025), though our final assessment of diet in all herbivores from Taima-Taima (Fig. 6) also include such data from elsewhere, given the relative paucity of dental material available so far. Future excavation and multiproxy studies will further clarify the diets of the megaherbivores of Taima-Taima.

The specimens from Muaco are housed in Museo Geológico Dr. José Royo y Gómez (UCV-VF), Caracas, Venezuela and the Museo Paleontológico de la Alcaldía del Municipio de Urumaco (AMU-CURS), Urumaco, Venezuela. The Taima-Taima specimens are housed in Centro de Investigaciones Antropológicas, Arqueológicas y Paleontológicas of the Universidad Experimental Francisco de Miranda (CIAAP-UNEFM), Coro, Venezuela.

**Table 1.** Classical mesowear scoring for *Equus* sp., *Palaeolama major* and *Xenorhinotherium bahiense* from Muaco, Falcón State, Venezuela.

| **Species** | **Specimen** | **Cusp shape** | **Relief** | **Mesowear score** |
| --- | --- | --- | --- | --- |
| *Equus* sp. | UCV-VF 449 | 3 | 3 | 3 |
| *Equus* sp. | UCV-VF 447 | 2 | 3 | 2.5 |
| *Equus* sp. | UCV-VF 474 | 1 | 3 | 2 |
| ***Equus* sp. *s*pecies mean** | | **2** | **3** | **2.5** |
| *Palaeolama major* | UCV-VF 393 | 1 | 1 | 1 |
| *Palaeolama major* | UCV-VF 436 | 1 | 1 | 1 |
| ***Palaeolama major* species mean** | | **1** | **1** | **1** |
| *Xenorhinotherium bahiense* | UCV-VF 354 | 1 | 1 | 1 |
| *Xenorhinotherium bahiense* | UCV-VF 352 | 1 | 2 | 1.5 |
| *Xenorhinotherium bahiense* | UCV-VF 361 | 1 | 1 | 1 |
| ***Xenorhinotherium bahiense* species mean** | | **1** | **1.33** | **1.17** |

**Table 2.** Mesowear angles for herbivores from Taima-Taima and Muaco, Falcón State, Venezuela.

| **Locality** | **Order** | **Species** | **Specimen** | **Mesowear angle, ˚** | **Source** |
| --- | --- | --- | --- | --- | --- |
| Taima-Taima | Proboscidea | *Notiomastodon platensis* | CIAAP-UNEFM 1483 | 125 | Wilson et al., 2024 |
| Taima-Taima | Proboscidea | *Notiomastodon platensis* | CIAAP-UNEFM 1479 | 134 | Wilson et al., 2024 |
| Taima-Taima | Proboscidea | *Notiomastodon platensis* | CIAAP-UNEFM 224 | 135 | Wilson et al., 2024 |
| Taima-Taima | Proboscidea | *Notiomastodon platensis* | CIAAP-UNEFM 214 | 129 | Wilson et al., 2024 |
| Taima-Taima | Proboscidea | *Notiomastodon platensis* | CIAAP-UNEFM 80 | 119 | Wilson et al., 2024 |
| **Taima-Taima *Notiomastodon platensis* species mean** | | | | **128.40** | |
| Muaco | Proboscidea | *Notiomastodon platensis* | UCV-VF 117 | 148 | Wilson et al., 2024 |
| Muaco | Proboscidea | *Notiomastodon platensis* | UCV-VF 112 | 131 | Wilson et al., 2024 |
| Muaco | Proboscidea | *Notiomastodon platensis* | UCV-VF 263 | 133 | Wilson et al., 2024 |
| Muaco | Proboscidea | *Notiomastodon platensis* | UCV-VF 262 | 130 | Wilson et al., 2024 |
| **Muaco *Notiomastodon platensis* species mean** | | | | **135.50** | |
| **Regional *Notiomastodon platensis* species mean** | | | | **131.56** | |
| Taima-Taima | Pilosa | *Eremotherium laurillardi* | CIAAP-UNEFM-596 | 103 | Wilson et al., 2024 |
| Taima-Taima | Pilosa | *Eremotherium laurillardi* | CIAAP-UNEFM-597 | 110 | Wilson et al., 2024 |
| **Taima-Taima *Eremotherium laurillardi* mean** | | | | **106.5** | |
| Muaco | Pilosa | *Eremotherium laurillardi* | UCV-VF-972 | 87 | Wilson et al., 2024 |
| Muaco | Pilosa | *Eremotherium laurillardi* | UCV-VF-177 | 92.5 | Wilson et al., 2024 |
| Muaco | Pilosa | *Eremotherium laurillardi* | UCV-VF-178 | 104.5 | Wilson et al., 2024 |
| Muaco | Pilosa | *Eremotherium laurillardi* | UCV-VF-175 | 104.5 | Wilson et al., 2024 |
| Muaco | Pilosa | *Eremotherium laurillardi* | UCV-VF-1178 | 92 | Wilson et al., 2024 |
| Muaco | Pilosa | *Eremotherium laurillardi* | UCV-VF-171 | 109 | Wilson et al., 2024 |
| **Muaco *Eremotherium laurillardi* species mean** | | | | **98.25** | |
| Quebrada Ocando | Pilosa | *Eremotherium laurillardi* | CIAAP-UNEFM-852 | 77 | Wilson et al., 2024 |
| **Regional *Eremotherium laurillardi* species mean** | | | | **97.72** | |
| Muaco | Pilosa | *Glossotherium tropicorum* | UCV-VF-205 | 152.5 | Wilson et al., 2024 |
| Muaco | Notoungulata | *Mixotoxodon larensis* | UCV-VF-391 | 101 | Wilson et al., 2024 |
| Muaco | Notoungulata | *Mixotoxodon larensis* | UCV-VF-381 | 80 | Wilson et al., 2024 |
| **Muaco *Mixotoxodon larensis* species mean** | | | | **90.5** | |
| Muaco | Litopterna | *Xenorhinotherium bahiense* | UCV-VF-354 | 80 | This study |
| Muaco | Litopterna | *Xenorhinotherium bahiense* | UCV-VF-353 | 101 | This study |
| Muaco | Litopterna | *Xenorhinotherium bahiense* | UCV-VF-352 | 103 | This study |
| Muaco | Litopterna | *Xenorhinotherium bahiense* | UCV-VF-361 | 88 | This study |
| **Muaco *Xenorhinotherium bahiense* species mean** | | | | **93** | |
